# Developing, characterizing and modeling CRISPR-based point-of-use pathogen diagnostics

**DOI:** 10.1101/2024.07.03.601853

**Authors:** Jaeyoung K. Jung, Kathleen S. Dreyer, Kate E. Dray, Joseph J. Muldoon, Jithin George, Sasha Shirman, Maria D. Cabezas, Anne E. D’Aquino, Matthew S. Verosloff, Kosuke Seki, Grant A. Rybnicky, Khalid K. Alam, Neda Bagheri, Michael C. Jewett, Joshua N. Leonard, Niall M. Mangan, Julius B. Lucks

**Author notes:** Center for Bio/Molecular Science and Engineering, US Naval Research Laboratory (Washington, D.C., USA).

## Abstract

Recent years have seen intense interest in the development of point-of-care nucleic acid diagnostic technologies to address the scaling limitations of laboratory-based approaches. Chief among these are combinations of isothermal amplification approaches with CRISPR-based detection and readouts of target products. Here, we contribute to the growing body of rapid, programmable point-of-care pathogen tests by developing and optimizing a one-pot NASBA-Cas13a nucleic acid detection assay. This test uses the isothermal amplification technique NASBA to amplify target viral nucleic acids, followed by Cas13a-based detection of amplified sequences. We first demonstrate an in-house formulation of NASBA that enables optimization of individual NASBA components. We then present design rules for NASBA primer sets and LbuCas13a guide RNAs for fast and sensitive detection of SARS-CoV-2 viral RNA fragments, resulting in 20 – 200 aM sensitivity without any specialized equipment. Finally, we explore the combination of high-throughput assay condition screening with mechanistic ordinary differential equation modeling of the reaction scheme to gain a deeper understanding of the NASBA-Cas13a system. This work presents a framework for developing a mechanistic understanding of reaction performance and optimization that uses both experiments and modeling, which we anticipate will be useful in developing future nucleic acid detection technologies.

## INTRODUCTION

The past several years have seen a surge of interest in developing point-of-care (POC) nucleic acid diagnostic technologies^1–5^. This was motivated by the SARS-CoV-2 pandemic and public health emergency, which highlighted the challenges of scaling laboratory-based testing capacity to scales necessary to monitor a global pandemic^6^. Although the current gold standard for pathogen testing, reverse transcription–polymerase chain reaction (RT-PCR), is sensitive and reliable, it necessitates technical expertise, centralized laboratory facilities and multiple reaction steps performed at different temperatures^7,8^. For these reasons, RT-PCR struggles to meet surges in demand, and is not suitable for accessible, cost-effective, and distributed point-of-care (POC) nucleic acid diagnostic technologies. As such, there is a recognized need for pathogen diagnostic tests that can be implemented outside of a laboratory setting and produce results with minimal human intervention, simple protocols, and reduced equipment This need has been widely recognized, resulting in POC pathogen tests that provide decreased time to readout and fewer reaction steps. These POC pathogen tests generally involve two steps: (1) isothermal amplification of specific pathogen nucleic acid sequences and (2) detection of the amplified sequences. Among isothermal amplification methods, loop-mediated isothermal amplification (LAMP)^9^ and recombinase polymerase amplification (RPA)^10^ have been used extensively in conjunction with detection techniques such as lateral flow assays^3,11^, colorimetric assays^12^, fluorescent readouts^3,11,13^ and next-generation sequencing^14,15^. Additionally, amplification-free detection methods such as clustered regularly interspaced short palindromic repeats (CRISPR)-Cas based detection^16^ and antigen-based tests that have been distributed for rapid screening^17–19^. Mobile-based devices and a “suitcase testing lab” also have been developed to improve portability and minimize subjective interpretation in analyzing test results^16,20,21^.

Here, we contribute to the growing body of POC tests by developing a diagnostic test that detects specific RNA sequences and produces a fluorescent readout. In contrast to prior work, we used nucleic acid sequence-based amplification (NASBA)^22^ to isothermally amplify a target RNA. NASBA uses three enzyme components – reverse transcriptase (RT), RNaseH, and T7 RNA polymerase (RNAP) – to amplify a target RNA based on supplied single stranded DNA (ssDNA) primers: reverse transcriptase (RT) and RNase H convert input single-stranded RNA (ssRNA) into T7 promoter-containing double-stranded DNA (dsDNA), which is transcribed by T7 RNA polymerase (RNAP) into an activator RNA. The activator RNA serves as an input to the cycle, promoting exponential amplification. The activator RNA is also detected by a CRISPR-Cas13a, an RNA-guided and RNA-activated ribonuclease^23,24^ that has been used in other nucleic acid detection strategies^1,25^. Upon recognition of the activator RNA by the designed Cas13a guide RNA (gRNA), Cas13a indiscriminately cleaves uracil residues in an ssRNA-based reporter to generate fluorescence^4^ (Figure 1a). This system can be reconfigured to detect a range of different RNAs simply by modifying the primers and gRNA. We chose NASBA because of its lower operating temperature (37∼41°C)^22,26^ compared to LAMP (60∼65°C)^9^, its low cost compared to RPA^26^, and its off-patent status, potentially allowing for rapid innovation and adoption, as well as distributed manufacturing of reaction components^27^. The lower operating temperature also could make it more amenable than other technologies to POC uses.

**Fig. 1.**
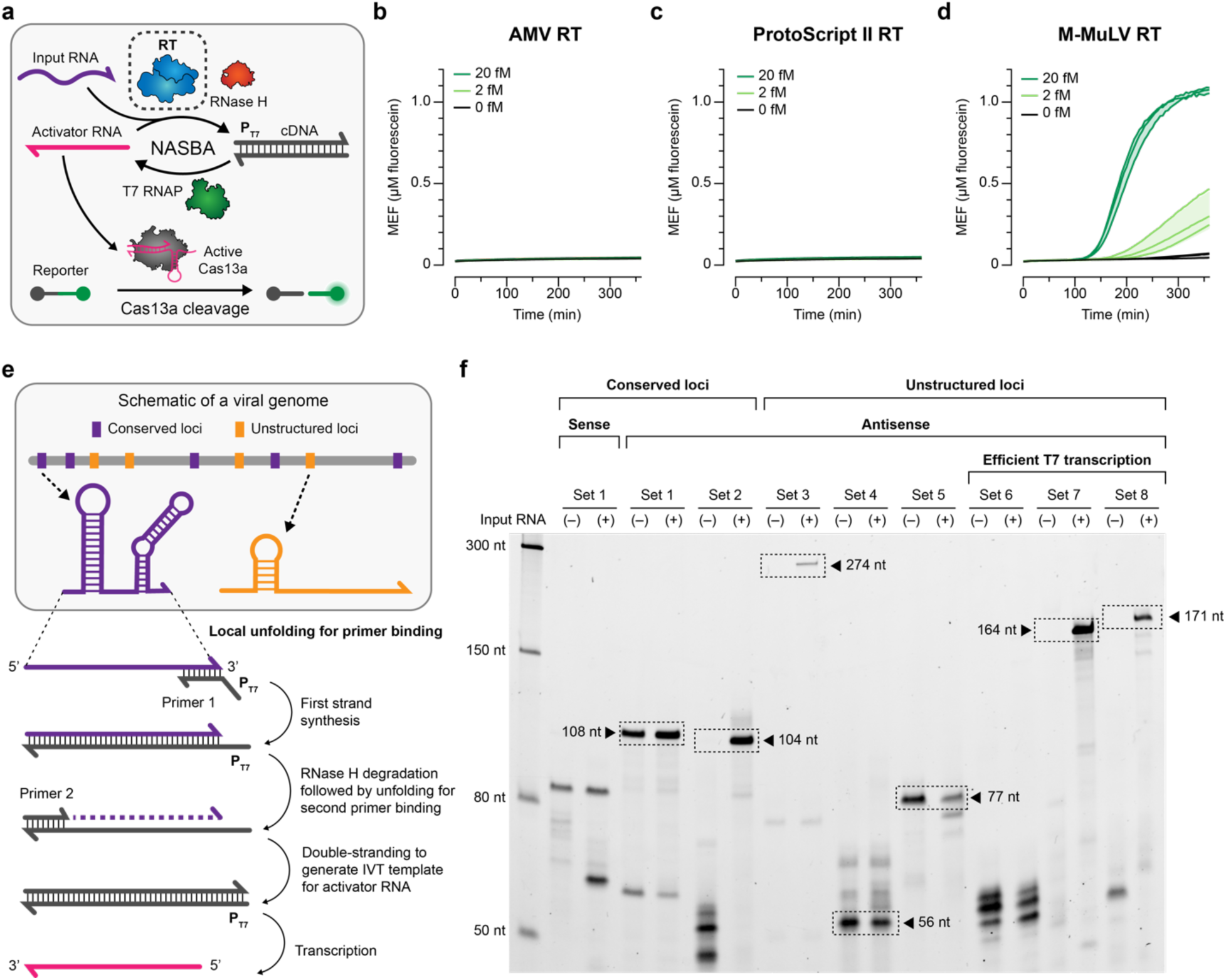
In-house NASBA formulation provides flexibility for reaction optimization. **a**, Schematic overview of NASBA, which uses cycles of reverse transcription, RNase H-mediated degradation and T7 transcription to convert and amplify an input RNA into an antisense activator RNA. The antisense activator serves as an input to Cas13a-based detection which generates a fluorescent output signal. In-house NASBA formulation enables screening of different reverse transcriptases (RTs). One-pot in-house NASBA-Cas13a targeting the ORF1ab of the SARS-CoV-2 genome, with 0, 2 or 20 fM synthetic SARS-CoV-2 genome and 1 U/µL of **b**, avian myeloblastosis virus (AMV) RT, **c**, ProtoScript II RT, or **d**, Moloney murine leukemia virus (M-MuLV) RT. Readout was observed only with M-MuLV RT. **e**, Schematic of the steps in NASBA with a cartoon of viral genome structures that could influence where NASBA primers bind and impact NASBA efficiency. **f**, To test different primer sets, RNA products were extracted from one-pot NASBA (lacking Cas13a) and analyzed by urea-PAGE. Reactions were initiated using 2.5 U/µL M-MuLV RT with 0 (-) or 20 (+) fM synthetic SARS-CoV-2 genome. The expected RNA product for each primer set is boxed and its length is indicated, unless the band was not present as in the case of sets 1 and 6 (expected products 104 nt and 164 nt, respectively). Data in **b**–**d** are n=3 independent biological replicates, each plotted as a line with raw fluorescence standardized to MEF. Shading in **b**–**d** indicates the average of the replicates ± standard deviation. Data in **f** are one representative of n=3 independent biological replicates; the other replicates and the uncropped, unprocessed image in **f** are in **Supplementary Data 2**. Sequences of primers and gRNAs are listed in **Supplementary Data 1**.

**Fig. 2.**
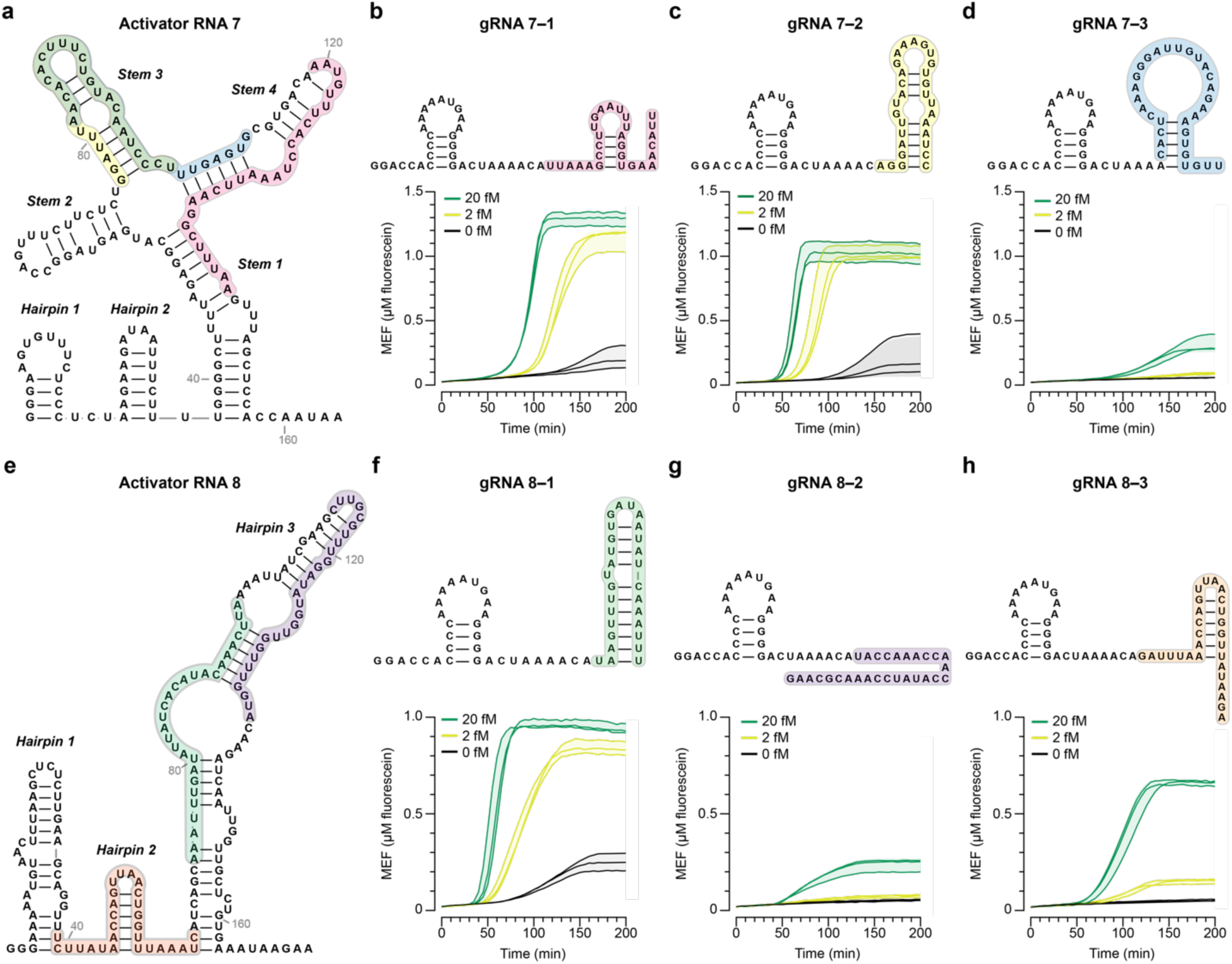
Screening of LbuCas13a gRNAs identifies factors that could impact cleavage efficiency. **a**, Predicted secondary structure of activator RNA 7 generated from NASBA with Primer Set 7. Regions targeted by gRNAs are shaded in different colors. Fluorescence kinetics from NASBA-Cas13a at varying concentrations of synthetic SARS-CoV-2 genome with **b**, gRNA 7–1, **c**, gRNA 7–2, or **d**, gRNA 7–3 with predicted secondary structures of each gRNA shown above. **e,** Predicted secondary structure of activator RNA 8 generated from NASBA with Primer Set 8. Regions targeted by gRNAs are shaded in different colors. Fluorescence kinetics from NASBA-Cas13a at varying concentrations of synthetic SARS-CoV-2 genome with **f**, gRNA 8–1, **g**, gRNA 8–2, or **h**, gRNA 8–3 with predicted secondary structures of each gRNA shown above. Data are n=3 independent biological replicates, each plotted as a line with raw fluorescence standardized to MEF. Shading indicates the average of the replicates ± standard deviation.

We chose to develop the system in the context of detecting specific sequences of the SARS-CoV-2 genome, and also demonstrated that the device can be used to detect the plant virus cucumber mosaic virus (CMV) in plant lysate. We first demonstrate that NASBA-Cas13a can be performed in a one-pot isothermal reaction using *Leptotrichia buccalis* (Lbu) Cas13a. We then develop an in-house reaction formulation that provides flexibility for optimization by adjusting individual components and their concentrations. With the in-house formulation, we identify design rules for NASBA primer sets, as well as LbuCas13a gRNAs, to achieve efficient and sequence-specific detection of target RNAs. We next used mechanistic modeling of NASBA-Cas13a to better understand this system. We reasoned that the use of the well-characterized processes of reverse transcription, transcription, and nuclease activity would make the combined NASBA-Cas13a reaction scheme amenable to mechanistic modeling, which we used to explore design principles of the system. We constructed an ordinary differential equation (ODE) mechanistic model describing the core reaction scheme processes, as well as potential off target reactions that could occur in a one-pot formulation of the system. The use of a high-throughput acoustic liquid handling instrument enabled the generation of a large training dataset that was used with the Generation and Analysis of Models for Exploring Synthetic Systems (GAMES)^28^ framework to develop and train the model. We found that the variability of the high throughput generated data created challenges to model building. However, we were able to extract non-intuitive design principles related to reaction inhibition due to high concentrations of certain enzyme species. The introduction of empirical heuristics was necessary to recapitulate measured trends, pointing to potentially unknown biochemical mechanisms that may be at play in one-pot reaction formulations. Finally, we explore reaction optimizations and show the ability to detect hundreds of aM of the SARS-CoV-2 genomic sequence.

Overall, this study provides an additional technique to the repertoire of nucleic acid detection technologies and sets the stage for combining high-throughput experimental screening of reaction conditions with mechanistic modeling to drive further innovation of these technologies.

## MATERIALS AND METHODS

### Bacterial strains and growth medium

*E. coli* strain K12 (Turbo Competent *E. coli*, NEB #C2984) was used for cloning. *E. coli* strain Rosetta 2(DE3)pLysS (Novagen #71401) was used for recombinant protein expression. Luria Broth supplemented with the appropriate antibiotic(s) (100 µg/mL carbenicillin, 100 µg/mL kanamycin, and/or 34 µg/mL chloramphenicol) was used as growth medium.

### Plasmids and genetic parts assembly

DNA oligonucleotides for cloning and sequencing were synthesized by IDT. NASBA primers were ordered PAGE-purified to minimize any off-target NASBA products. Genes encoding gRNAs and SARS-CoV-2 and CMV input RNA fragments were synthesized either as gBlocks or Ultramers (IDT). A plasmid for expressing LbuCas13a was obtained from Addgene (#83482).

Transcription templates for expressing gRNA variants and SARS-CoV-2 or CMV input RNA fragments were generated by PCR (Phusion high-fidelity PCR kit, NEB #E0553) of the gBlock or Ultramer template that included a T7 promoter and the gRNA or input RNA coding sequence. For the gRNA-expressing templates, an additional cis-cleaving Hepatitis D ribozyme and an optional T7 terminator were included on the 3’ end of the gRNA coding sequence. We define the T7 promoter as a minimal 17 base pair (bp) sequence (TAATACGACTCACTATA) excluding the first G that is transcribed. PCR-amplified templates were purified (QIAquick PCR purification kit, Qiagen #28106) and verified for the presence of a single DNA band of the expected size on a 1% TAE-agarose gel. DNA concentrations were measured using a Qubit dsDNA BR assay kit (Invitrogen #Q32853). Plasmids and DNA templates were stored at 4°C. Oligonucleotides and primers are listed in **Supplementary Data 1.**

### RNA expression and purification

Guide RNAs were expressed from a transcription template encoding a 3’ cis-cleaving Hepatitis D ribozyme (**Supplementary Data 1**) using overnight IVT at 37°C with the following components: IVT buffer (40 mM Tris-HCl pH 8, 8 mM MgCl_2_, 10 mM DTT, 20 mM NaCl, and 2 mM spermidine), 11.4 mM NTPs pH 7.5, 0.3 U thermostable inorganic pyrophosphatase (NEB #M0296S), 100 nM transcription template, 50 ng T7 RNAP, and MilliQ ultrapure H_2_O to a total volume of 100 µL. Overnight reactions were ethanol-precipitated and purified by resolving on a 15% urea-PAGE-TBE gel, isolating the band of the expected size (∼60 nt) and eluting at 4°C overnight in MilliQ ultrapure H_2_O. Eluted gRNAs were ethanol-precipitated, resuspended in MilliQ ultrapure H_2_O, quantified using a Qubit RNA BR assay kit (Invitrogen #Q10211), and stored at –20°C. The SARS-CoV-2 and CMV input RNA fragments used in **Supplementary Fig. 3a–c** (which did not contain the ribozyme sequence) were expressed and purified as described above.

### LbuCas13a expression and purification

LbuCas13a expression and purification was carried out as described previously^29^ with minor modifications. The LbuCas13a expression plasmid (N-terminally tagged with a His_6_-MBP-TEV cleavage site) was transformed into Rosetta 2(DE3)pLysS *E. coli*. A 4 L cell culture was grown in Luria Broth at 37°C, induced with 0.5 mM of IPTG at an optical density (600 nm) of ∼0.5, and grown overnight at 16°C. Cultures were pelleted by centrifugation (4,000 x g) and resuspended in lysis buffer (50 mM Tris-HCl pH 7, 500 mM NaCl, 5% glycerol, 1 mM TCEP, and EDTA-free protease inhibitor (Roche)). Resuspended cells were lysed on ice through ultrasonication, and insoluble materials were removed by centrifugation. Clarified supernatant containing LbuCas13a was purified using His-tag affinity chromatography with a Ni-NTA column (HisTrap FF 5mL column, GE Healthcare Life Sciences) followed by size exclusion chromatography (Superdex HiLoad 26/600 200 pg column, GE Healthcare Life Sciences) using an AKTAxpress fast protein liquid chromatography (FPLC) system. The His_6_-MBP tag was removed from the eluted fractions by adding His_6_-tagged TEV protease in 2 L cleavage buffer (50 mM Tris-HCl, 250 mM NaCl, 1 mM EDTA, 1 mM TCEP, 5% glycerol) at 37°C for 1 h and then at 4°C overnight. The TEV-cleaved LbuCas13a was buffer-exchanged at 4°C into 3 L of the final storage buffer (20 mM Tris-HCl pH 7, 200 mM KCl, 5% glycerol, 1 mM TCEP), which was split into three 1 L buffers that were swapped out every 30 min. The His_6_-tagged TEV protease was removed by reloading the fractions onto a Ni-NTA column (HisTrap FF 5mL column, GE Healthcare Life Sciences) and collecting the fractions from a 5% imidazole wash. Protein concentrations were determined using a Qubit protein assay kit (Invitrogen #Q33212). Protein purity and size were validated on an SDS-PAGE gel (Bio-Rad Mini-PROTEAN TGX and Mini-TETRA cell). Purified proteins were stored at –80°C.

### NASBA-Cas13a with commercial NASBA reactions

NASBA-Cas13a reactions depicted in **Supplementary Fig. 2** were performed using the commercial NASBA Liquid kits from Life Sciences Advanced Technologies Inc. (SKU #NWK-1). 3X Reaction Buffer and 6X Nucleotide Mix were combined with 250 nM of primer each and the input viral RNA template (PAGE-purified synthetic SARS-CoV-2 fragment or CMV-infected plant lysate) at varying concentrations to a volume of 7.5 µL to make 1.3X NASBA master mix. The master mix was heated at 65°C for 2 min and cooled to 41°C for 5 min to facilitate binding of the primers to the input viral RNA template. 2.5 µL of the Enzyme Mix and 10 µL of LbuCas13a cleavage reaction mix (see **In-house NASBA-Cas13a** for details) were added to the master mix to initiate the reaction, and fluorescence was monitored on a plate reader (see **Plate reader quantification and micromolar equivalent fluorescein (MEF) standardization** for details). The final concentration of each reaction component is listed in **Supplementary Data 3.**

### In-house NASBA-Cas13a

An in-house NASBA-Cas13a reaction was prepared by combining three different reaction mixes – NASBA master mix, NASBA enzyme mix, LbuCas13a cleavage reaction mix – that were prepared separately to a final volume of 20 µL. The NASBA master mix was prepared by combining the following components (listed at final concentration): NASBA reaction buffer (50 mM Tris-acetate, 8 mM Mg-acetate, 75 mM K-acetate, 10 mM DTT, pH 8.3), 12 mM Tris-buffered NTPs, 4 mM dNTPs (NEB #N0447L), 250 nM PAGE-purified forward and reverse primers, 5 mM fresh DTT, 15% DMSO, and an input RNA at varying concentrations. This master mix was incubated at 65°C for 5 min and cooled to 37°C to promote primer binding. In parallel with the above steps, the LbuCas13a cleavage reaction mix was prepared by first incubating the gRNA at 95°C for 5 min and snap-cooling on ice. Then, the following components were combined (listed at final concentration): cleavage buffer (40 mM Tris-HCl, 60 mM NaCl, 6 mM MgCl_2_, pH 7.3), 90 nM LbuCas13a, 45 nM gRNA, RNase inhibitor (Invitrogen #10777019), and 2.5 µM RNA reporter (6’FAM-UUUUU-IABkFQ). The cleavage reaction mix was incubated at 37°C for ∼10 min to promote the complexing of LbuCas13a and gRNA. During these incubation steps, a NASBA enzyme mix was prepared by combining the following components (listed at final concentration): 0.1 µg/µL BSA (NEB #B9000S), 5 U/µL T7 RNAP (NEB #M0460T), 0.0005 U/µL RNase H (NEB #M0297L), and 2.5 U/µL M-MuLV RT (NEB #M0253L) unless indicated otherwise. Lastly, the NASBA master mix, NASBA enzyme mix, and the LbuCas13a cleavage reaction were all combined and mixed by gentle pipetting.

### RNA extraction from NASBA

For RNA products in the gel image in **Fig. 1f**, NASBA reactions were set up as described above, followed by phenol-chloroform extraction and ethanol precipitation to remove any proteins. Reactions were rehydrated in 1X TURBO^TM^ DNase buffer with 2U TURBO^TM^ DNase (Invitrogen #QAM2238) to a total volume of 20 µL and incubated at 37°C for 30 min to remove any DNA products generated during NASBA. Phenol-chloroform extraction followed by ethanol precipitation was performed again to remove DNase, with rehydration in MilliQ ultrapure H_2_O. Concentrations of extracted RNA products were measured using a Qubit RNA HS assay kit (Invitrogen #Q32852), and they were stored at –20° C until analysis. PAGE analysis of extracted RNA products used 10% urea-PAGE-TBE gels. Gels were imaged using a ChemiDoc^TM^ Touch gel imaging system (BioRad Image Lab Touch Software 1.2.0.12).

### Sequential NASBA

Two separate reactions per RT were prepared for sequential NASBA reactions in **Supplementary Fig. 3a.** First, all reactions were prepared by combining the following components (listed at final concentration): NASBA reaction buffer (50 mM Tris-acetate, 8 mM Mg-acetate, 75 mM K-acetate, 10 mM DTT, pH 8.3), 1 mM dNTPs (NEB #N0447L), 250 nM of the first primer (IDT, PAGE-purified), and 10 nM SARS-CoV-2 input RNA fragment (**RNA expression and purification**). The reaction mixtures were incubated at 65°C for 2 min and cooled to 37°C to promote initial primer binding. The first cDNA synthesis was initiated by adding 1 U/µL RT at final concentration (NEB #M0277L for AMV RT, NEB #M0368L for ProtoScript II RT, and NEB #M0253L for M-MuLV RT). After 20 minutes of incubation at 37°C, 0.005 U/µL of RNase H (NEB #M0297L) was added to each reaction to digest the input RNA fragment. After an additional 20 min of incubation at 37°C for 20 min, one of the reactions was placed on ice to halt the reaction until further purification (Reaction 1). 250 nM of the second primer (IDT, PAGE-purified) was added to the other reaction, followed by a 40-minute incubation at 37°C to complete dsDNA synthesis (Reaction 2). Then, all reactions were treated with 4 M NaOH at 95°C for 5 minutes to remove any residual RNA and neutralized with HCl. Reactions were ethanol-precipitated, and DNA products were analyzed on a 10% PAGE-TBE gel without denaturing agent.

For NASBA products indicated in **Supplementary Fig. 3b,c**, two reactions were prepared by combining the following components (listed at final concentration): NASBA reaction buffer (50 mM Tris-acetate, 8 mM Mg-acetate, 75 mM K-acetate, 10 mM DTT, pH 8.3), 1 mM dNTPs (NEB #N0447L), 250 nM of the first primer (IDT, PAGE-purified), and 50 nM input RNA fragment (SARS-CoV-2 fragment for **b** and CMV fragment for **c**). The mixtures were incubated at 65°C for 2 min and cooled to 41°C to promote the initial primer binding. Then, the first cDNA synthesis was initiated by adding 0.5 U/µL AMV RT (NEB #M0277L). After 30 min of incubation at 41°C, 0.005 U/µL RNase H (NEB #M0297L) was added to one of the reactions. The other reaction was placed on ice to halt the first cDNA synthesis temporarily. After incubating for 20 min with RNase H at 41°C, 250 nM of the second primer (IDT, PAGE-purified) was added to both reactions, and the mixtures were incubated for 30 min at 41°C to complete dsDNA synthesis. Then, all reactions were treated with 4 M NaOH at 95°C for 5 minutes to remove any residual RNA and neutralized with HCl. Reactions were ethanol-precipitated, and DNA products were analyzed on a 10% PAGE-TBE gel without denaturing agent.

### Plate reader quantification and micromolar equivalent fluorescein (MEF) standardization

A NIST traceable standard (Invitrogen #F36915) was used to convert fluorescence signal in arbitrary units to micromolar equivalent fluorescein (MEF). Serial dilutions from a 50 µM stock were prepared in 100 mM sodium borate buffer at pH 9.5, including a 100 mM sodium borate buffer blank (12 samples in total). For each concentration, three replicates of samples were prepared, and fluorescence was read at an excitation wavelength of 490 nm and emission wavelength of 525 nm for 6-FAM (fluorescein)-activated fluorescence (Synergy H1, BioTek Gen5 v2.04). Fluorescence values for any fluorescein concentration in which a single replicate saturated the plate reader were excluded from analysis. The remaining replicates were averaged at each fluorescein concentration, and the average fluorescence of the blank was subtracted from all values. To estimate a conversion factor, linear regression was performed for concentrations within the linear range between the measured fluorescence values in arbitrary units and the concentration of fluorescein. For each plate reader and gain setting, we estimated a linear conversion factor that was used to convert arbitrary fluorescence values to MEF (**Supplementary Data 3**).

To characterize reaction kinetics, 19 µL reactions were loaded into a 384-well optically clear, flat-bottom plate using a multichannel pipette and covered with a plate seal, and their signals were measured via plate reader (Synergy H1, BioTek Gen5 v2.04). Kinetic analysis of 6-FAM (fluorescein)-activated fluorescence was performed by reading the plate at 5-minute intervals with excitation and emission wavelengths of 490 nm and 525 nm, respectively, for four hours at 37°C. Arbitrary fluorescence units were converted to MEF using the appropriate calibration conversion factor. No background subtraction was performed in the analysis of these reactions.

### RNA structure prediction

Input viral RNA templates and gRNA secondary structures were predicted using RNAStructure^30^ and NUPACK^31^ at a temperature of 37°C with their respective default parameters. Both prediction algorithms were used for all RNAs. If there was a discrepancy between the two predicted structures, the secondary structure predicted with NUPACK was used since its default parameters resemble the reaction conditions more closely.

### High-throughput screening of NASBA-Cas13 reactions with an Echo liquid handling platform

NASBA enzyme mixes testing different enzyme concentrations were constructed using a liquid-handling robot (Beckman Coulter, Echo 550) as previously described (**In-house NASBA-Cas13a**) with minor modifications to accommodate the requirements of the Echo platform. A 2 µg/µL BSA solution (in water) was transferred from a 384-well polypropylene 2.0 Plus Source microplate (Beckman) using the 384PP_Plus_BP fluid type into a 384-well destination plate (BioRad, HSP 3805) using the Echo 525 (Beckman Coulter). While the mixture was being dispensed, NASBA enzymes were diluted to appropriate concentrations for the reaction conditions to be tested onto a 384-well polypropylene 2.0 Plus Source microplate (Beckman). Once the BSA–water mixture dispense was complete, NASBA enzyme dilutions from the source plate were dispensed onto the same destination plate by the Echo 550 suing the 384PP_AQ_CP fluid type. During the NASBA enzyme dispense, a NASBA master mix and a LbuCas13a cleavage reaction mix were prepared following the **In-house NASBA-Cas13a** protocol. Once the NASBA enzyme mix dispense was complete, the NASBA master mix and LbuCas13a cleavage reaction mix were manually pipetted onto the destination plate using a multichannel pipette (Integra Voyager). Then, the destination plate was sealed was loaded onto a plate reader (Synergy H1, BioTek Gen5 v2.04), and the readout was measured (**Plate reader quantification and micromolar equivalent fluorescein (MEF) standardization**).

Using this protocol, the following enzyme concentrations were tested: (1) 1, 5, and 20 U/µL T7 RNAP, (2) 0.5, 2.5, and 10 U/µL M-MuLV RT, and (3) 0.001, 0.005, and 0.02 U/µL RNase H. For each of these concentrations, two concentrations of Cas13a-gRNA complex were tested: (2.25 and 45 nM) where Cas13a was added in two-fold molar excess of gRNA in the assembly step of the LbuCas13a cleavage reaction mix (**In-house NASBA-Cas13a).** In addition, three SARS-CoV-2 genome (input RNA) concentrations (0, 1, and 10 fM) were used to initiate reactions. Using this screen setup, 162 triplicate reaction conditions were tested: (2 Cas13a-gRNA × 3 input RNA × 3 T7 RNAP × 3 M-MuLV RT × 3 RNase H) × 3 replicates = 486 reactions. Screening was performed in two batches: one for conditions with 2.25 nM Cas13a-gRNA and the other for conditions with 45 nM Cas13a-gRNA. This screen setup was performed a total of three separate times (Echo replicate 1, 2, and 3) for a total of 1,458 reactions.

### Iterative model development and analysis

We performed iterative model formulation and parameter estimation based on a previously described workflow for dynamic model development^28^. To initialize this process and set criteria for success, we defined a set of qualitative modeling objectives (**Table 1**), chose a subset of the data to use as training data for each of the three high-throughput screening experiments (**Data Sets 1-3**, in order of collection date) (**Supplementary Data 3**, **Supplementary Fig. 6b**), and formulated a base case model. Next, we evaluated the parameter estimation method to ensure that the method could identify the best possible parameter sets given the structure of the base case model and each training data set (**Supplementary Note 2**). This process serves as a positive control for parameter estimation and ensures that the method used is implemented correctly and is appropriate for the given parameter estimation problem. Then, we used the same parameter estimation method to estimate parameters based on each of the training data sets independently. We inspected the agreement between each experimental data set and the corresponding simulated values in the context of the modeling objectives, proposed mechanistic updates intended to improve the agreement when observations motivated such amendment, and mathematically implemented these updates in a new model. We iterated this process until we identified a model that satisfied all modeling objectives for each data set (**Table 1**, **Fig. 4** for **Data Set 2, Supplementary Fig. 22** for **Data Set 1**, and **Supplementary Fig. 24** for **Data Set 3**).

**Table 1.** Summary of modeling objectives and candidate models for each data set^1^.

|  | Model A |  |  | Model B |  |  | Model C |  |  | Model D |  |  |
| --- | --- | --- | --- | --- | --- | --- | --- | --- | --- | --- | --- | --- |
| Additional mechanism→<br>Modeling objective ↓ | Base case |  |  | Cas13a-gRNA deactivates over time |  |  | Negative relationship between transcription rate and T7 RNAP |  |  | Non-monotonic relationship between RNase H cleavage rate and RNase H |  |  |
|  | Data set 1 | Data set 2 | Data set 3 | Data set 1 | Data set 2 | Data set 3 | Data set 1 | Data set 2 | Data set 3 | Data set 1 | Data set 2 | Data set 3 |
| Objective 1: Time course trajectories have a sigmoidal shape | Yes | Yes<sup>2</sup> | Yes<sup>2</sup> | Yes | Yes<sup>2</sup> | Yes<sup>2</sup> | Yes | Yes | Yes | Yes | Yes | Yes |
| Objective 2: Plateaus in readout can occur at various times depending on the condition | Yes | Yes<sup>2</sup> | Yes<sup>2</sup> | Yes | Yes<sup>2</sup> | Yes<sup>2</sup> | Yes | Yes | Yes | Yes | Yes | Yes |
| Objective 3: The magnitude of the plateau can vary depending on the condition | No | No | No | Yes | No | No | Yes | Yes | Yes | Yes | Yes | Yes |
| Objective 4: Increasing RT to a relatively high concentration increases the readout | No | No | Yes | Yes | Yes | Yes | Yes | No | Yes | Yes | Yes | Yes |
| Objective 5: Increasing T7 RNAP to a relatively high concentration decreases the readout | No | No | No | No | No | No | Yes | Yes | Yes | Yes | Yes | Yes |
| Objective 6: Increasing RNase H to a relatively high concentration increases, then decreases the readout | No | No | No | No | No | No | No | No | No | No<sup>3</sup> | Yes | Yes |
| Quantitative agreement (R<sup>2</sup>) | 0.43 | 0.26 | 0.31 | 0.53 | 0.27 | 0.33 | 0.99 | 0.87 | 0.78 | 0.99 | 0.96 | 0.85 |
| Quantitative agreement (MSE) | 0.062 | 0.040 | 0.051 | 0.045 | 0.037 | 0.048 | 0.0021 | 0.0066 | 0.018 | 0.001<br>9 | 0.002<br>6 | 0.016 |

<sup>1</sup>Each column is an additional mechanism added to the model. For example, Model C includes mechanisms from Models A and B. ‘Yes’ indicates that a version met an objective, and ‘No’ indicates that it did not. Calibration and analysis of suboptimal candidate models are described in **Supplementary Fig. 11**. Quantitative agreement is reported as the MSE or R<sup>2</sup> between experimental data and simulated data for the subset that was used for training in **Fig. 4 (Supplementary Fig. 6b)**.
<sup>2</sup>The determination of whether these objectives were met for **Data Sets 2 and 3** is based only on the quantitative metrics from the Hill fit (i.e., a high R<sup>2</sup> value between the trajectories and Hill fits and a range of $t_{1/2}$ values) (**Supplementary Note 3**).
<sup>3</sup>Although this modeling objective was not met for **Data Set 1**, given the experimental error in the RNase H sweeps it is unclear whether this data set has a non-monotonic relationship between RNase concentration and readout, which is indicative of potential error in the amount of each reagent dispensed from the liquid handling robot for each experiment.

### Approximation of dynamics

Simulations were run using custom Python scripts (Python 3.9.12) and Python package SciPy’s^32^ solve_ivp solver with the LSODA algorithm. The Jacobian matrix was provided and explicitly calculated for each timestep, rather than relying on a finite difference approximation. Initially, we used the default solve_ivp error tolerances (an absolute tolerance of 10^-6^ and a relative tolerance of 10^-3^) to run simulations.

### Fluorescence data normalization

Raw fluorescence data (i.e., **Data Set 2** in **Fig. 3**), were normalized by the following method for two reasons: (1) to enable comparison between the two batches of experiments performed on different dates, and (2) to enable comparison between experimental observations and simulations. Raw experimental fluorescence values were first converted to absolute units, MEF (µM fluorescein), using the method described above (**Plate reader quantification and micromolar equivalent fluorescein (MEF) standardization**). Then, the MEF value at each time interval was normalized to the maximum value over the entire data set. An analogous normalization was applied to each simulated data set in which each data point was divided by the maximum value in the simulated data set.

**Fig. 3.**
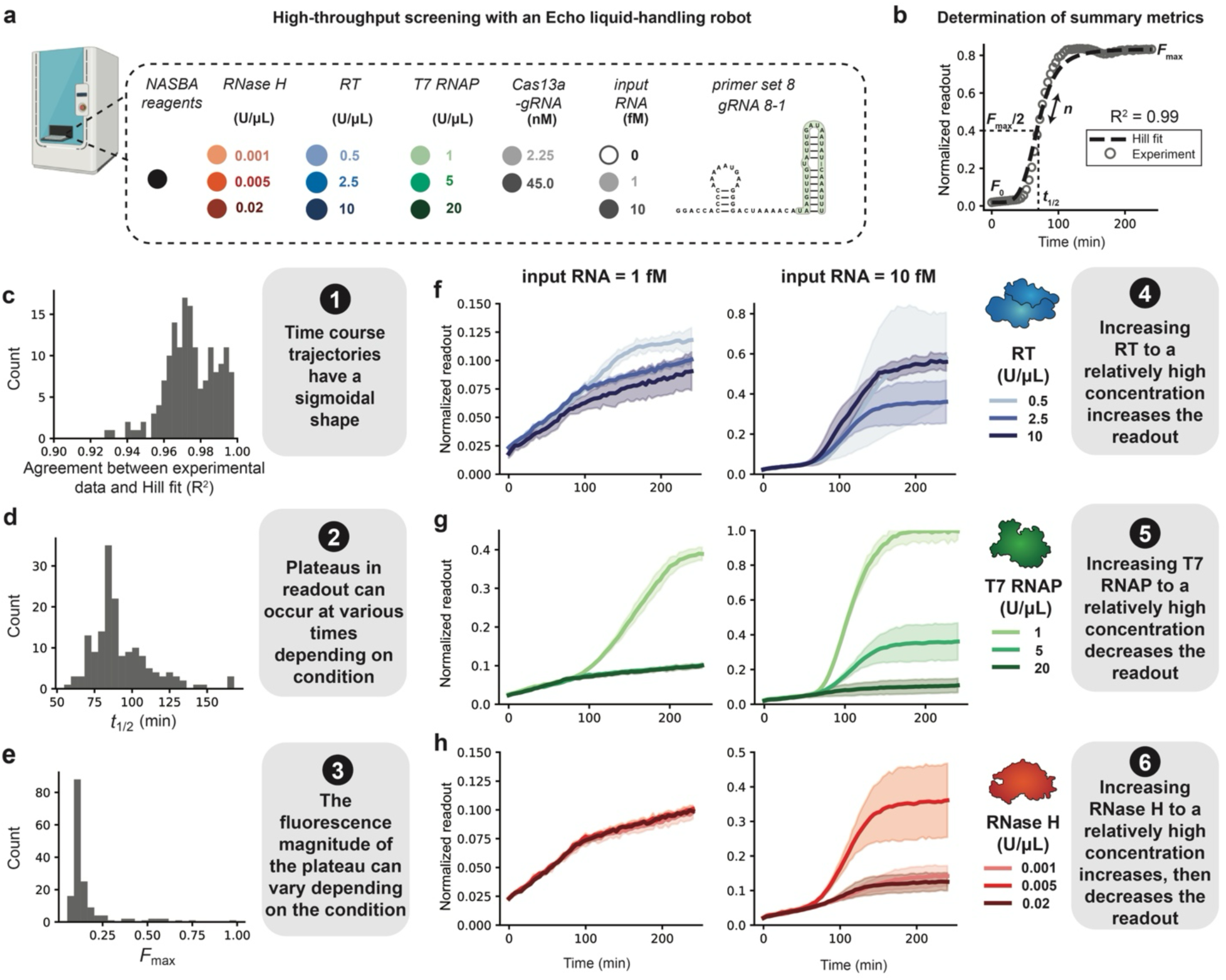
High-throughput screening of the enzyme concentration landscape suggests model assumptions and reveals reaction design principles, shown for Data Set 2. **a**, Different amounts of input RNA, RT, T7 RNAP, RNase H, and Cas13a-gRNA were dispensed in triplicate (independent replicates) using an Echo liquid handling platform. Assembled NASBA-Cas13a reactions were run and fluorescence data collected and averaged across triplicate measurements to arrive at a mean dynamic trajectory. The dynamic trajectory was then normalized by the maximum readout value, such that the maximum readout value across the entire experiment (all conditions) was set to 1. **b**, Hill functions were fit to each normalized time course trajectory, and summary metrics (*n*, *t*_1/2_, *F*_0_, and *F*_max_) were parameterized. A representative time course trajectory and Hill plot is shown as an example. **c**, For each time course, R^2^ values for the normalized experimental data (points) and Hill fit (dotted line) were calculated and plotted as a histogram. Histograms of values across all conditions were computed for: **d**, *t*_1/2_ , and **e**, *F*_max_. f– h, Time course trajectories for data subsets varying: **f**, RT, **g**, T7 RNAP, and **h**, RNase H, each using two different input RNA concentrations. Shading indicates the average of the triplicates ± standard deviation. This process was repeated for each experimental data set, but Data Set 2 is highlighted here because it was in closest alignment with all modeling objectives.

**Fig. 4.**
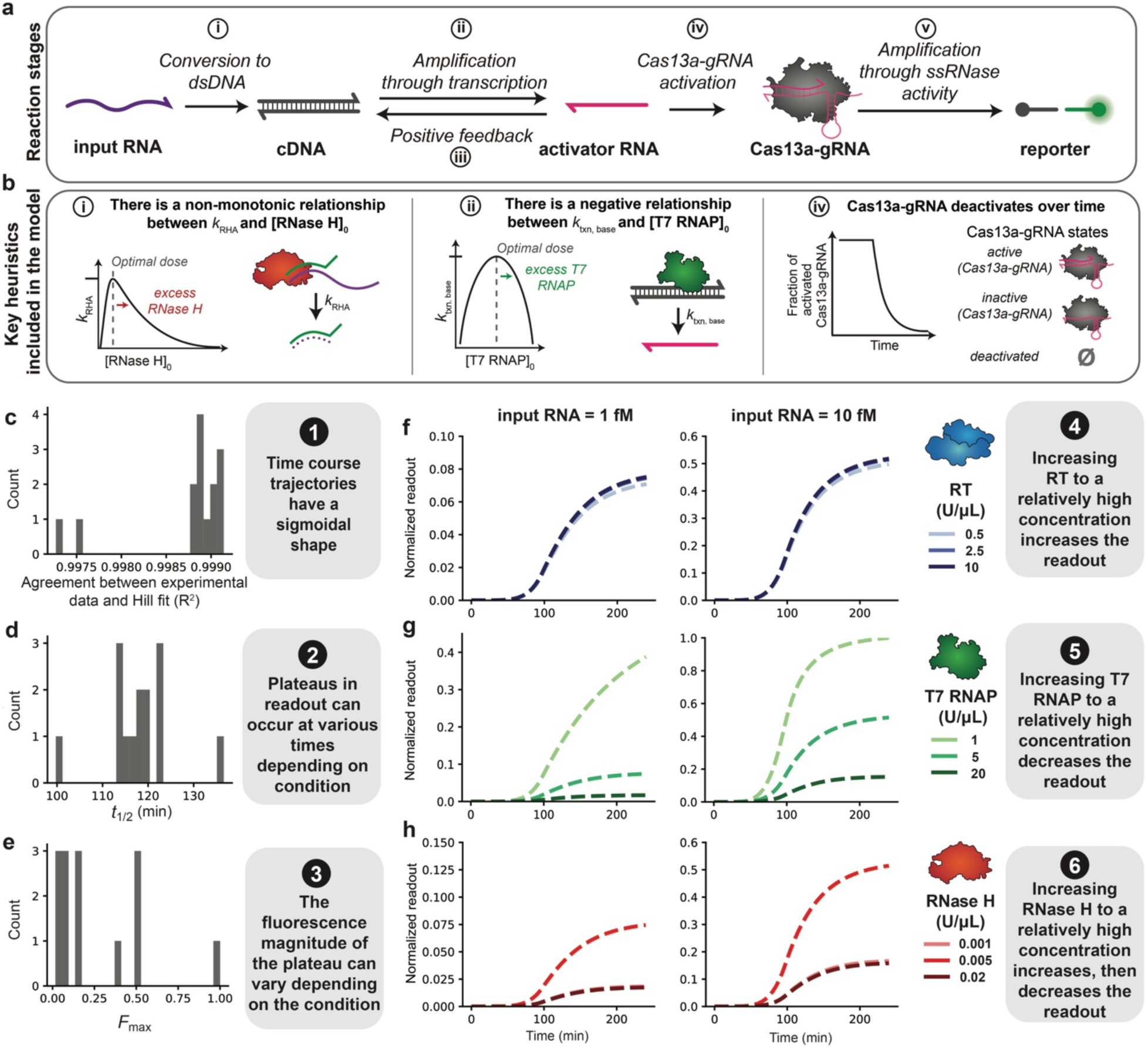
Mathematical modeling recapitulates key experimental observations. **a**, Schematic of key reaction stages (top) and mechanisms (bottom) in the model. A more detailed depiction of the model is in **Supplementary Fig. 21**. **b,** Key mechanisms included in the model. Each mechanism is involved in the reaction stage indicated to the left of each mechanism description. **c-e**, Hill-like functions were fit to each simulated time course trajectory, and summary metrics (*n*, *t*_1/2_, *F*_0_, and *F*_max_) were parameterized (**Fig. 3b** is a visual representation of these metrics). **c**, For each time course, R^2^ for the normalized simulated data and Hill fit was calculated; values are plotted as a histogram. Histograms of values across all conditions in the simulated training data set were calculated for: **d**, *t*_1/2_, and **e**, *F*_max_, **f–h**, Time course trajectories for simulated data subsets: **f**, mid-range RNase H and T7 RNAP and high Cas13a-gRNA, **g**, mid-range RNase H and RT and high Cas13a-gRNA, and **h**, mid-range T7 RNAP and RT and high Cas13a-gRNA.

### Definition of training data for model development

We chose not to train the model on conditions including only background signal because *F*_max_ (maximum fluorescence) values for these conditions were generally below practical visibility. For this reason, we omitted the conditions lacking input RNA and the conditions with low Cas13a-gRNA from each training data set. We then selected a subset of conditions for model training from each Echo replicate, consisting of concentration sweeps of one NASBA enzyme while holding mid-level concentrations of the other two enzymes. For example, this selected subset includes the conditions with 0.001, 0.005 and 0.02 U/μL RNase H, each with 2.5 U/μL RT and 5 U/μL T7 RNAP. These training data are referred to as **Data Set 1, 2 and 3**, corresponding to the three runs of the Echo screen (Echo replicates) (**Supplementary Fig. 6b**). We chose to incorporate data from each of the Echo replicates into the training data to gain a holistic, mechanistic understanding of the system that could sufficiently recapitulate experimental observations despite variation between experiments. The remaining data was held out for validation and are referred to as out-of-sample from each Echo replicate.

### Pre-processing of training data

We pre-processed the data to remove conditions for which there was low confidence due to high measurement error. First, we calculated the mean proportion of measurement error 𝑝*_j_* for each condition (set of unique enzyme concentrations) *j*, starting with **Data Set 1**:

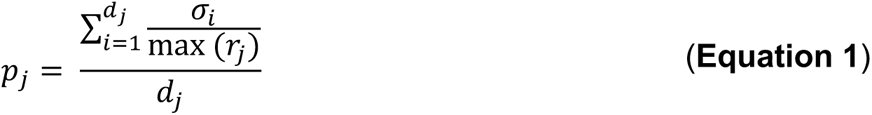

Here, 𝑑*_j_* is the number of data points collected for each condition *j*, 𝜎*_i_* is the measurement error (standard deviation) associated with data point *i* in condition *j*, and max(𝑟*_j_*) is the maximum readout value for condition *j*. The distribution of 𝑝*_j_* across all conditions (**Supplementary Fig. 9a**) indicated that a small subset of conditions in **Data Set 1** had high mean proportion measurement error (the highest 𝑝*_j_* was nearly 0.80, or 80%). Including conditions with high measurement error can bias parameters by fitting to random trends in noise instead of underlying biological mechanisms.

To determine which conditions to remove from the training data for **Data Set 1**, we investigated the time course trajectory for each replicate in the NASBA enzyme sweeps, excluding conditions for which input RNA = 0 or Cas13a-gRNA = 2.25 nM (**Supplementary Fig. 9b**). The condition with 𝑝*_j_* ≥ 0.30 had one replicate with near-zero readout regardless of the time point, in contrast to the other replicates in the condition, potentially suggesting an experimental error in implementing this condition. Therefore, we chose to remove the condition with 𝑝*_j_* > 0.30 from the training data for **Data Set 1** (**Supplementary Fig. 6b**). We calculated the mean proportion error distributions for **Data Sets 2** and **3** (**Supplementary Fig. 9c,d**, respectively), but there were no conditions with 𝑝*_j_* ≥ 0.30 within the subset of conditions used for the training data, so no conditions were removed from the training data for either data set.

### Cost function

The cost function, which calculates the agreement between experimental and simulated data, was defined as the mean of squared error (MSE) evaluated between each normalized experimental and normalized simulated data point. In the equation below, 𝑑 is the total number of data points in the training data set, 𝑦*_k_^exp^* is the k^th^ datapoint in the normalized experimental data, and 𝑦*_k_*(𝜃) is the normalized simulated value of the k^th^ datapoint using the parameter set 𝜃.

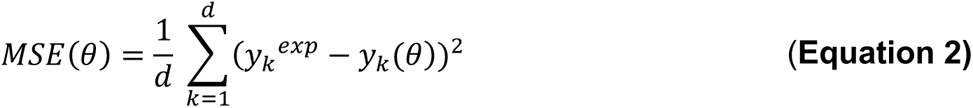

#### Cost function filter

We applied a cost function filter to remove from consideration any parameter sets yielding low (desirable) cost function values that were undesirable for other reasons. We noticed that parameter sets yielding very low simulated readout across all conditions were still able to achieve low cost function values due to the maximum value-based normalization strategy that we used. Therefore, we removed all parameter sets yielding maximum simulated fluorophore readout values of less than 2000 nM, as we expect the maximum value in the experimental data set to be at least on the order of ∼2500 nM, which is the initial concentration of quencher-fluorophore.

#### Coefficient of determination

The coefficient of determination (R^2^) was used along with MSE to evaluate the goodness of fit between simulated data and experimental data.

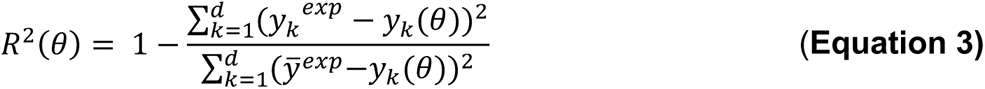

Here, 𝑑 is the number of data points in the training data set, 𝑦*_k_^exp^* is the k^th^ data point in the normalized experimental data, 𝑦*_k_*(𝜃) is the normalized simulated value of the k^th^ data point using the parameter set 𝜃, and 𝑦*^exp^* is the mean of the training data set. R^2^ is a more interpretable metric than MSE because the magnitude of MSE values depends on many factors such as the number of data points^28^. Possible R^2^ values span 0 to 1, with R^2^ = 1 indicating perfect agreement between the two data sets.

### Parameter estimation method

A multi-start local optimization algorithm was used to estimate parameters (**Supplementary Fig. 10a, Supplementary Note 3**). First, a global search with *n*_search_ total parameter sets was performed and the cost function was calculated for each parameter set. The *n*_init_ parameter sets with the lowest cost function values were used to initialize optimization runs using the Levenberg-Marquardt optimization algorithm. The resulting parameter set with the lowest cost function value following optimization was chosen as the best (i.e., calibrated) parameter set. This algorithm was implemented using custom Python scripts along with Python packages SALib^33^ for global search and LMFit^34^ for optimization.

While the default numerical tolerances kept computational time minimal, simulated concentration values sometimes took negative values, which is an unphysical result due to numerical error. To check whether estimated parameters were relatively insensitive to these errors, we reduced the absolute tolerance to 10^-13^ and relative tolerance to 10^-10^, reran the optimization using the same parameters for initialization as in the default error tolerance runs, and found that the optimization results were consistent when the cost function was low. When the model resulted in a poor fit to the training data results were not always consistent, suggesting that the difference in the parameters in these cases was a result of the model formulation. A representative example comparing the time course trajectories with the default versus decreased ODE solver tolerances for the final model for **Data Set 2** is in **Supplementary Fig. 25**.

### Parallelization of computational tasks

Simulations were parallelized across eight independent cores (chosen based on the number of cores available in the hardware used to run the simulations) to improve computational efficiency. Parallelization was implemented using custom scripts and the multiprocessing package in Python.

### Sensitivity analysis

We performed a sensitivity analysis on the calibrated parameters for the final model for each data set to determine which parameters had the greatest impact on the simulated time course trajectories and overall fit to experimental data. We independently varied each parameter by ± 10% of the calibrated value and calculated the *t*_1/2_ (time to reach half-maximum readout), *F*_max_, and MSE for each parameter variation. The percent change in each metric was calculated relative to the metric for the calibrated parameter set to quantify the model’s sensitivity to each parameter.

### Definition of test data

We selected five sets of test data for the final models trained on **Data Set 1, 2,** and **3**, including the training data from the other replicates generated with the Echo liquid handler. For example, the test data for the final model trained on **Data Set 1** includes the out-of-sample data from the first Echo replicate (**Supplementary Fig. 6b**), training **Data sets 2** and **3**, and out-of-sample data from the 2^nd^ and 3^rd^ Echo replicate. We used the final model for each data set to simulate time course trajectories for each condition in the test data set, and calculated MSE and R^2^ metrics to quantify the fit.

## RESULTS

### Screening of NASBA reverse transcriptases (RTs)

Before developing an in-house NASBA formulation, we used a commercial kit to assess the feasibility of combining NASBA and CRISPR-Cas13a cleavage in a one-pot isothermal reaction (**Fig. 1a, Materials and Methods**). The NASBA primers targeted the genome of either SARS-

CoV-2 or cucumber mosaic virus (CMV)^35^ (**Supplementary Data 1**) and were designed to yield an RNA product complementary to the pathogen sequence (**Supplementary Fig. 1**). For reactions detecting the SARS-CoV-2 genome (synthesized by TWIST, SKU 102019), we observed an input RNA concentration-dependent effect on fluorescent signal (**Supplementary Fig. 2a, b**). We also tested for detection of CMV genome from infected plant lysate and confirmed that the reaction can take place in a complex matrix (**Supplementary Fig. 2c, d**).

For the SARS-CoV-2 detecting reaction, we observed a substantial signal in the absence of the input RNA (**Supplementary Fig. 2b**). Because it is difficult to pinpoint the source of leak with a commercial kit, we developed an in-house formulation of NASBA (**Materials and Methods**). Among all the NASBA components (RT, T7 RNAP, RNase H, primers and buffer composed of various salts), we identified RT choice and primer design as important determinants of reaction functional characteristics such as sensitivity, magnitude of fluorescence, and the time at which signal is activated^36–38^. We tested several commercially available RTs: avian myeloblastosis virus (AMV) RT, ProtoScript II RT (a recombinant M-MuLV RT with reduced RNase H activity and increased thermostability), and Moloney murine leukemia virus (M-MuLV) RT (**Fig. 1a**)^39,40^. When NASBA-Cas13a was run using the same primer set and input RNA concentrations, an input RNA concentration-dependent fluorescent signal was observed only with M-MuLV RT and not the other RTs (**Fig. 1b–d**). Notably the leak was diminished in the in-house NASBA reaction compared to the commercial kit results.

To further investigate the impact of RT choice and the presence of any off-target RT products that might interfere with the reaction, we performed NASBA in two steps by staggering the addition of the reaction components (**Materials and Methods**). Native PAGE analysis indicated that a high molecular weight off-target dsDNA product formed during the double-stranding step with AMV RT and ProtoScript II RT (**Supplementary Fig. 3a**, **Supplementary Data 2**). We suspect that this off-target dsDNA product might explain the absence of fluorescent signal with these RTs (**Fig. 1b, c**). In addition, this off-target product was favored in the absence of RNase H (**Supplementary Fig. 3b**), and the same outcome was observed from reactions initiated with CMV as the input RNA (**Supplementary Fig. 3c**). On the other hand, there was an off-target ssDNA product with M-MuLV RT that appears during the first cDNA synthesis step, as well as several off-target dsDNA products in the double-stranding step (**Supplementary Fig. 3a**). These results suggest that the type of off-target NASBA products generated depends on the choice of RT.

To minimize the presence of off-target products observed in **Supplementary Fig. 3**, we explored additional components in the NASBA buffer. We found that including DMSO substantially improved NASBA efficiency (**Supplementary Fig. 4a–d**), presumably by increasing the specificity of the first primer binding^41,42^. Fresh DTT and BSA improved the efficiency as well, though less so than DMSO (**Supplementary Fig. 4e–h**). In summary, NASBA and LbuCas13a-mediated cleavage are compatible in a one-pot format, and the choice of RT and presence of DMSO are important determinants of reaction efficiency.

### Screening of NASBA primer sets

Once we identified an in-house formulation of NASBA that effectively generates activator RNA with little off-target products, we proceeded with designing NASBA primers that target different regions of the SARS-CoV-2 genome. We considered two main factors when designing NASBA primers: (1) the directionality of the primer set and (2) the transcription efficiency of the DNA template generated by reverse transcription. To the first point, we reasoned that a primer set with the T7 promoter incorporated through the reverse primer (for the first cDNA synthesis) should confer more efficient amplification than one with the promoter incorporated through the forward primer. In the former case, a single round of reverse transcription and double strand synthesis generates a DNA template from which an antisense RNA is transcribed (**Supplementary Fig. 1a**), whereas in the latter, additional rounds of DNA synthesis are needed to create a double-stranded T7 promoter and DNA template from which a sense RNA is transcribed (**Supplementary Fig. 1b**). To the second point, we designed several primer sets that result in DNA templates with higher transcription efficiency by incorporating an additional initiating guanine in the reverse primers^43^. In all, there were eight primer sets (**Fig. 1f**). Primer Set 1 targets the gene encoding the S1 spike protein, Primer Set 2 targets the origin of replication, and the remainder sets (3–8) target various regions within SARS-CoV-2 genome that are predicted to be conserved and unstructured^44^. In addition, for Primer Set 1 we designed two different versions targeting the same viral genome region but with opposite primer directionality, so that one amplifies the antisense strand and the other amplifies the sense strand. Of the remaining six sets targeting unstructured regions (Primer Sets 3–8), three are intended to have high transcription efficiency (Primer Sets 6–8).

To test these primer sets, in-house NASBA was run with each set for 3 h using 0 or 20 fM synthetic SARS-CoV-2 genome, and the final RNA products were extracted for urea PAGE analysis (**Fig. 1f**, **Materials and Methods**). We observed three types of outcomes: (1) no expected RNA product (Primer Set 1 – sense; Primer Set 6); (2) the expected RNA product was generated even in the absence of input RNA (Primer Set 1 – antisense; Primer Sets 4 and 5); or (3) the expected RNA product was observed only with input RNA, with little to no off-target products (Primer Sets 2, 3, 7, and 8). In the third category, Primer Sets 2, 7 and 8 had a prominent band of the expected RNA product. This result indicates that primer sets targeting regions with low predicted secondary structure and generating DNA templates designed for efficient T7 transcription produced a large quantity of expected NASBA products only in the presence of input RNA with little to no off-target products.

In summary, we showed that the directionality of the primer set and the sequence of the reverse primer that impacts T7 transcription efficiency can impact NASBA amplification efficiency.

### Optimization of LbuCas13a gRNAs

Once efficient NASBA primer sets were identified, we next sought to investigate gRNA design principles. Previous work engineering Cas13-based detection assays have suggested a number of Cas13 gRNA design principles that could impact assay performance including the number and location of mismatches between gRNA and target RNA, the sequence of the protospacer-flanking site, and the secondary structure of target RNA^23,45–48^. We sought to expand upon this work by screening a panel of LbuCas13a gRNAs targeting each NASBA product. Based on the results in **Fig. 1f**, we focused on the products generated by Primer Sets 2, 7 and 8 for three reasons: (1) minimal product without input RNA, (2) minimal off-target products with input RNA and (3) high-intensity of the expected RNA bands with input RNA.

LbuCas13a complexes with gRNA by recognizing a short hairpin on the 5’ end of the gRNA, followed by a 28-nt spacer that binds to an activator RNA. Previously, it was determined that the structure of the activator RNA, the RNA product in this case generated by NASBA, can impact cleavage efficiency^23^. Taking this into account, we first analyzed predicted secondary structures of each activator RNA and designed two to four gRNAs per activator RNA that target regions of varying secondary structure. Each gRNA is named with two numbers—the first for the primer set and the second for the gRNA variant (e.g., gRNA 2–1 refers to the first gRNA in the series targeting the RNA product generated by Primer Set 2). When in-house NASBA-Cas13a was run with Primer Set 2 and each corresponding gRNA, the reactions with gRNA 2–1 had low leak (signal without input RNA), while those with gRNA 2–2 had high leak (**Supplementary Fig. 5a, b**).

Next, we screened gRNAs targeting the activator RNA generated by Primer Set 7 (**Fig. 2a**, **Supplementary Fig. 5c**). This activator RNA is predicted to form a four-way junction^30,31^, and we designed four gRNAs targeting different regions of the junction: gRNA 7–1 binds to the first and the fourth stems, gRNA 7–2 binds to the third stem, gRNA 7–3 binds to the third and the fourth stems, and gRNA 7-4 binds to the second hairpin and the first stem. In NASBA-Cas13a with these gRNAs, gRNA 7–1 and gRNA 7–2 generated a rapid input RNA-dependent signal and had low leak (**Fig. 2b, c**). Interestingly, gRNA 7–2 generated a signal sooner (and with a steeper slope) than did gRNA 7–1. On the other hand, gRNAs 7–3 and 7–4 performed poorly: gRNA 7–3 conferred low activation and gRNA 7–4 conferred high levels of leak (**Supplementary Fig. 5c**). Interestingly, the two gRNAs with high leak (gRNA 2–2 and gRNA 7–4) contained sequences on their 3’ ends that are complementary to the forward primer, which we suspect could be causing interference with other NASBA reaction components, as it was previously determined that the gRNA 3’ end resides outside of the central channel within the NUC lobe of LbuCas13a^49^. It is unclear from this limited analysis what could cause poor performance of gRNA 7–3.

We also designed three gRNAs to bind the activator RNA generated by Primer Set 8 (**Fig. 2e**). Activator RNA 8 is predicted to form a structure consisting of three hairpins, with the third hairpin including single-stranded regions that are potentially accessible to the gRNA^31^. gRNA 8–1 was designed to bind to the largest single-stranded region in the third hairpin, gRNA 8–2 to a smaller region in the same hairpin and gRNA 8–3 to the second hairpin and the surrounding single-stranded regions. As expected, the fastest signal activation was for NASBA-Cas13a with gRNA 8–1 (**Fig. 2f**). NASBA-Cas13a with gRNA 8–2 targeting a smaller single-stranded region in the same hairpin had much poorer performance, with a low endpoint fluorescent signal and worse sensitivity (**Fig. 2g**). gRNA 8–3 also performed more poorly with a slower activation time than gRNA 8–1 although it is designed to target the smallest hairpin with the lowest number of bps in the activator RNA (**Fig. 2h**).

Despite its low NASBA efficiency, we also screened gRNAs targeting an activator RNA generated by Primer Set 6 to determine whether the observed patterns were similar (**Supplementary Fig. 5e**). Surprisingly, reactions with gRNA 6–1, which is designed to bind a long single-stranded bulge in activator RNA 6, showed very poor performance (**Supplementary Fig. 5f**). This observation contradicts the result seen with gRNA 8–1, which also targets a long single-stranded region within Activator RNA 8 and shows a fast detection time and rapid signal generation (**Fig. 2e, f**). In addition, the gRNAs targeting a stem loop in the activator 6 RNA (gRNA 6–2 and gRNA 6–3) performed better with faster signal activation and higher endpoint fluorescence (**Supplementary Fig. 5g, h**). Finally, we tested a gRNA that binds to an activator RNA generated by Primer Set 3 and observed a poor limit of detection, potentially due to low NASBA efficiency (**Fig. 1f**) and an incorrect hairpin structure for complexing with LbuCas13a (**Supplementary Fig. 5d**).

Overall, the screen identified gRNA candidates that functioned well and could serve as useful starting points for further optimization, and revealed that factors such as local secondary structure of the activator RNA and structure of the gRNA affect NASBA-Cas13a performance.

### Creating a model-driven approach to explore NASBA-Cas13 assay development

Much of diagnostic assay development is done through laborious manual screening of reaction conditions. The advent of new liquid handling instruments provides a way to explore larger spaces of reaction parameters^50^, potentially enabling the training of computational models of reaction mechanisms which could further facilitate exploration of reaction mechanism and optimization.

Here we explored high-throughput screening in the context of the NASBA-Cas13 assay, focusing on varying component concentrations as important parameters that could influence reaction performance (**Fig. 3**). Using Primer Set 8 and gRNA 8-1, we designed a high-throughput screen of NASBA-Cas13 reactions containing different concentrations of input RNA, RT, T7 RNAP, RNase H, and Cas13a-gRNA (**Fig. 3**). NASBA enzyme mix component combinations were dispensed via an Echo liquid-handling robot, added to manually prepared NASBA master mix and LbuCas13a cleavage reaction mix, and characterized through fluorescence measurement by plate reader (**Fig. 3a**, **Supplementary Fig. 6a, Materials and Methods**).

Towards the goal of generating a model to help interpret this high-dimensional data set, we defined qualitative modeling objectives—observations that were representative of all three high-throughput screening experiments—which a formal mathematical representation of this system would need to recapitulate to be useful for guiding interpretation. To this end, we fit Hill functions to the time course trajectories and extracted summary metrics: *n* (Hill coefficient with respect to time), *t*_1/2_ (time to reach half-maximum readout), *F*_0_ (initial fluorescence) and *F*_max_ (maximum fluorescence) (**Fig. 3b**). Distributions of each summary metric across all conditions were used to define the first three modeling objectives. Objective 1: each trajectory had a sigmoidal shape, as indicated by strong agreement (R^2^ ≥ 0.95) between the trajectories and Hill function fits (**Fig. 3c**, **Supplementary Figs. 22a, 24a**). Objective 2: plateaus in readout occurred at various times depending on the condition, as indicated by a range of *t*_1/2_ values (**Fig. 3d**, **Supplementary Figs. 22b, 24b**). Objective 3: the final fluorescent magnitude depends on the reaction condition, as indicated by a range of *F*_max_ values (**Fig. 3e**, **Supplementary Figs. 22c, 24c**). Conditions yielding *F*_max_ ∼0.10 generally corresponded to those lacking input RNA or with low Cas13a-gRNA (e.g., 2.25 nM). Additional modeling objectives were formulated from qualitative observations of the trends in *F*_max_ values as one NASBA enzyme concentration was varied and the other NASBA enzymes were held at mid-level with Cas13a-gRNA at a high level (**Materials and Methods** – **Definition of training data for model development**). Key observations include that readout increased with increasing RT (**Fig. 3f**, **Supplementary Figs. 22d, 24d**, Objective 4), increasing T7 RNAP counterintuitively decreased the readout (**Fig. 3g**, **Supplementary Figs. 22e, 24e**, Objective 5), and increasing RNase H had a non-monotonic effect on the fluorescence readout, in which readout increased from the low to mid-range dose and decreased from the mid-range to the high dose (**Fig. 3h**, **Supplementary Figs. 22f, 24f**, Objective 6). It is unclear for **Data Set 1** whether there is a non-monotonic relationship between RNase concentration and readout (**Supplementary Fig. 22f**) due to the experimental error in the RNase H sweeps, but the trend was clear for **Data Sets 2** (**Supplementary Fig. 23f**) and **3** (**Supplementary Fig. 24f**).

We note that conditions with low Cas13a-gRNA had *F*_max_ values similar to background, indicating no substantial readout (**Supplementary Fig. 8a-c**). We therefore did not incorporate conditions with low input RNA or low Cas13a-gRNA in the modeling objectives because both conditions yielded experimental readout values that are below practical visibility (**Materials and Methods**). The other metrics (*n*, *F*_0_) (**Supplementary Fig. 7**) were not used to define modeling objectives.

Together, the six qualitative objectives defined features of the experimental data that we next aimed to describe using a mathematical model to improve our understanding of the NASBA-Cas13 reaction mechanisms by testing whether a proposed model structure is consistent with these experimental observations. Ordinary differential equation (ODE) models are well-suited to this task, as they describe the continuous, time-dependent evolution of component concentrations such as in genetic systems^51–55^.

### Identifying new putative mechanisms via model development

Our overall approach was to use iterative model formulation and parameter estimation (**Materials and Methods**) to evaluate several candidate models until we arrived at a final model that satisfied all objectives and was in quantitative agreement with each training data set. We selected the conditions representing all three Echo replicates (Data Set 1, 2, and 3) in **Fig. 3f–h**, **Supplementary Figs. 22d-f**, **24d-f** as training data, as these conditions incorporate information on each objective (**Supplementary Fig. 6b** and **Materials and Methods**). Starting with a base case model (**Fig. 4a**), we estimated parameters and inspected whether the simulated values for the optimal parameter set met the modeling objectives (**Materials and Methods**, **Supplementary Note 2**, and **Supplementary Fig. 10a**). With the base model, there was already strong agreement (R^2^ ≥ 0.95) between the simulated trajectories and Hill fits, which indicated that each trajectory had a sigmoidal shape (Objective 1), and the distribution of simulated *t_1/2_* values indicated that plateaus in readout occurred at various times (Objective 2) (**Table 1**, model A, and **Supplementary Figs. 22a-b**, **23a-b**, **24a-b** for the final model). Notably, while the fits for **Data Sets 2 and 3** met Objective 1, they were visually less sigmoidal than the fits to **Data Set 1**. To meet additional modeling objectives and improve the visual fit, we refined our mechanistic descriptions and implemented each change as a new candidate model and repeated the parameter estimation and model evaluation. Next, we describe key observations and refinements encountered while developing a model to fit the training data (Data Set 1, 2, and 3) from Echo replicates.

The first key mechanistic refinement we encountered was the need to describe a loss of Cas13a-gRNA indiscriminate ssRNase activity over time. In the model A simulations, plateaus in readout (*F_max_*) could occur only at the maximum possible value, corresponding to cleavage of all available reporter molecules, which is most evident in the fit to **Data Set 1** (**Supplementary Fig. 12**). To enable a range of simulated *F_max_* values across different reaction conditions, consistent with the experimental data (Objective 3), it was necessary to implement a heuristic function capturing the loss of Cas13a-gRNA indiscriminate ssRNase activity over time (**Table 1**, model B, **Fig. 4b**, right, and **Supplementary Note 1**). The addition of this heuristic successfully resulted in varying *F_max_* values. In addition, the simulated trajectories for RT in model B indicated that increasing RT concentration increased readout, which is in agreement with Objective 4 (**Table 1**, **Supplementary Figs. 15a, 16a, 17a**). However, the fits for **Data Sets 2 and 3** remained visually less sigmoidal than those for **Data Set 1** (**Supplementary Figs. 16 and 17**). The lack of improvement was likely due to higher average experimental error in **Data Sets 2 and 3** and a more complex, clearly non-monotonic relationship between RNase H concentration and readout, compared to **Data Set 1**. The need to account for deactivation of indiscriminate ssRNase activity to yield simulations that are consistent with experimental data suggests a previously unconsidered, potential mechanism affecting the performance of the diagnostic. Additionally, this hypothesis suggests that selecting a different Cas13a-gRNA that deactivates over longer timescales could improve the system.

To match the experimentally observed increase of readout with decrease of T7 RNAP concentration (Objective 5) and non-monotonic readout with varying RNase H concentrations (Objective 6), we revised the mechanistic descriptions of T7 RNAP and RNase H function. Initially, the simulations of models A and B indicated that increasing T7 RNAP concentration should increase readout (violating Objective 5). To achieve an increase in readout with decreasing T7 RNAP concentration (and satisfy Objective 5), it was necessary to implement another heuristic function describing a negative relationship between k_txn, base_ (the rate constant for T7 transcription of activator RNA) and the initial T7 RNAP concentration (**Table 1**, model C, **Fig. 4b**, middle, and **Supplementary Note 1**). This description is plausible, since excess T7 RNAP can inhibit transcription reactions and decrease product yield^56^. Simulations of models A, B, and C also indicated that increasing RNase H concentration should increase readout while experimental results showed non-monotonic behavior (violating Objective 6). To achieve an increase in readout from the low to mid-range concentration and decrease in readout from the mid-range to the high concentration (and satisfy Objective 6), it was necessary to implement a heuristic function with a non-monotonic relationship between k_RHA_ (RNase H activity) and RNase H concentration (**Table 1**, model D, **Fig. 4b**, left, and **Supplementary Note 1**). Although Objective 6 was not satisfied for fits to **Data Set 1**, given the experimental error in the RNase H sweeps, the relationship between RNase concentration and readout is unclear. Therefore, this model is still consistent with fits to **Data Set 1**. To explain the apparent inconsistency in this trend between fits to **Data Set 1** and fits to **Data Sets 2** and **3**, we speculate that technical error could have led to differences in the amount of each reagent dispensed from the liquid handling robot each time the experiment was performed. It is known that dispensing reagents with variable viscosities at small volumes can potentially result in imprecise and/or inaccurate amounts of reagent dispensed that may not be reported by the instrument^57^. The addition of the T7 RNAP and RNase H heuristics resulted in a model that satisfied all modeling objectives for each data set and identified additional hypotheses that could be experimentally tested in future work.

In summary, we arrived at a model structure that qualitatively satisfied each objective for all three data sets when a subset of the data was used for training. The final model (model D) includes three heuristics implementing the loss of Cas13a-gRNA indiscriminate ssRNase activity over time, a negative relationship between the rate of T7 transcription of activator RNA and T7 RNAP concentration, and a non-monotonic relationship between RNase H activity and RNase H concentration (**Table 1**). A detailed schematic of model D is in **Supplementary Fig. 21**, model states are in **Supplementary Table 1**, parameters are in **Supplementary Table 2**, calibrated parameter values are in **Supplementary Table 3**, ODEs are in **Supplementary Table 4**, and comparisons between the experiments and simulations for each data set are in **Supplementary Figs. 22-24**. As noted above (**Materials and Methods**), in this initial modeling effort, we opted not to train the model on conditions without viral RNA or with low Cas13a-gRNA because *F*_max_ values for these conditions were negligible. However, we suspect that agreement between the model and each experimental data set could be further improved by adding a mechanism for background signal (produced in the absence of viral RNA), as experimental maximum readout values for low Cas13a-gRNA conditions resemble background. Altogether, our model development effort yielded a highly explanatory result and identified specific opportunities for future hypothesis-guided experimental and computational investigation.

### Sensitivity analysis of model parameters

To assess whether the estimated parameters were well-constrained across the three data sets, we next performed a parameter sensitivity analysis. We evaluated which parameters had the greatest impact on the simulated time course trajectories (quantified by percent change in *F*_max_ and *t*_1/2_) and overall fit to experimental data (quantified by percent change in MSE) (**Materials and Methods**). Three parameters—k_Cas13_ (the rate of binding of Cas13-gRNA to RNA target), k_loc,deactivation_, and k_scale,deactivation_ (the time and rate of Cas13 deactivation, respectively)—were highly sensitive across all three data sets; varying these parameters resulted in a high percent change in each of the three metrics relative to that incurred when varying the other parameters (**Supplementary Figs. 26, 27**). For the highly sensitive parameters, the magnitude of the percent changes to the performance metrics *F*_max_ and *t*_1/2_ was generally similar. We also compared the calibrated parameter values obtained when using the ODE solver with default versus decreased error tolerances (**Materials and Methods**). Within each data set, the highly sensitive parameter values varied within 1 order of magnitude across solver scenarios (**Supplementary Table 5**). We observed variations in parameter values greater than an order of magnitude (within any data set) only for parameters to which the model is less sensitive, indicating that these parameters are not fully constrained by the data. These observations provide confidence in the numerical methods used to solve the model ODEs.

### Evaluation of final model fits to test data

To assess the predictive capability of the final model with parameters optimized for each data set, we quantified the prediction of each model to test data not included in the training data set. For the final model trained to each replicate training data set, we selected 5 sets of test data, including the data sets used to train the other two models and out-of-sample data on which no model was trained (**Materials and Methods**). Overall, each model produced reasonable fits to the other Echo replicate training data sets (**Supplementary Table 6**). Additionally, these predictions were generally better than the prediction of the out-of-sample data for the same Echo replicate; except for the final model for **Data Set 1** which produced similar fits to each of these test data sets. We suspect that the models performed better on the other Echo replicate training sets compared to the new out-of-sample data, because the other replicate training data are at the same concentration conditions. The fit of each model to Echo replicate 3 out-of-sample data was generally the poorest of the test data fits for a given model, which we attribute to the high average experimental error for Echo replicate 3. Overall, we find that the final model formulation meets the modeling objectives when trained on all three replicates. These models can predict out-of-sample data for the same and new component concentrations with reasonable accuracy, and they perform better for concentrations on which they have been trained. The decrease in prediction accuracy from models trained on **Data Set 1** to **3** is likely due to variation in experimental error in each experiment. Overall, these observations provide helpful guidance as to how future experimental campaigns may best inform model development to align with explanatory and predictive uses of such models. In particular, the variation in component concentrations across high-throughput screens should be more carefully analyzed and incorporated into the model training procedure.

### Limit of detection analysis

Finally, we sought to determine the limit of detection of the assay using the optimized primers and gRNAs (**Supplementary Data 3**). Based on these results, we examined four pairs: (1) Primer Set 2 and gRNA 2–1 (**Fig. 5a**), (2) Primer Set 7 and gRNA 7–1 (**Fig. 5b**), (3) Primer Set 7 and gRNA 7–2 (**Fig. 5c**) and (4) Primer Set 8 and gRNA 8–1 (**Fig. 5d**). Among the pairs, gRNA 7–1, gRNA 7–2 and gRNA 8–1 pairs were sufficiently sensitive for detecting hundreds of attomolar of input RNA. This demonstrates that one-pot formulations of the NASBA-Cas13a reactions have the potential to meet analytical sensitivity requirements of pathogen detection approaches^1,3^.

**Fig. 5.**
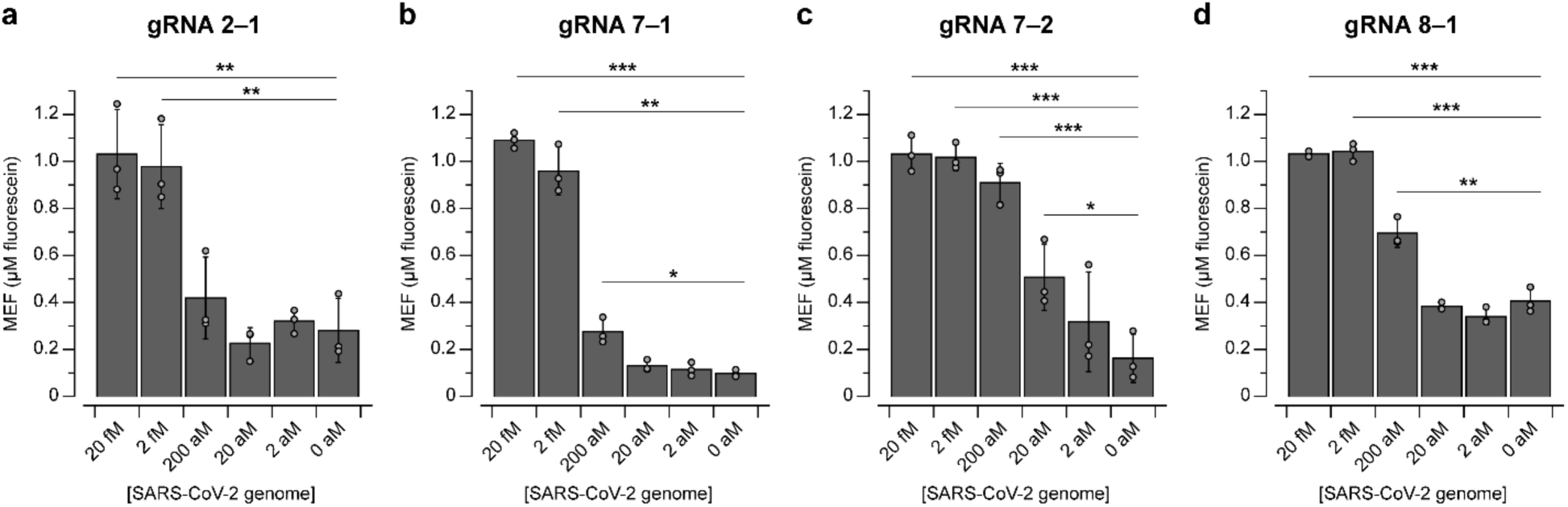
Limit of detection analysis. Fluorescence values at 150 minutes from NASBA-Cas13a with varying concentrations of synthetic SARS-CoV-2 genome using **a**, gRNA 2–1, **b**, gRNA 7–1, **c**, gRNA 7–2, or **d**, gRNA 8–1. Data are n=3 independent biological replicates, each plotted as a point with raw fluorescence values standardized to MEF. Bar height represents the average of the replicates. Error bars indicate the average of the replicates ± standard deviation. Input RNA concentrations for which signal is distinguishable from background (without input RNA) were determined using a two-sided, heteroscedastic Student’s *t*-test. \*\*\**P* < 0.001, \*\**P* = 0.001–0.01, \**P* = 0.01–0.05). *P* values and degrees of freedom are listed in **Supplementary Data 3**.

## DISCUSSION

We developed a test for RNA detection that uses NASBA to amplify a viral RNA and CRISPR-Cas13a activation to cleave a reporter and produce a fluorescent signal. We demonstrated a one-pot isothermal formulation (**Supplementary Fig. 2 and 4**) and screened different reaction components to improve the sensitivity of the test and magnitude of the readout (**Fig. 1** and **Fig. 2**). These investigations led to a test with nucleic acid detection sensitivity around 20-200 aM (**Fig. 5**). High-throughput screening of the NASBA enzyme and input RNA concentration landscape (**Fig. 3**) supported the development of a mechanistic model that explained the effects of component doses on the readout and improved our understanding of the assay (**Fig. 4**). The in-house NASBA formulation was important in facilitating a one-pot isothermal reaction, collecting a data set for model training, and reducing the per-reaction cost. We speculate that it also could enable large-scale test production by eliminating reliance on a commercial kit.

RNA structure was an important consideration when designing primer sets and gRNAs. Among the primers tested, those targeting more structurally flexible regions in the genome led to more efficient amplification (**Fig. 1f**). Similarly, gRNAs targeting less structured regions in the activator RNA generally facilitated a faster readout, especially at low input RNA (**Fig. 2**), although other factors also affected Cas13a activity (**Supplementary Fig. 5**). These factors could arise due to the presence of other components (e.g., the NASBA primers and different buffer compositions that could impact gRNA folding and ribonucleoprotein complexing) that interfere with cleavage reactions.

One limitation of NASBA-Cas13a is that the readout time (1–2 hours) is not as fast as some commercially available antigen tests (15 minutes)^17^ and a RT-LAMP-based nucleic-acid based POC test (30 minutes)^12^. However, we note that NASBA-Cas13a detection is more sensitive than antigen tests and does not require a high incubation temperature of RT-LAMP (60∼65°C). The system also has not yet been validated on patient samples, which potentially contain reaction inhibitors. Our focus was on investigating the impact of various design choices on effective nucleic acid detection. Field deployment would be a logical step to pursue in subsequent work focusing on translational deployment.

An innovation that went into optimizing NASBA-Cas13a was the use of ODE modeling for this system. Through iterative model development, we identified previously unconsidered mechanisms that led to lower-than-expected readout. We found that it was necessary to invoke mechanisms for Cas13a deactivation, an inverse relationship between T7 RNAP concentration and the readout within the relevant local concentrations, and a non-monotonic relationship between RNase H concentration and the readout within the relevant local concentration regime (**Table 1**). The identification of these relationships demonstrates the power of explanatory computational modeling to translate results from an empirical scan into specific hypotheses that could be pursued by experimental investigation to build mechanistic understanding. Future work could include testing targeted interventions to mitigate these limitations or predicting interventions that could improve certain performance metrics (considerations are listed in **Supplementary Note 4**). The model development process used in this study is an extension of the GAMES workflow^28^ and the first instance in which the workflow was used to describe experimental observations. We anticipate that this approach may be extensible to other molecular diagnostic tests.

While the results from the high-throughput and manual reaction data were not reconcilable, this is an area of future work that can link our understanding of the two distinct methods and our modeling work. We decided to move forward with the high-throughput data as a proof-of-concept that such a data set could be used to train a model of a complex molecular system. Despite the discrepancies observed between the two setups, the model trained with the high-throughput assay was useful for identifying mechanisms that were not elucidated from the manual reaction setup. We expect that with further refinement, some of the observed experimental noise from these platforms could be reduced so that they can be used to efficiently test more reaction conditions. Above all, found value in using an ODE modeling workflow to describe CRISPR-Cas-based diagnostic assays, which to the best of our knowledge has not been previously done.

There is a growing interest in CRISPR-Cas–based nucleic acid detection techniques. This study contributes to the growing body of POC tests and provides a starting point for model-driven characterization and engineering of CRISPR-based POC tests. Uniting systematic manual experimental characterization, high-throughput screening experiments and rigorous mechanistic mathematical modeling will set the stage for model-driven experimental design of *in vitro* systems.

## DATA AVAILABILITY

All data are available as **Supplementary Data** deposited in Northwestern University Research and Data Depository (doi: 10.21985/n2-bn16-9b08).

## CODE AVAILABILITY

Code is provided at https://github.com/leonardlab/COVID_Dx_GAMES under an open-source license. Simulation outputs are provided in the Reported_results directory within the COVID_Dx_GAMES repository. We refactored the code in the GAMES v2.0.0 framework (https://github.com/leonardlab/GAMES) to improve accessibility, and this version of the code is available at https://github.com/leonardlab/COVID_Dx_GAMES2.

## FUNDING

This work was supported in part by the National Science Foundation Graduate Research Fellowship Program (DGE-1842165: K.E.D. and G.A.R), the National Institute of Biomedical Imaging and Bioengineering of the NIH (1R01EB026510, 5R01EB026510-06 to J.N.L.), the National Science Foundation (2028651 to J.B.L. and M.J.C; 2310382 to J. B. L.), and the AFOSR DURIP program (FA9550-23-1-0420). J.B.L. was also supported by a John Simon Guggenheim Memorial Foundation Fellowship. This work was supported in part by the Bill & Melinda Gates Foundation [INV-038694]. Under the grant conditions of the Foundation, a Creative Commons Attribution 4.0 Generic License has already been assigned to the Author Accepted Manuscript version that might arise from this submission.

## CONFLICT OF INTEREST DISCLOSURE

K.K.A. M. C. J. & J.B.L. are founders and have financial interest in Stemloop, Inc, and these interests are reviewed and managed by Northwestern University and Stanford University in accordance with their conflict-of-interest policies. All other authors declare no conflicts of interest.

## ACKNOLWEDGEMENTS

We thank Andrea Thompson (NU), Charlotte Knopp (NU), and Lauren Clark (NU) for managing reagents and equipment; Dr. Sergii Pshenychnyi (NU Recombinant Protein Production Core) for assistance in protein purification; Walter Thavarajah (NU) for initial validation studies; Roman Iwasaki (University of Colorado, Boulder) for providing LwCas13a, which helped in initial studies; Jessica S. Yu (UW) for feedback on figure design for modeling work.

## AUTHOR CONTRIBUTIONS

Conceptualization: JKJ, KSD, KED, JJM, JG, SS, MDC, AD, MSV, KS, GAR, KKA, NB, JNL, MCJ, NMM, JBL

Data curation: JKJ, JSD

Formal analysis: JKJ, KSD, KED, JJM, JG, SS, NB, JNL, MCJ, NMM, JBL

Funding acquisition: JNL, MCJ, NMM, JBL

Investigation: JKJ, KSD, KED, JJM, JG, SS, MDC, AD, MSV, KS, GAR, KKA, NB, JNL, MCJ, NMM, JBL

Project administration: NMM, JBL Software: KSD, KED, JJM, JG, SS

Validation: JKJ, KSD, KED Visualization: JKJ, KSD, KED

Writing – original draft: JKJ, KSD, KED, JJM, JG, SS, JNL, NMM, JBL

Writing – review and editing: JKJ, KSD, KED, JJM, JG, SS, MDC, AD, MSV, KS, GAR, KKA, NB, JNL, MCJ, NMM, JBL

## Notes

https://dx.doi.org/10.21985/n2-bn16-9b08

https://github.com/leonardlab/COVID_Dx_GAMES

https://github.com/leonardlab/COVID_Dx_GAMES2

## REFERENCES

1. Gootenberg, J. S. et al. Nucleic acid detection with CRISPR-Cas13a/C2c2. Science 356, 438–442 (2017).

2. Chen, J. S. et al. CRISPR-Cas12a target binding unleashes indiscriminate single-stranded DNase activity. Science 360, 436–439 (2018).

3. Joung, J. et al. Detection of SARS-CoV-2 with SHERLOCK One-Pot Testing. N. Engl. J. Med. 383, 1492–1494 (2020).

4. Gootenberg, J. S. et al. Multiplexed and portable nucleic acid detection platform with Cas13, Cas12a, and Csm6. Science 360, 439–444 (2018).

5. Harrington, L. B. et al. Programmed DNA destruction by miniature CRISPR-Cas14 enzymes. Science 362, 839–842 (2018).

6. Taipale, J., Romer, P. & Linnarsson, S. Population-scale testing can suppress the spread of COVID-19. bioRxiv (2020) doi:10.1101/2020.04.27.20078329.

7. Akst, J. RNA extraction kits for COVID-19 tests are in short supply in US. The Scientist Magazine® https://www.the-scientist.com/rna-extraction-kits-for-covid-19-tests-are-in-short-supply-in-us-67250 (2020).

8. More coronavirus tests will be available next month, Fauci says, as U.S. struggles with shortage. The Washington Post (2021).

9. Notomi, T. et al. Loop-mediated isothermal amplification of DNA. Nucleic Acids Res. 28, E63 (2000).

10. Piepenburg, O., Williams, C. H., Stemple, D. L. & Armes, N. A. DNA detection using recombination proteins. PLoS Biol. 4, e204 (2006).

11. Broughton, J. P. et al. CRISPR–Cas12-based detection of SARS-CoV-2. Nat Biotechnol 38, 870–874 (2020).

12. Amaral, C. et al. A molecular test based on RT-LAMP for rapid, sensitive and inexpensive colorimetric detection of SARS-CoV-2 in clinical samples. Sci. Rep. 11, 16430 (2021).

13. Sun, Y. et al. One-tube SARS-CoV-2 detection platform based on RT-RPA and CRISPR/Cas12a. J. Transl. Med. 19, 74 (2021).

14. Ludwig, K. U. et al. LAMP-Seq enables sensitive, multiplexed COVID-19 diagnostics using molecular barcoding. Nat. Biotechnol. 39, 1556–1562 (2021).

15. Bloom, J. S. et al. Massively scaled-up testing for SARS-CoV-2 RNA via next-generation sequencing of pooled and barcoded nasal and saliva samples. Nat Biomed Eng 5, 657–665 (2021).

16. Fozouni, P. et al. Amplification-free detection of SARS-CoV-2 with CRISPR-Cas13a and mobile phone microscopy. Cell 184, 323–333.e9 (2021).

17. Gonzalez, F. O. & Moore, N. Performance of the BinaxNOW COVID-19 Antigen Rapid Diagnostic Test for the Detection of SARS-CoV-2. Preprint at 10.21203/rs.3.rs-1206340/v1 (2021).

18. Cerutti, F. et al. Urgent need of rapid tests for SARS CoV-2 antigen detection: Evaluation of the SD-Biosensor antigen test for SARS-CoV-2. J. Clin. Virol. 132, 104654 (2020).

19. Porte, L. et al. Evaluation of a novel antigen-based rapid detection test for the diagnosis of SARS-CoV-2 in respiratory samples. Int. J. Infect. Dis. 99, 328–333 (2020).

20. El Wahed, A. A., et al. Suitcase Lab for Rapid Detection of SARS-CoV-2 Based on Recombinase Polymerase Amplification Assay. Anal. Chem. 93, 2627–2634 (2021).

21. Minimally instrumented SHERLOCK (miSHERLOCK) for CRISPR-based point-of-care diagnosis of SARS-CoV-2 and emerging variants. *SCIENCE ADVANCES* 12 (2021).

22. Compton, J. Nucleic acid sequence-based amplification. Nature 350, 91–92 (1991).

23. Abudayyeh, O. O. et al. C2c2 is a single-component programmable RNA-guided RNA-targeting CRISPR effector. Science 353, aaf5573 (2016).

24. Tambe, A., East-Seletsky, A., Knott, G. J., Doudna, J. A. & O’Connell, M. R. RNA Binding and HEPN-Nuclease Activation Are Decoupled in CRISPR-Cas13a. Cell Rep. 24, 1025– 1036 (2018).

25. Agrawal, S. et al. Rapid, point-of-care molecular diagnostics with Cas13. medRxiv (2021) doi:10.1101/2020.12.14.20247874.

26. Oliveira, B. B., Veigas, B. & Baptista, P. V. Isothermal amplification of nucleic acids: The race for the next “gold standard.” Front. Sens. 2, (2021).

27. Kahl, L. et al. Opening options for material transfer. Nat. Biotechnol. 36, 923–927 (2018).

28. Dray, K. E., Muldoon, J. J., Mangan, N. M., Bagheri, N. & Leonard, J. N. GAMES: A Dynamic Model Development Workflow for Rigorous Characterization of Synthetic Genetic Systems. ACS Synth. Biol. acssynbio.1c00528 (2022) doi:10.1021/acssynbio.1c00528.

29. East-Seletsky, A. et al. Two distinct RNase activities of CRISPR-C2c2 enable guide-RNA processing and RNA detection. Nature 538, 270–273 (2016).

30. Reuter, J. S. & Mathews, D. H. RNAstructure: software for RNA secondary structure prediction and analysis. BMC Bioinformatics 11, 129 (2010).

31. Zadeh, J. N. et al. NUPACK: Analysis and design of nucleic acid systems. J. Comput. Chem. 32, 170–173 (2011).

32. Virtanen, P. et al. SciPy 1.0: fundamental algorithms for scientific computing in Python. Nat. Methods 17, 261–272 (2020).

33. Herman, J. & Usher, W. SALib: An open-source Python library for Sensitivity Analysis. J. Open Source Softw. 2, 97 (2017).

34. Newville, M., Stensitzki, T., Allen, D. B. & Ingargiola, A. LMFIT: Non-Linear Least-Square Minimization and Curve-Fitting for Python. (Zenodo, 2014). doi:10.5281/ZENODO.11813.

35. Verosloff, M., Chappell, J., Perry, K. L., Thompson, J. R. & Lucks, J. B. PLANT-Dx: A Molecular Diagnostic for Point-of-Use Detection of Plant Pathogens. ACS Synth. Biol. 8, 902–905 (2019).

36. Wu, Q. et al. INSIGHT: a population scale COVID-19 testing strategy combining point-of-care diagnosis with centralised high-throughput sequencing. bioRxiv 2020.06.01.127019 (2020) doi:10.1101/2020.06.01.127019.

37. McCalla, S. E. et al. A simple method for amplifying RNA targets (SMART). J. Mol. Diagn. 14, 328–335 (2012).

38. Luebke, K. J. Prioritized selection of oligodeoxyribonucleotide probes for efficient hybridization to RNA transcripts. Nucleic Acids Research 31, 750–758 (2003).

39. Kotewicz, M. L., Sampson, C. M., D’Alessio, J. M. & Gerard, G. F. Isolation of cloned Moloney murine leukemia virus reverse transcriptase lacking ribonuclease H activity. Nucleic Acids Res. 16, 265–277 (1988).

40. Lim, D. et al. Crystal structure of the moloney murine leukemia virus RNase H domain. J. Virol. 80, 8379–8389 (2006).

41. Jensen, M. A., Fukushima, M. & Davis, R. W. DMSO and betaine greatly improve amplification of GC-rich constructs in de novo synthesis. PLoS One 5, e11024 (2010).

42. Varadharajan, B. & Parani, M. DMSO and betaine significantly enhance the PCR amplification of ITS2 DNA barcodes from plants. Genome 64, 165–171 (2021).

43. Conrad, T., Plumbom, I., Alcobendas, M., Vidal, R. & Sauer, S. Maximizing transcription of nucleic acids with efficient T7 promoters. Commun Biol 3, 439 (2020).

44. Rangan, R. et al. RNA genome conservation and secondary structure in SARS-CoV-2 and SARS-related viruses: a first look. RNA 26, 937–959 (2020).

45. Wessels, H.-H. et al. Massively parallel Cas13 screens reveal principles for guide RNA design. Nat. Biotechnol. 38, 722–727 (2020).

46. Mantena, S. et al. Model-directed generation of CRISPR-Cas13a guide RNAs designs artificial sequences that improve nucleic acid detection. bioRxiv (2023) doi:10.1101/2023.09.20.557569.

47. Meeske, A. J. & Marraffini, L. A. RNA Guide Complementarity Prevents Self-Targeting in Type VI CRISPR Systems. Mol. Cell 71, 791–801.e3 (2018).

48. Bandaru, S. et al. Structure-based design of gRNA for Cas13. Sci. Rep. 10, 11610 (2020).

49. Liu, L. et al. The Molecular Architecture for RNA-Guided RNA Cleavage by Cas13a. Cell 170, 714–726.e10 (2017).

50. Borkowski, O. et al. Large scale active-learning-guided exploration for in vitro protein production optimization. Nat. Commun. 11, 1872 (2020).

51. Donahue, P. S. et al. The COMET toolkit for composing customizable genetic programs in mammalian cells. Nat. Commun. 11, 779 (2020).

52. Gómez-Schiavon, M., Dods, G., El-Samad, H. & Ng, A. H. Multidimensional Characterization of Parts Enhances Modeling Accuracy in Genetic Circuits. ACS Synth. Biol. 9, 2917–2926 (2020).

53. Agrawal, D. K. et al. Mathematical Modeling of RNA-Based Architectures for Closed Loop Control of Gene Expression. ACS Synth. Biol. 7, 1219–1228 (2018).

54. Frei, T. et al. Characterization and mitigation of gene expression burden in mammalian cells. Nat. Commun. 11, 4641 (2020).

55. Lillacci, G., Benenson, Y. & Khammash, M. Synthetic control systems for high performance gene expression in mammalian cells. Nucleic Acids Res. 46, 9855–9863 (2018).

56. Milligan, J. F. & Uhlenbeck, O. C. [5] Synthesis of small RNAs using T7 RNA polymerase. In RNA Processing Part A: General Methods vol. 180 51–62 (Academic Press, 1989).

57. Rhea, K. A. et al. Variability in cell-free expression reactions can impact qualitative genetic circuit characterization. Synth. Biol. 7, ysac011 (2022).

