## Supplementary Information for "Developing, characterizing and modeling CRISPR-based point-of-use pathogen diagnostics"

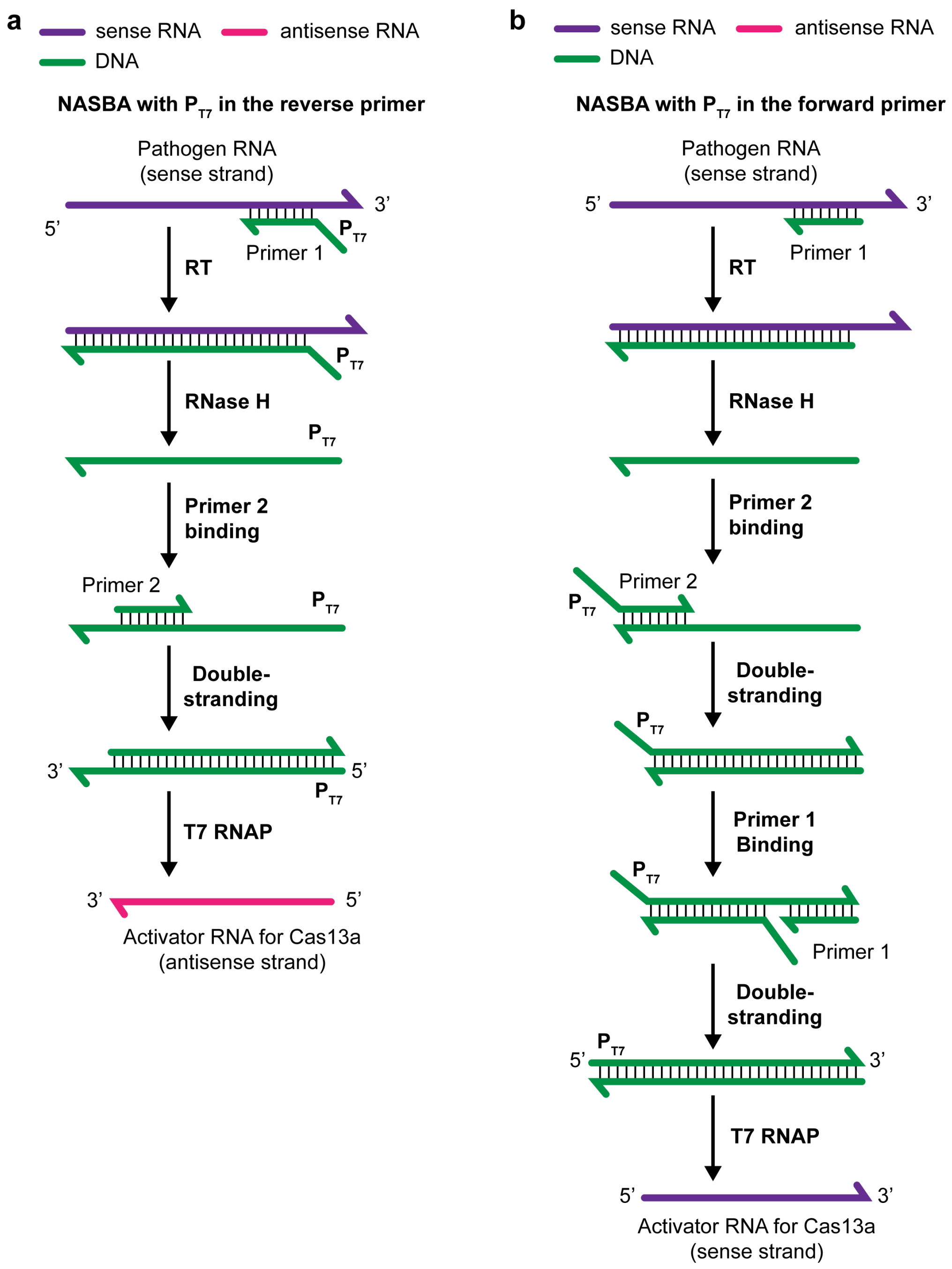


**Supplementary Fig. 1 | NASBA with different primer directionalities. a**, Diagram of NASBA using a primer set in which the T7 promoter is incorporated through the reverse primer, i.e., the primer for the first cDNA synthesis. This primer set generates antisense RNA (complementary to the input RNA). **b**, Diagram of NASBA using a primer set in which the T7 promoter is incorporated through the forward primer. This primer set amplifies the same sequence as the input RNA, and there is an extra round of DNA synthesis required to double strand the T7 promoter prior to generating the RNA product.

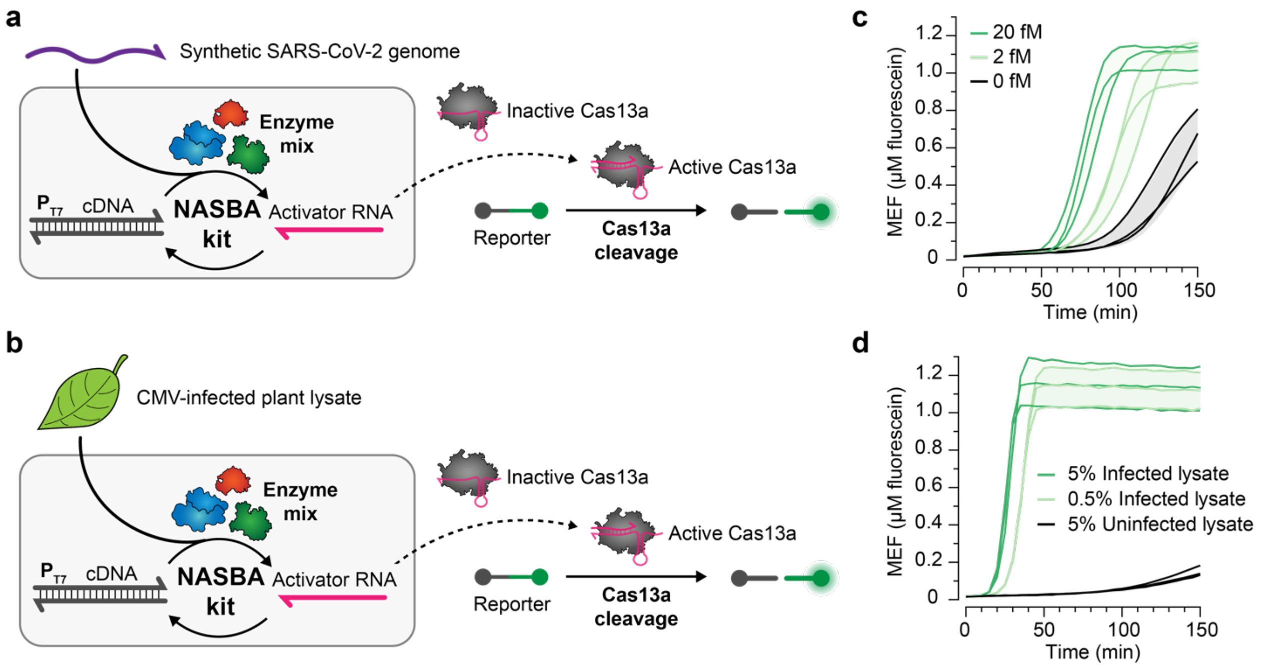


**Supplementary Fig. 2 | NASBA and CRISPR-Cas13a cleavage can be performed in a one-pot isothermal reaction.** One-pot NASBA-Cas13a was performed with a commercial NASBA kit using **a**, synthetic SARS-CoV-2 genome or **b**, plant lysate enriched with cucumber mosaic virus (CMV) as RNA input. The kit generates activator RNA. Upon sensing this RNA, LbuCas13a indiscriminate ssRNase activity is activated, and LbuCas13a cleaves a reporter, producing a fluorescent signal. **c**, Fluorescence kinetics of SARS-CoV-2–sensing NASBA-Cas13a (primer set 8 – gRNA 1) initiated by 0, 2 or 20 fM synthetic SARS-CoV-2 genome. **d**, Fluorescence kinetics of the CMV–sensing NASBA-Cas13a initiated by 0.5% or 5% v/v infected plant lysate or 5% v/v uninfected plant lysate. Data shown are for n=3 independent biological replicates, each plotted as a line with raw fluorescence standardized to MEF (μM fluorescein). Shading indicates the average of the replicates ± standard deviation. Sequences of primers and gRNAs are listed in **Supplementary Data 1**.

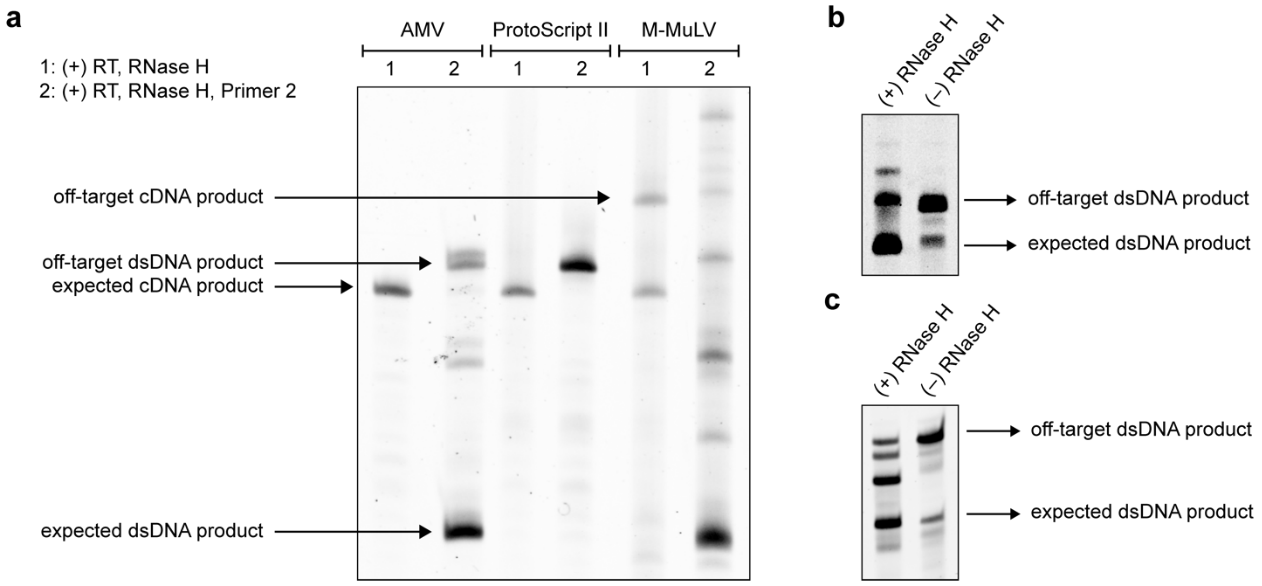


**Supplementary Fig. 3 | Sequential NASBA reveals various off-target products. a,** Sequential NASBA using RTs tested in **Fig. 1** generates different off-target products at different steps of the reaction (Sequential NASBA in **Materials and Methods**). Testing the effect of RNase H on off-target products. Reactions were performed sequentially with 0.5 U/µL AMV RT and initiated either by **b**, SARS-CoV-2 input RNA fragment targeted by primer set 5 or by **c**, CMV input RNA fragment. Data in **a**–**c** are a representative of n=3 independent biological replicates. Uncropped, unprocessed gel images in **a**–**c** are available as **Supplementary Data 2**.

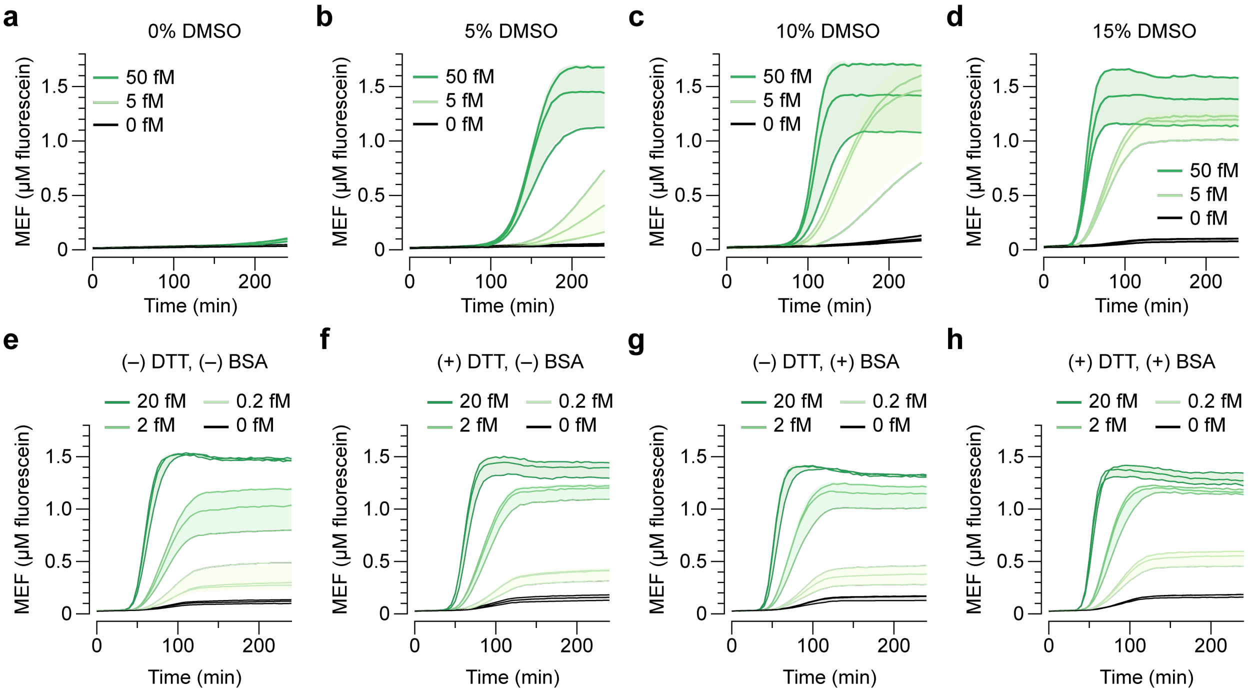


**Supplementary Fig. 4 | Optimization of in-house NASBA-Cas13a.** DMSO improves NASBA efficiency. Fluorescence kinetics from NASBA-Cas13a (primer set 8 – gRNA 1) initiated by 0, 5 or 50 fM synthetic SARS-CoV-2 genome in the presence of varying concentrations of DMSO: **a**, 0%, **b**, 5%, **c**, 10% and **d**, 15%. Adding fresh DTT and BSA with 15% DMSO increases the final fluorescence magnitude. Fluorescence kinetics from NASBA-Cas13a (primer set 8 – gRNA 1) initiated with 0, 0.2, 2 or 20 fM synthetic SARS-CoV-2 genome **e**, without fresh DTT or BSA, **f**, with 5 mM fresh DTT, **g**, with 0.1 µg/µL BSA or **h**, with 5 mM fresh DTT and 0.1 µg/µL BSA. Data shown are for n=3 independent biological replicates, each plotted as a line with raw fluorescence standardized to MEF (µM fluorescein). Shading indicates the average of the replicates ± standard deviation. Sequences of primers and gRNAs are listed in **Supplementary Data 1**.

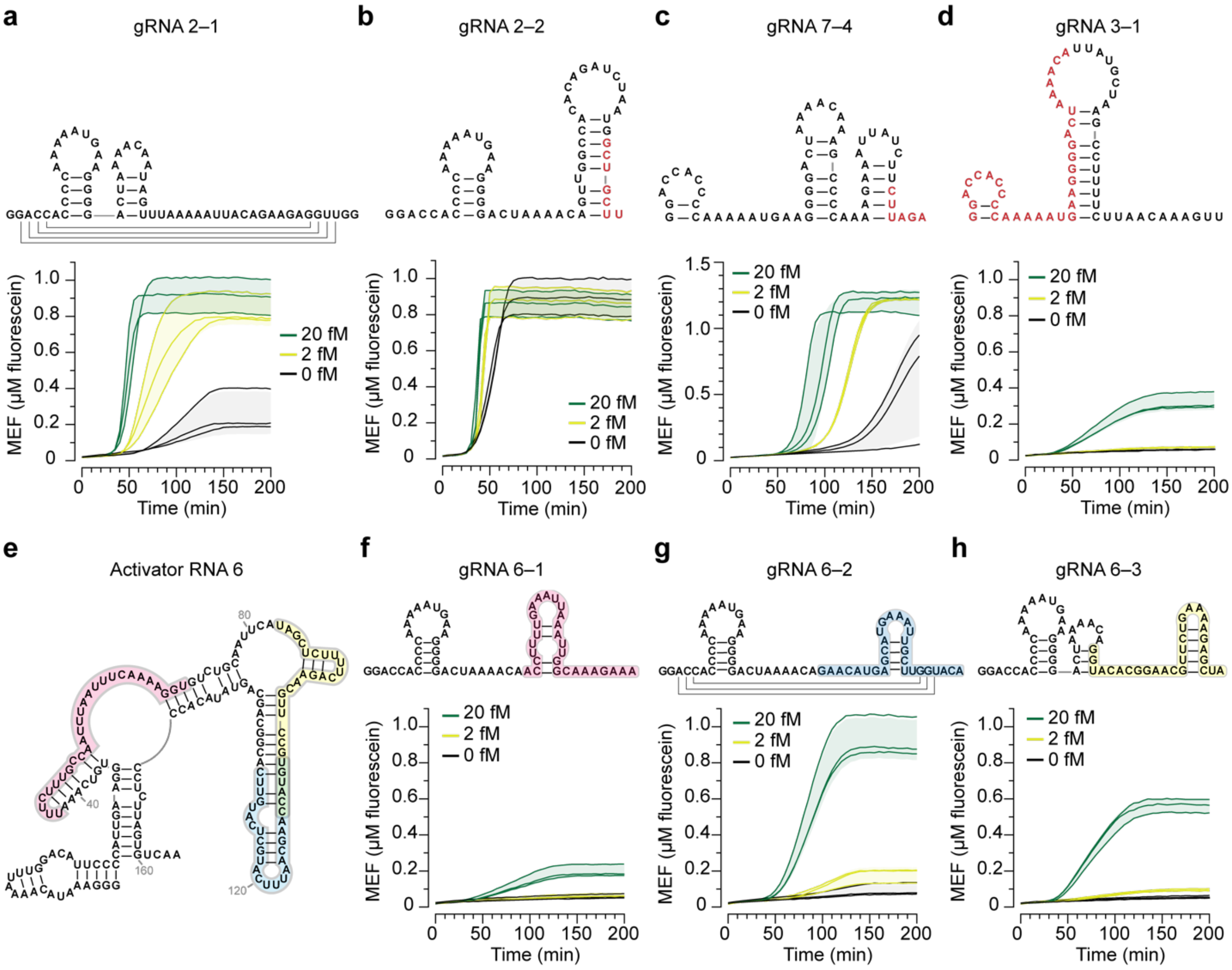

**Supplementary Fig. 5 | Screening of guide RNAs and activator RNAs.** Fluorescence kinetics for NASBA-Cas13a with varying concentrations of synthetic SARS-CoV-2 genome using **a**, gRNA 2–1, **b**, gRNA 2–2, **c**, gRNA 7–4 or **d**, gRNA 3–1. A predicted secondary structure of each gRNA including the constant region and the spacer sequence is depicted. The spacer sequence in gRNA 2–2 and gRNA 7–4 overlaps with the corresponding NASBA primer binding site (red). gRNA 3–1 is not predicted to form the necessary hairpin structure (red) required for complexing with LbuCas13a, which potentially contributes to the observed low cleavage efficiency. **e**, A predicted secondary structure of activator RNA 6. The region targeted by each gRNA is shaded. Fluorescence kinetics for NASBA-Cas13a with varying concentrations of synthetic SARS-CoV-2 genome using **f**, gRNA 6–1, **g**, gRNA 6–2 or **h**, gRNA 6–3 with predicted secondary structure of each gRNA above the kinetic traces. Lines on gRNA 2–1 (**a**) and gRNA 6–2 (**g**) indicate additional predicted long-range interactions. Data shown are for n=3 independent biological replicates, each plotted as a line with raw fluorescence standardized to MEF (μM fluorescein). Shading indicates the average of the replicates ± standard deviation.

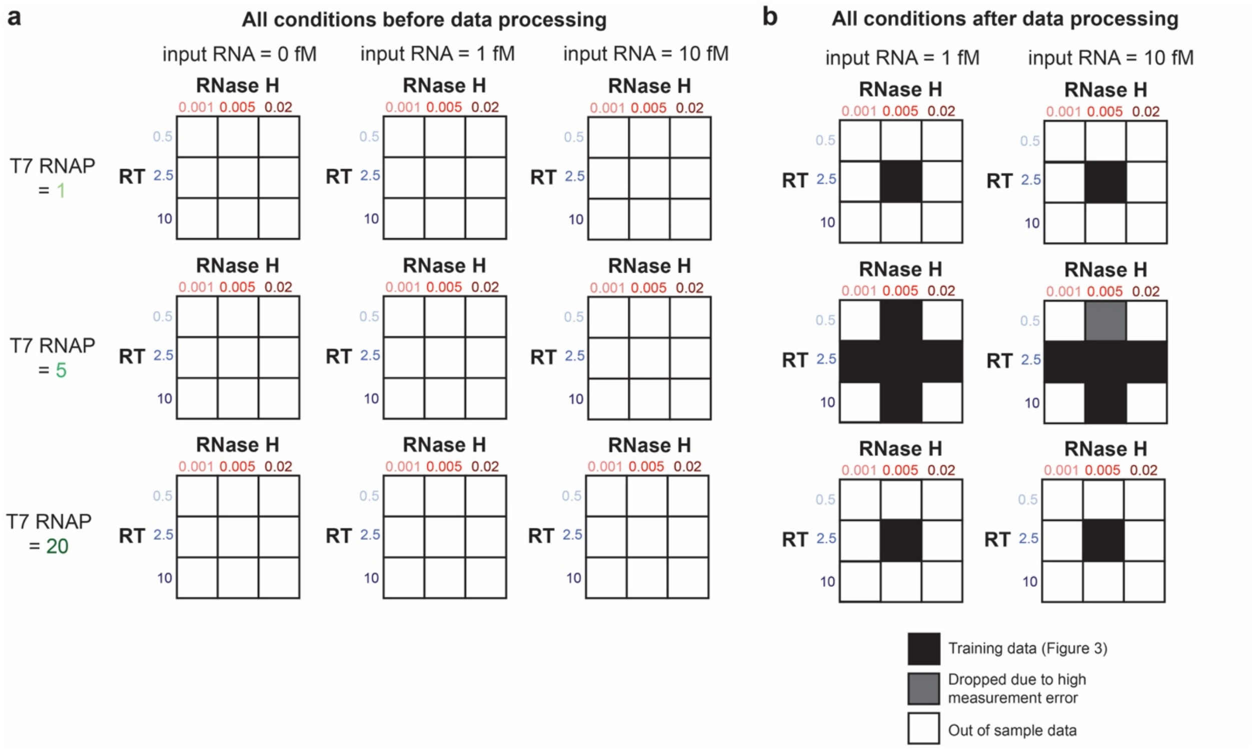


**Supplementary Fig. 6 | Experimental conditions before and after data processing. a,** Full experimental data set before pre-processing**.** The conditions shown here were run with both low (2.25 nM) and high Cas13a-gRNA (45 nM). Numbers for T7 RNAP, RNase H and RT are in units of U/μL. **b,** The model did not aim to describe low Cas13a-gRNA conditions or conditions with 0 input RNA, so these conditions were withheld. An additional condition (dark gray) was withheld from the training data for data set 1 due to high measurement error, but this condition was used in the training data for data sets 2 and 3 (**Supplementary Fig. 9**). For each data set, the out of sample data was defined as the conditions after pre-processing that were not in the training data.

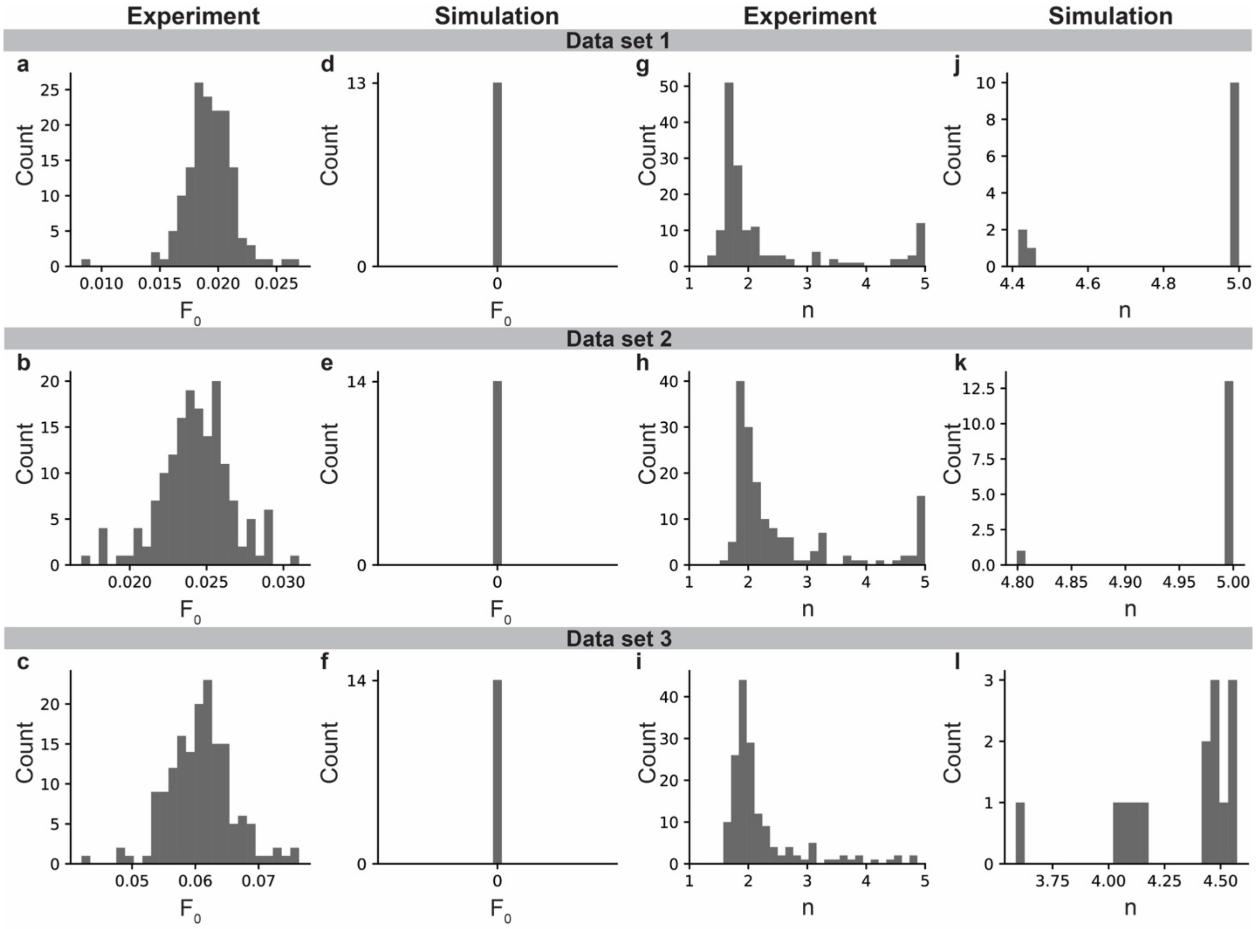


**Supplementary Fig. 7 | Distribution of summary metrics $F_0$ and *n*.** Histogram of $F_0$ values across **a-c**, all conditions in experimental Data Sets 1, 2, and 3, respectively and **d-f**, the subset of conditions used for parameter estimation in simulated Data Sets 1, 2, and 3, respectively. It is not intended for the model to include a mechanism for background signal (readout in the absence of input RNA). Histogram of *n* values across **g-i**, all conditions in experimental Data Sets 1, 2, and 3, respectively and **j-l**, the subset of conditions used for parameter estimation in simulated Data Sets 1, 2, and 3, respectively.

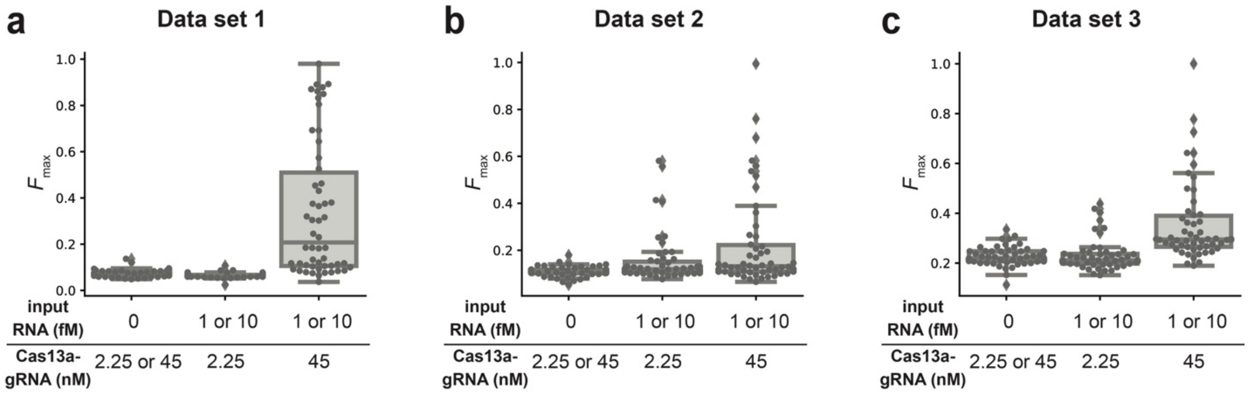


**Supplementary Fig. 8 | When Cas13a-gRNA is relatively low, the readout is similar to background level.** Fitted $F_{max}$ values across time courses with shared conditions. Data points represent fitted $F_{max}$ values, the box extends from the Q1 to Q3 quartile values, the line represents the median, and the whiskers denote the range of the data. **a-c**, Experimental data for Data Sets 1, 2, and 3, respectively. The left box is for conditions without input RNA and includes conditions with low and high Cas13a-gRNA (2.25 and 45 nM). The middle box is for conditions with input RNA (1 and 10 fM) and low Cas13a-gRNA. The right box is for conditions with input RNA and high Cas13a-gRNA. Conditions with low Cas13a-gRNA have $F_{max}$ values generally indistinguishable from without input RNA.

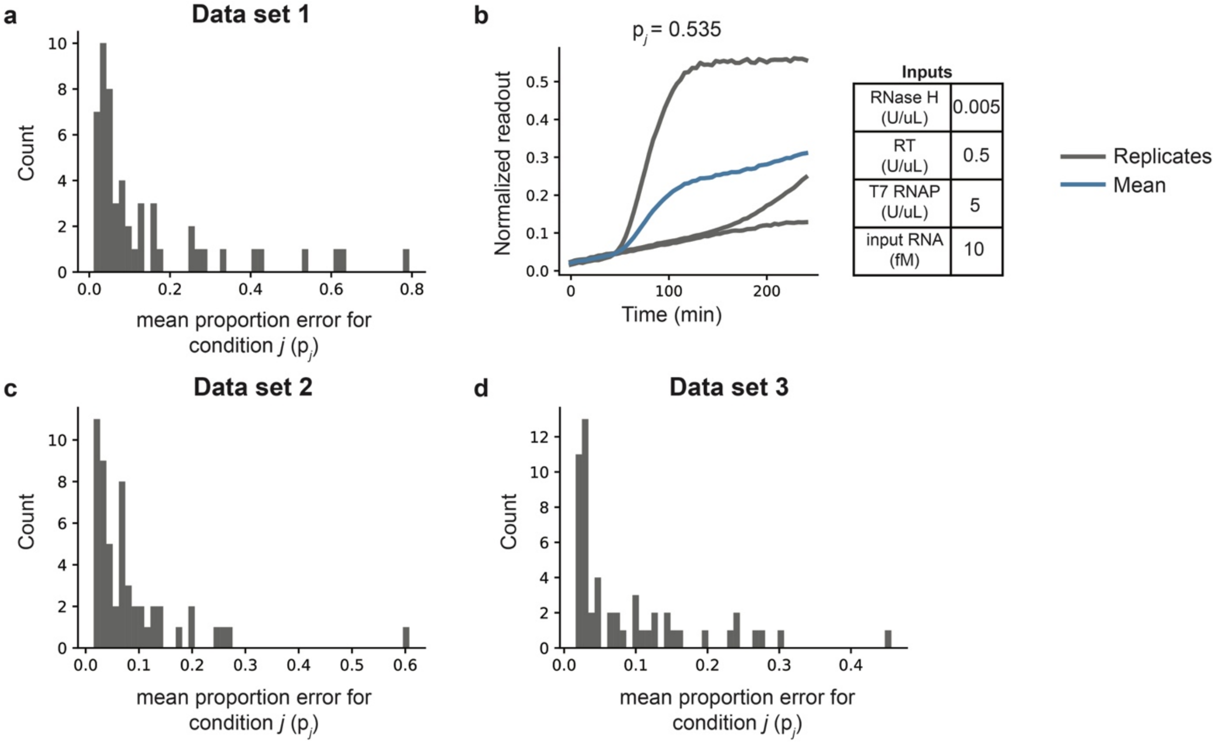


**Supplementary Fig. 9 | Pre-processing of training data to remove conditions with relatively high measurement error. a**, The mean proportion error, p, was calculated for each condition, *j* in data set 1 (**Materials and Methods**). **b**, For the condition in data set 1 with $p_j > 0.3$ within the subset of conditions before pre-processing (**Materials and Methods**), normalized readout trajectories for each replicate are shown (gray), along with the normalized mean readout trajectory across all 3 replicates (blue). This condition had one replicate for which the readout remained near zero throughout the time course, in contrast to the other replicates in the condition. **c-d**, For data sets 2 and 3, the mean proportion error, p, was calculated for each condition, *j*. The condition with $p_j > 0.3$ in each data set was not in the subset of conditions before pre-processing (**Materials and Methods**), so these conditions were not analyzed further or removed from training data.

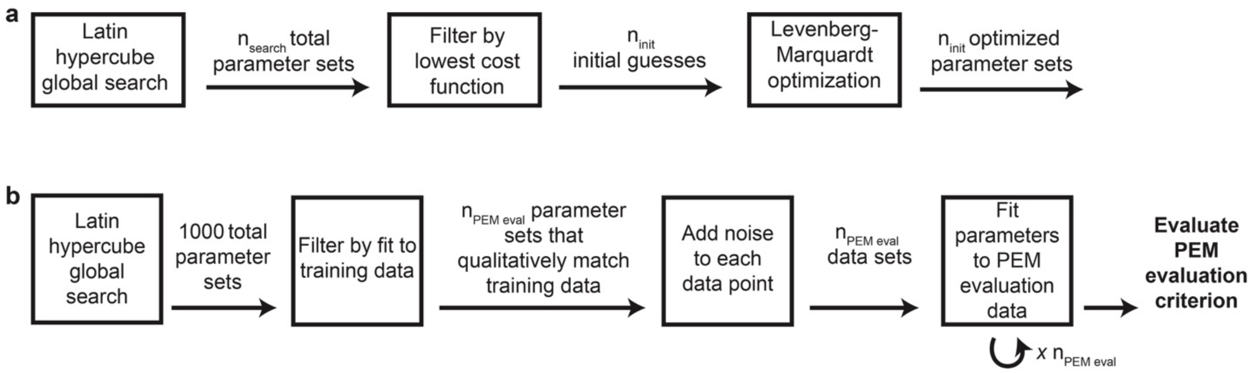


**Supplementary Fig. 10 | Parameter estimation method workflow and evaluation. a**, Parameters were estimated using a multi-start optimization strategy. First, a Latin hypercube global search with $n_{search}$ parameter sets was used to sample the parameter space. The top $n_{init}$ parameter sets (with the lowest cost function values) were each used to initialize independent optimization runs using the Levenberg-Marquardt algorithm. The optimized parameter set with the lowest cost function was defined as the calibrated parameter set. **b**, The parameter estimation method (PEM) was evaluated for each model. First, a Latin hypercube global search was performed, and the results were filtered based on fit to the training data. The top $n_{PEM\ eval}$ parameter sets were used to generate PEM evaluation training data by using each parameter set to simulate the training data and adding noise to each data point. Next, parameters were estimated using each PEM evaluation data set, and the results were analyzed.

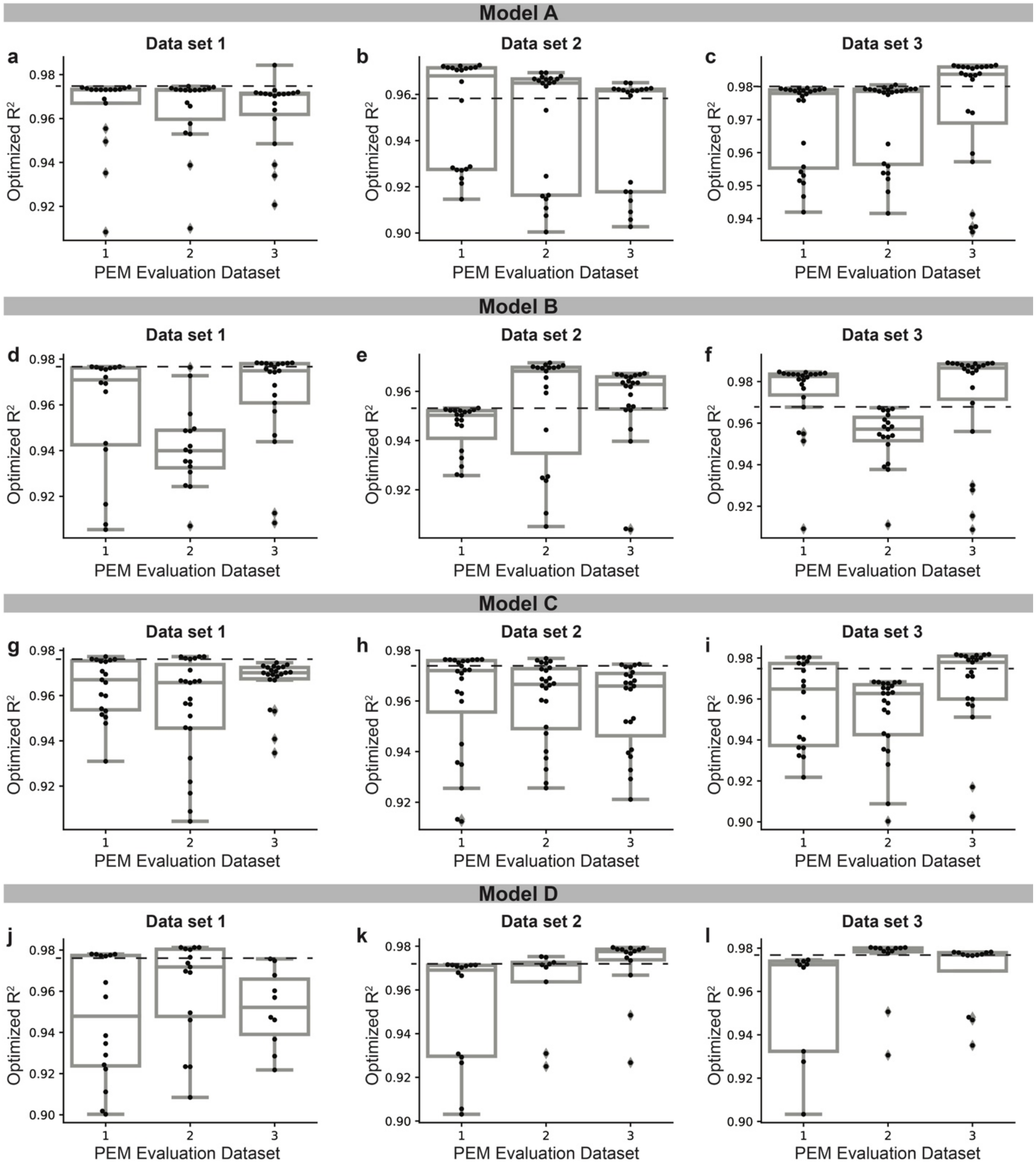


**Supplementary Fig. 11 | Parameter estimation evaluation results. a-c**, PEM evaluation results for model A for Data Sets 1, 2, and 3, respectively. **d-f**, PEM evaluation results for model B for Data Sets 1, 2, and 3, respectively. **g-i**, PEM evaluation results for model C for Data Sets 1, 2, and 3, respectively. **j-l**, PEM evaluation results for model D for Data Sets 1, 2, and 3, respectively. In each case, $n_{search}$ = 5000 and $n_{init}$ = 24. Note that the PEM evaluation criterion is not exactly met for model C, Data Set 1, PEM evaluation Data Set 3; model C, Data Set 3, PEM evaluation Data Set 2; and model D, Data Set 3, PEM evaluation Data Set 1. In each case, the PEM evaluation data set is just under (<0.005) the PEM evaluation criterion. This is a minor inconsistency that is unlikely to affect downstream parameter estimation results.

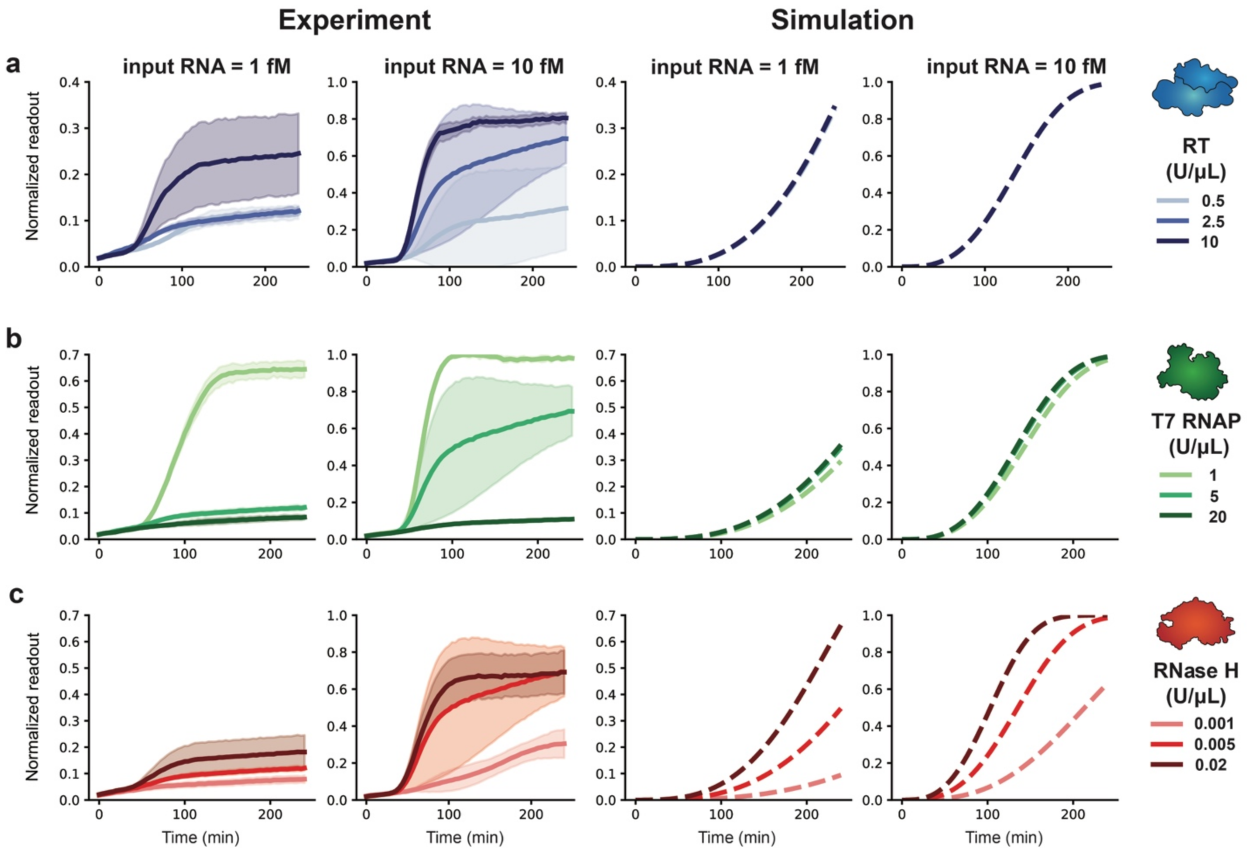


**Supplementary Fig. 12 | Calibration and analysis of sub-optimal candidate model A for Data set 1. a–c**, Time course trajectories for data subsets with simulated data generated with **a**, mid-range RNase H and T7 RNAP and high Cas13a-gRNA, **b**, mid-range RNase H and RT and high Cas13a-gRNA and **c**, mid-range T7 RNAP and RT and high Cas13a-gRNA.

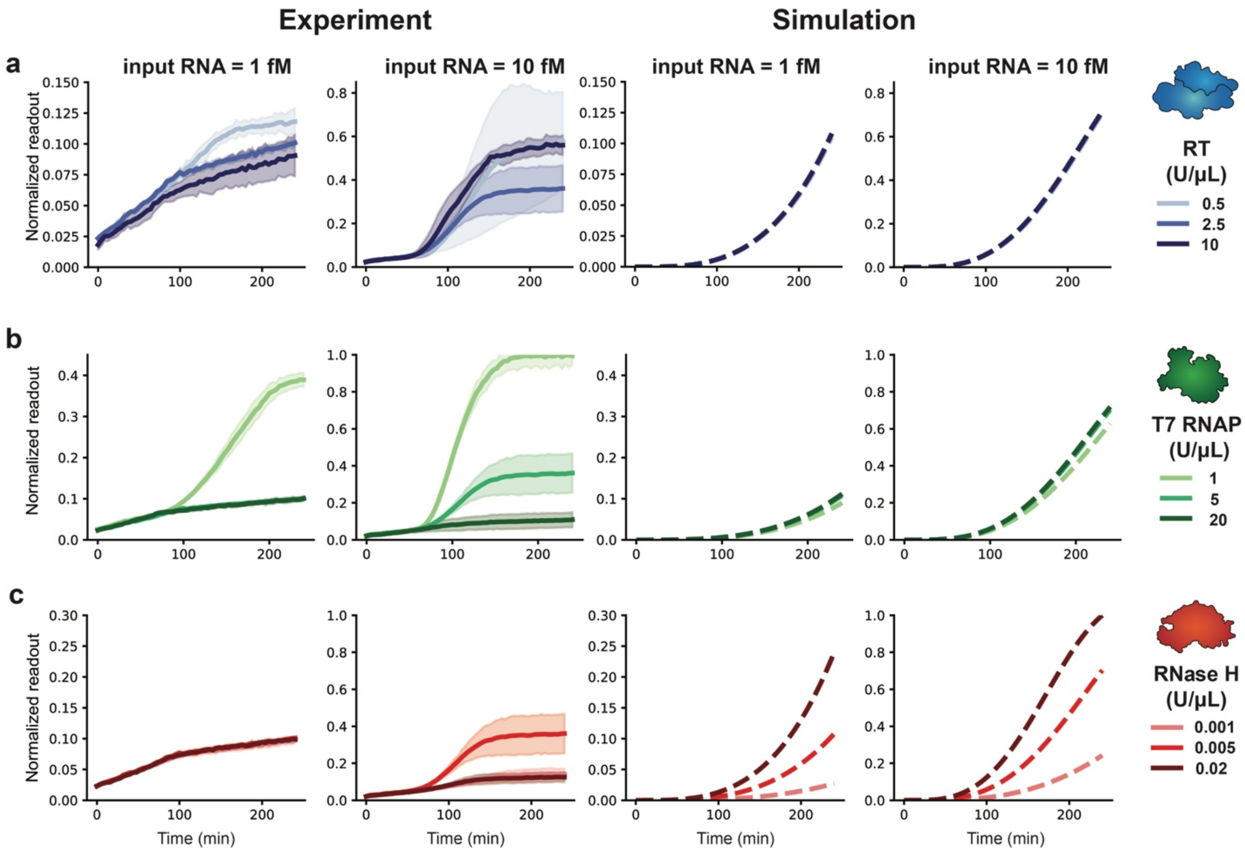


**Supplementary Fig. 13 | Calibration and analysis of sub-optimal candidate model A for Data Set 2. a**–**c**, Time course trajectories for data subsets with simulated data generated with **a**, mid-range RNase H and T7 RNAP and high Cas13a-gRNA, **b**, mid-range RNase H and RT and high Cas13a-gRNA and **c**, mid-range T7 RNAP and RT and high Cas13a-gRNA.

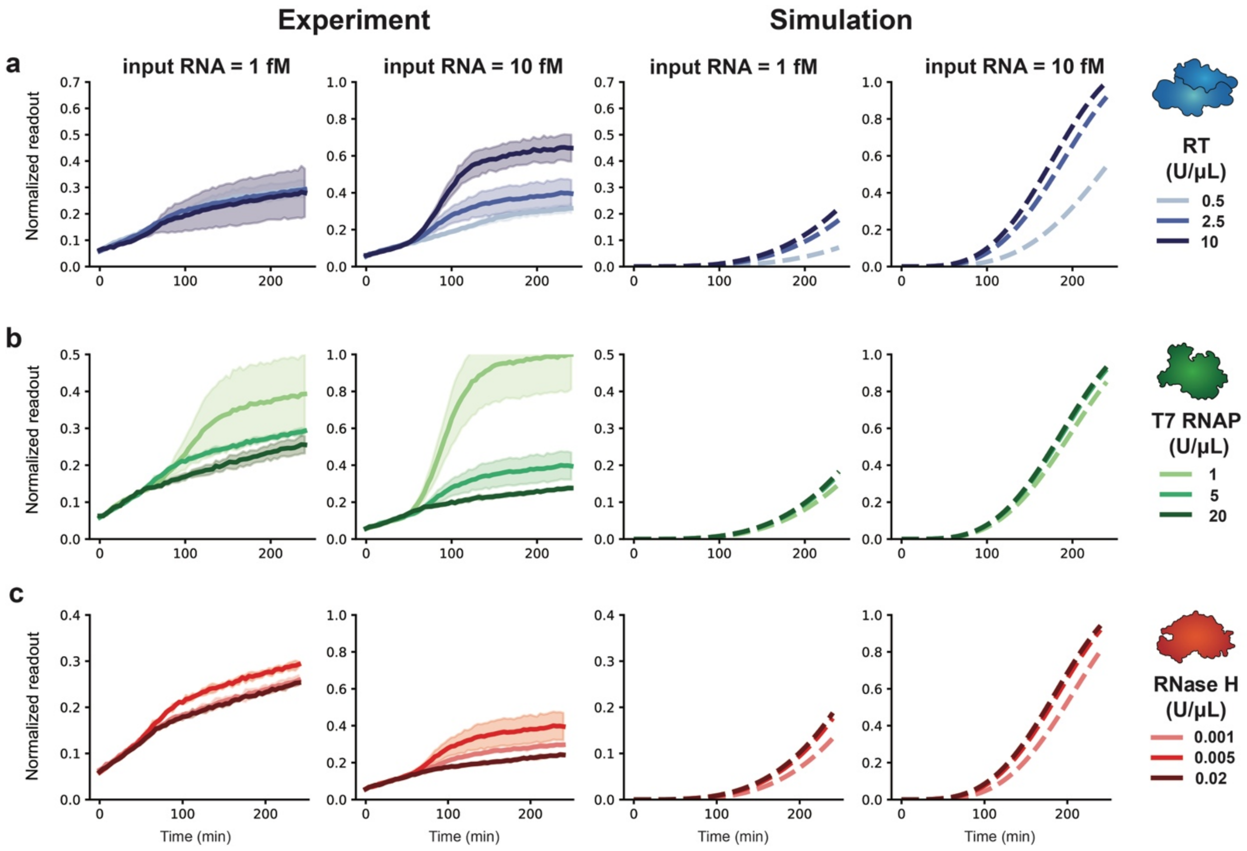


**Supplementary Fig. 14 | Calibration and analysis of sub-optimal candidate model A for Data Set 3. a–c**, Time course trajectories for data subsets with simulated data generated with **a**, mid-range RNase H and T7 RNAP and high Cas13a-gRNA, **b**, mid-range RNase H and RT and high Cas13a-gRNA and **c**, mid-range T7 RNAP and RT and high Cas13a-gRNA.

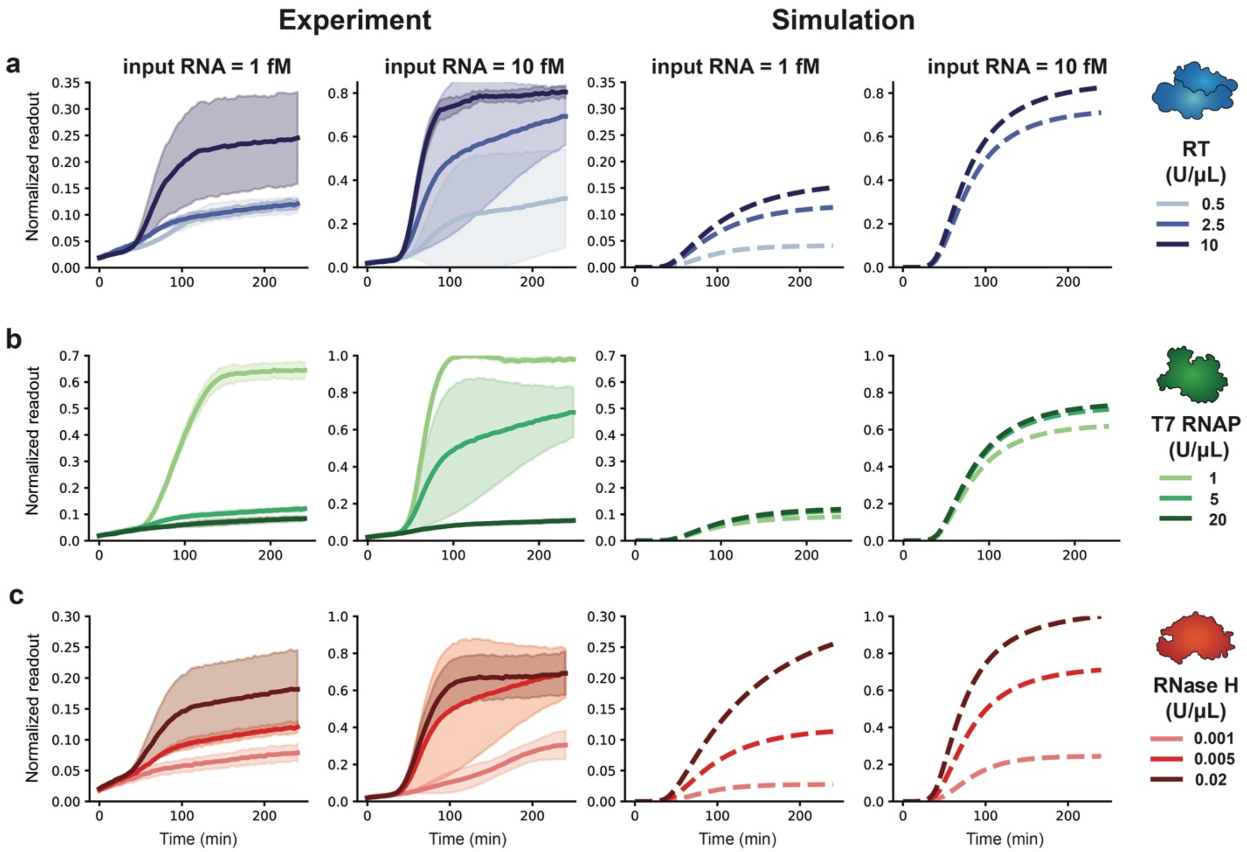


**Supplementary Fig. 15 | Calibration and analysis of sub-optimal candidate model B for Data Set 1. a**–**c**, Time course trajectories for data subsets with simulated data generated with **a**, mid-range RNase H and T7 RNAP and high Cas13a-gRNA, **b**, mid-range RNase H and RT and high Cas13a-gRNA and **c**, mid-range T7 RNAP and RT and high Cas13a-gRNA.

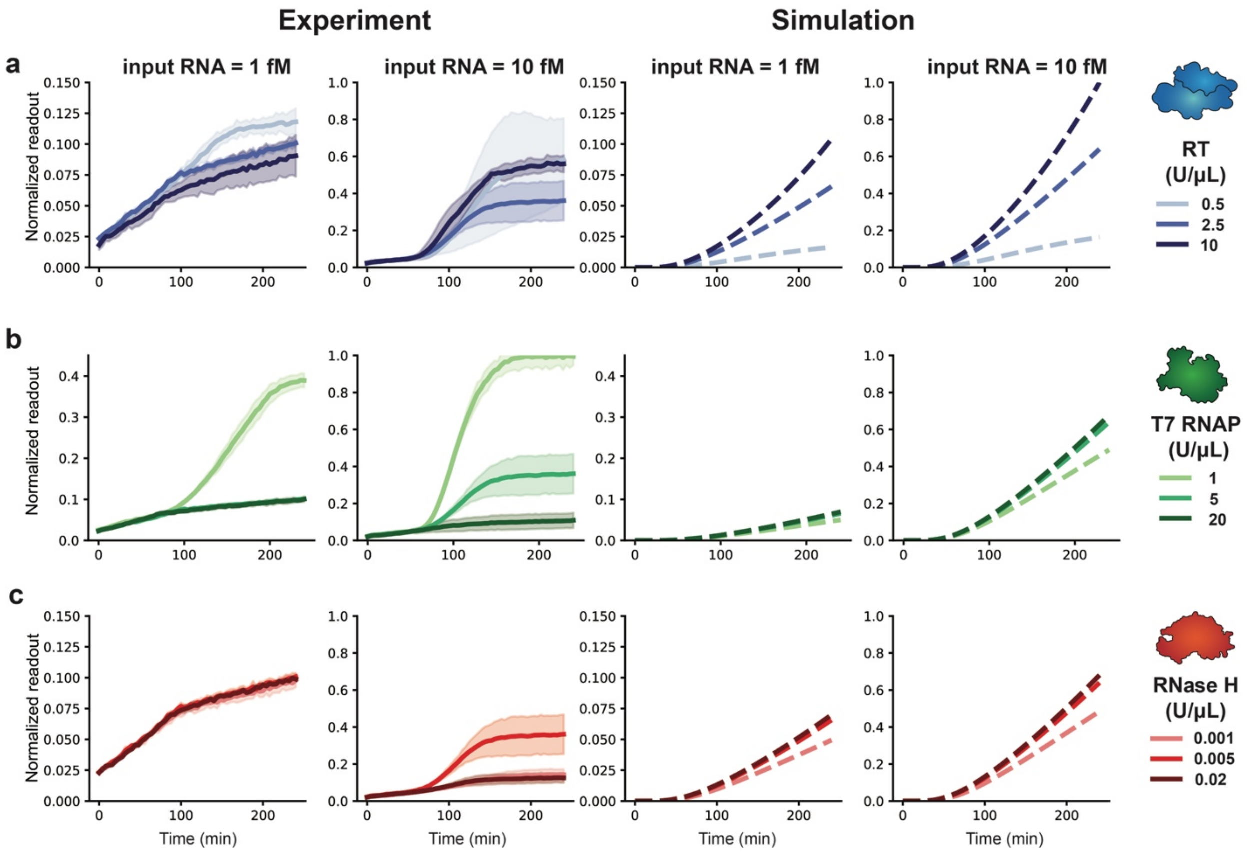


**Supplementary Fig. 16 | Calibration and analysis of sub-optimal candidate model B for data Set 2. a–c**, Time course trajectories for data subsets with simulated data generated with **a**, mid-range RNase H and T7 RNAP and high Cas13a-gRNA, **b**, mid-range RNase H and RT and high Cas13a-gRNA and **c**, mid-range T7 RNAP and RT and high Cas13a-gRNA. The calibrated parameter set for this model and data set did not pass the cost function filter (**Materials and Methods**), which is a further indication that it is not able to describe the training data.

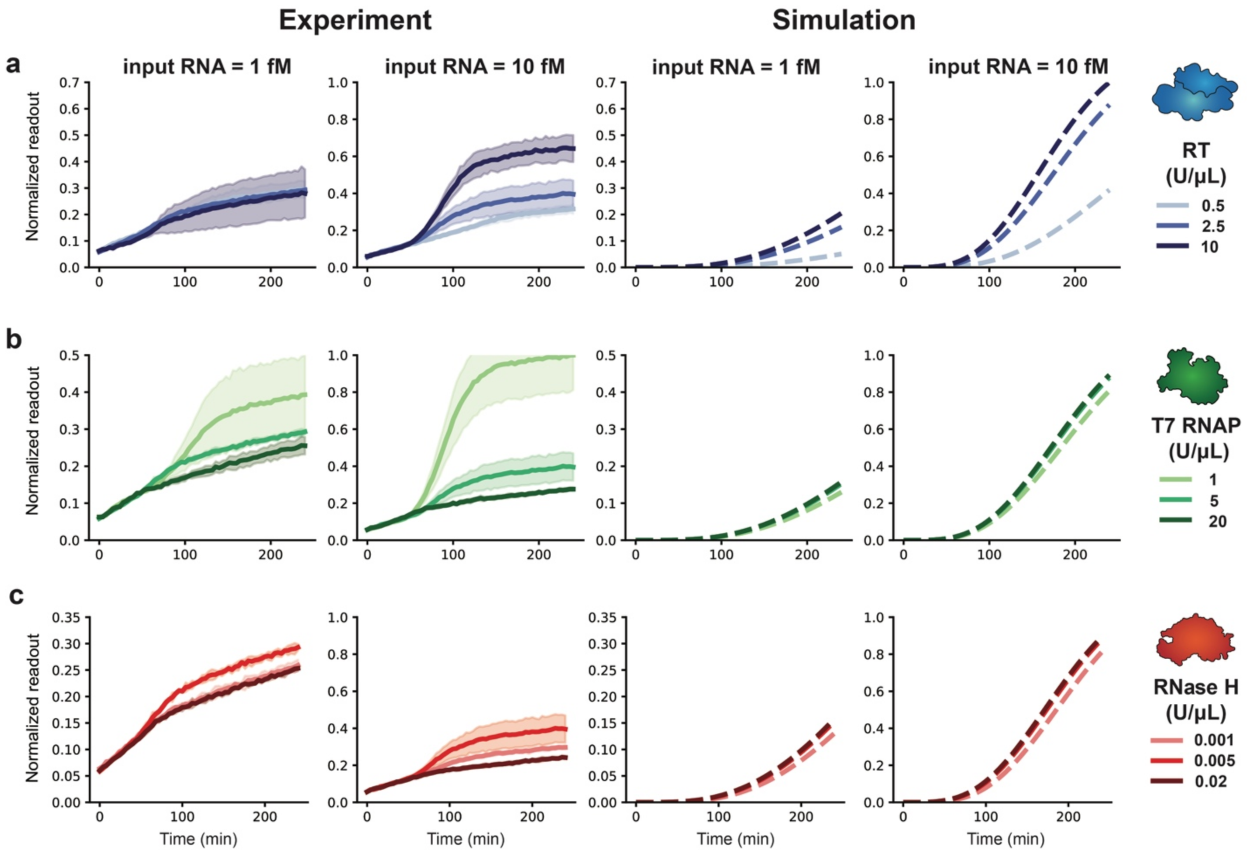


**Supplementary Fig. 17 | Calibration and analysis of sub-optimal candidate model B for Data Set 3. a–c**, Time course trajectories for data subsets with simulated data generated with **a**, mid-range RNase H and T7 RNAP and high Cas13a-gRNA, **b**, mid-range RNase H and RT and high Cas13a-gRNA and **c**, mid-range T7 RNAP and RT and high Cas13a-gRNA.

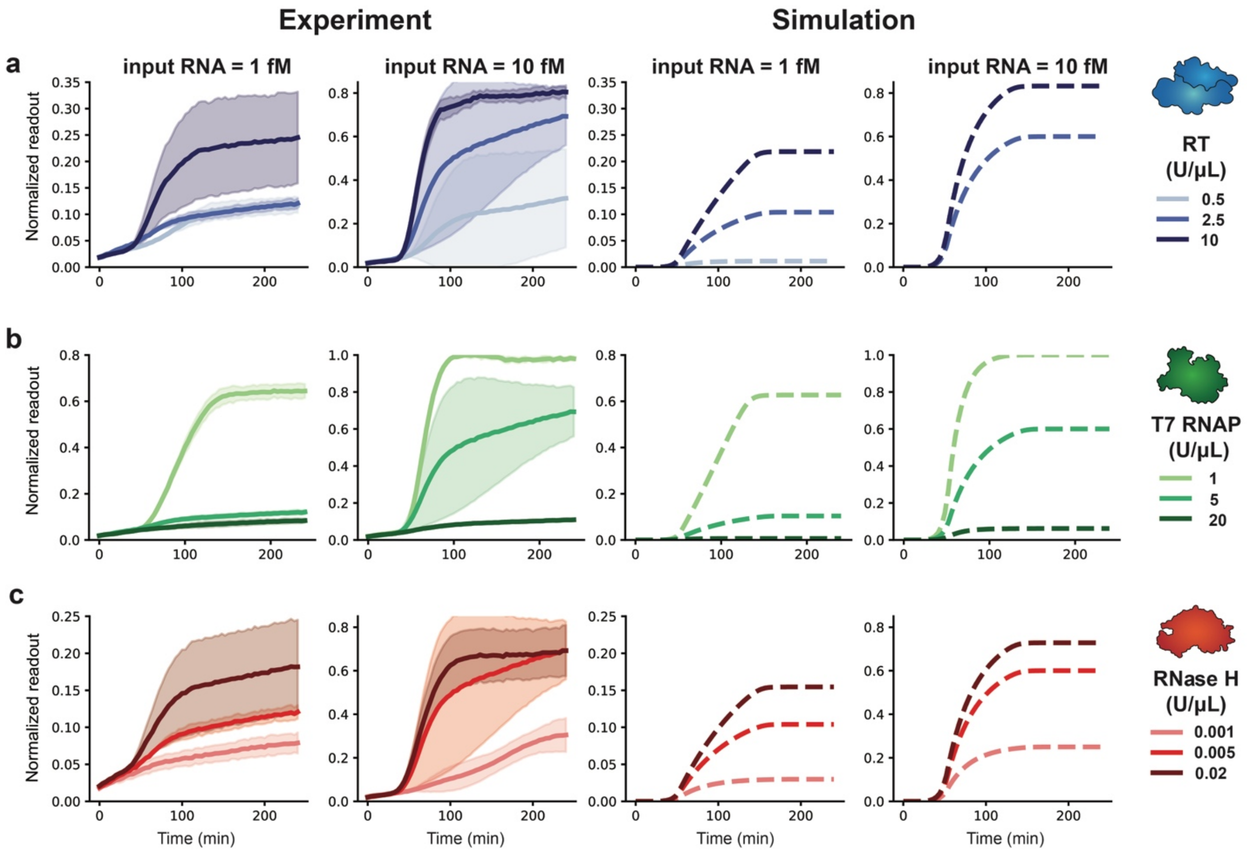


**Supplementary Fig. 18 | Calibration and analysis of sub-optimal candidate model C for Data Set 1. a–c**, Time course trajectories for data subsets with simulated data generated with **a**, mid-range RNase H and T7 RNAP and high Cas13a-gRNA, **b**, mid-range RNase H and RT and high Cas13a-gRNA and **c**, mid-range T7 RNAP and RT and high Cas13a-gRNA.

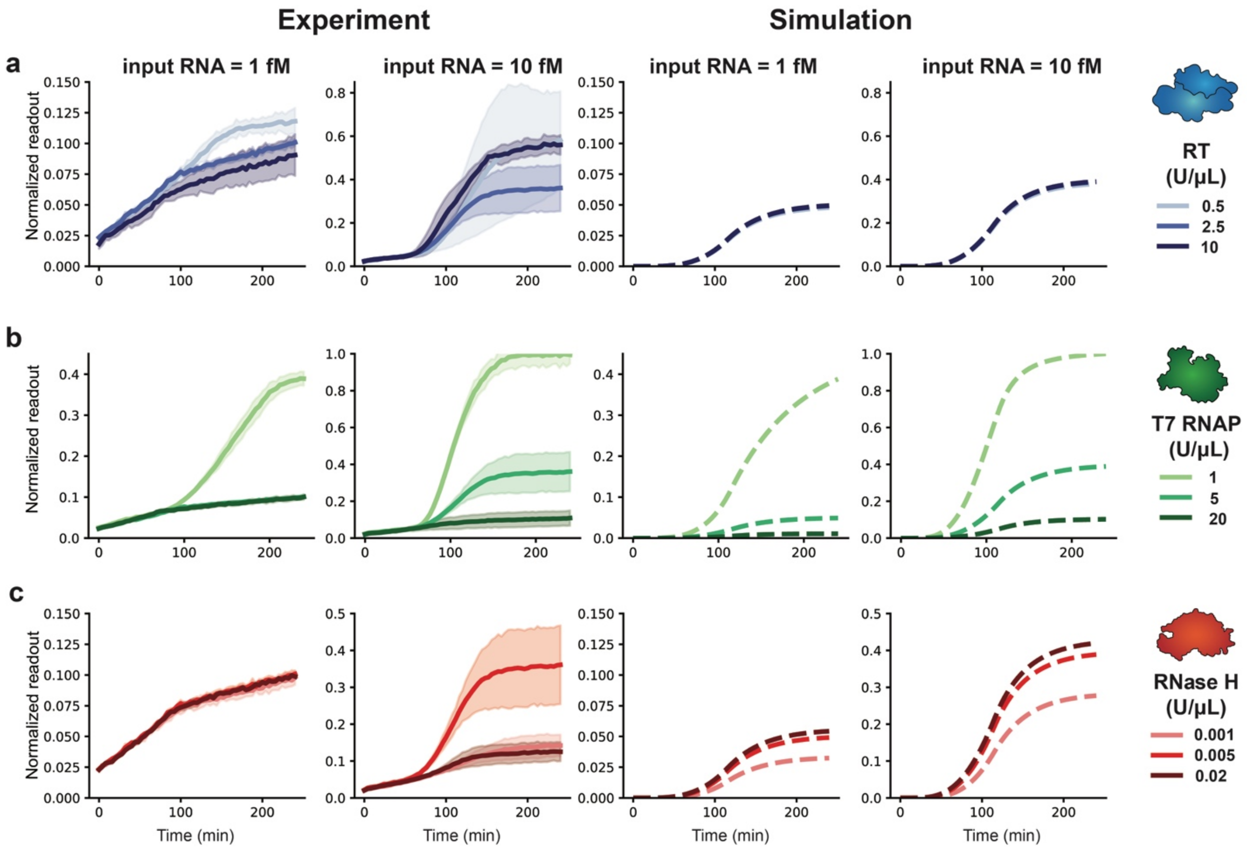


**Supplementary Fig. 19 | Calibration and analysis of sub-optimal candidate model C for Data Set 2. a**–**c**, Time course trajectories for data subsets with simulated data generated with **a**, mid-range RNase H and T7 RNAP and high Cas13a-gRNA, **b**, mid-range RNase H and RT and high Cas13a-gRNA and **c**, mid-range T7 RNAP and RT and high Cas13a-gRNA.

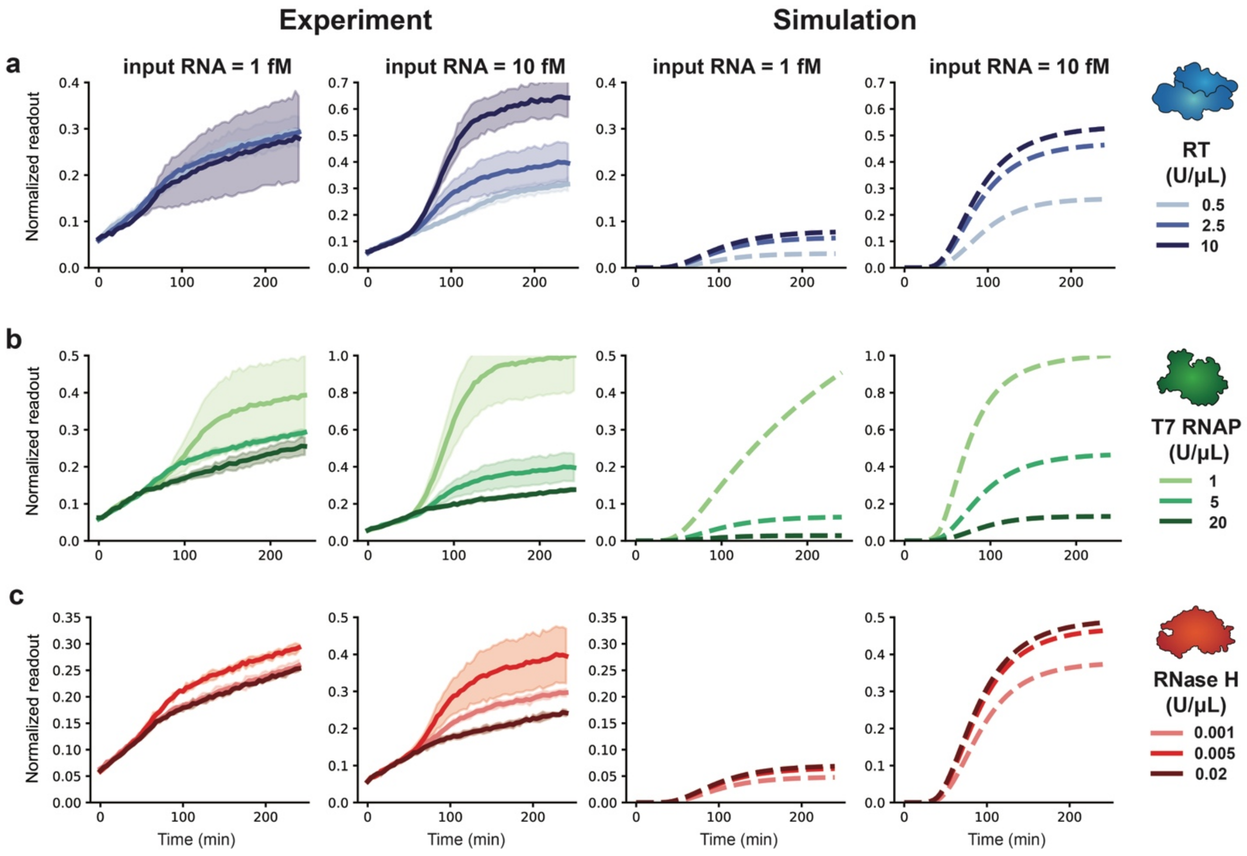


**Supplementary Fig. 20 | Calibration and analysis of sub-optimal candidate model C for Data Set 3. a–c**, Time course trajectories for data subsets with simulated data generated with **a**, mid-range RNase H and T7 RNAP and high Cas13a-gRNA, **b**, mid-range RNase H and RT and high Cas13a-gRNA and **c**, mid-range T7 RNAP and RT and high Cas13a-gRNA.

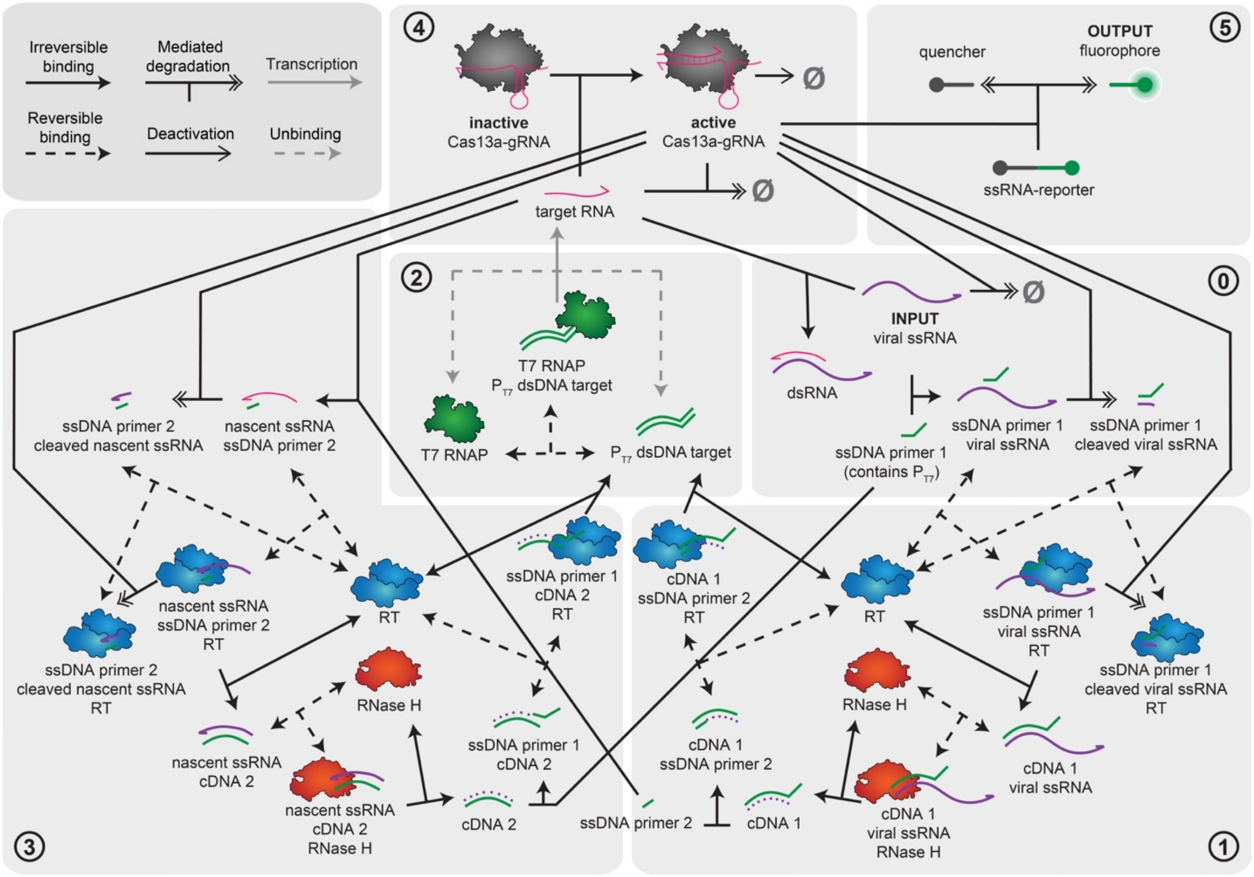


**Supplementary Fig. 21 | Detailed model schematic.** The schematic depicts model states and interactions between model states for the final model (Model C). Circled numbers group related processes into the overall stages of the NASBA-Cas13a mechanism.

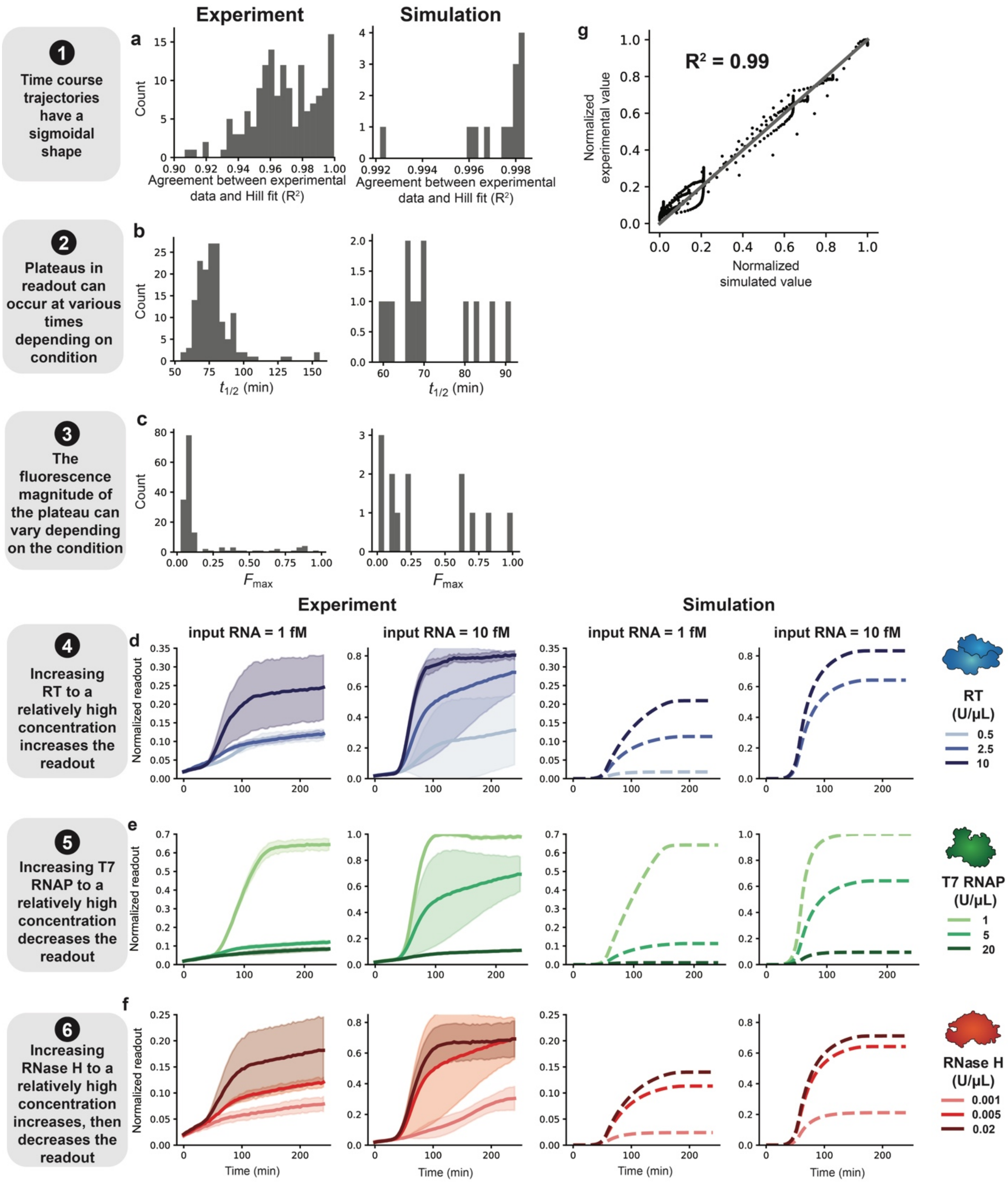


**Supplementary Fig. 22 | Side-by-side comparison of experimental and simulated modeling objectives for Data Set 1. a-c**, Hill-like functions were fit to each time course trajectory, and summary metrics ($n$, $t_{1/2}$, $F_0$, and $F_{max}$) were parameterized (**Fig. 3b** is a visual representation of these metrics). **a,** For each time course, $R^2$ for the normalized data and Hill fit were calculated; values are plotted as a histogram for all conditions in the data set (experimental column) or all conditions in the simulated training data set (simulation column). **b–c**, Histograms of values across all conditions simulated in the data set (experimental column) or all conditions in the training data set (simulation column) were calculated for: **b**, $t_{1/2}$ and **c**, $F_{max}$. **d–f**, Time course trajectories for data subsets: **d**, mid-range RNase H and T7 RNAP and high Cas13a-gRNA, (the 0.5 U/μL condition was omitted as it was removed from the training data due to high measurement error (**Supplementary Fig. 9**) **e**, mid-range RNase H and RT and high Cas13a-gRNA and **f**, mid-range T7 RNAP and RT and high Cas13a-gRNA. **g**, Parity plot for the correlation between normalized experimental data and normalized simulated data. Each point in the plot represents a combination of enzyme doses and time point. In a scenario of perfect agreement, all points would be on the *y* = *x* line (gray).

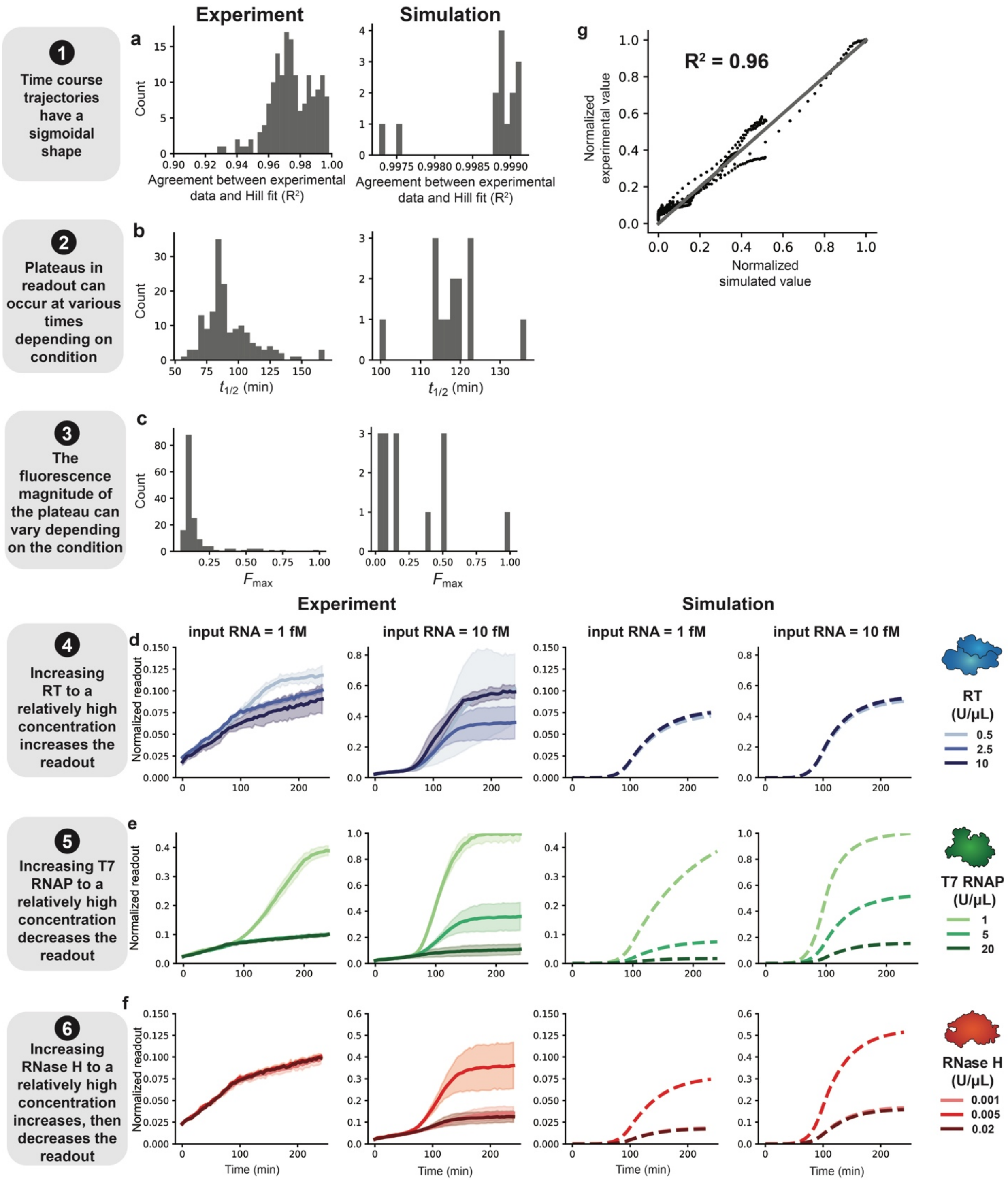


**Supplementary Fig. 23 | Side-by-side comparison of experimental and simulated modeling objectives for Data Set 2. a-c**, Hill-like functions were fit to each time course trajectory, and summary metrics ($n$, $t_{1/2}$, $F_0$, and $F_{max}$) were parameterized (**Fig. 3b** is a visual representation of these metrics). **a,** For each time course, $R^2$ for the normalized data and Hill fit were calculated; values are plotted as a histogram for all conditions in the data set (experimental column) or all conditions in the simulated training data set (simulation column). **b–c**, Histograms of values across all conditions simulated in the data set (experimental column) or all conditions in the training data set (simulation column) were calculated for: **b**, $t_{1/2}$ and **c**, $F_{max}$. **d–f**, Time course trajectories for data subsets: **d**, mid-range RNase H and T7 RNAP and high Cas13a-gRNA **e**, mid-range RNase H and RT and high Cas13a-gRNA and **f**, mid-range T7 RNAP and RT and high Cas13a-gRNA. **g**, Parity plot for the correlation between normalized experimental data and normalized simulated data. Each point in the plot represents a combination of enzyme doses and time point. In a scenario of perfect agreement, all points would be on the *y* = *x* line (gray).

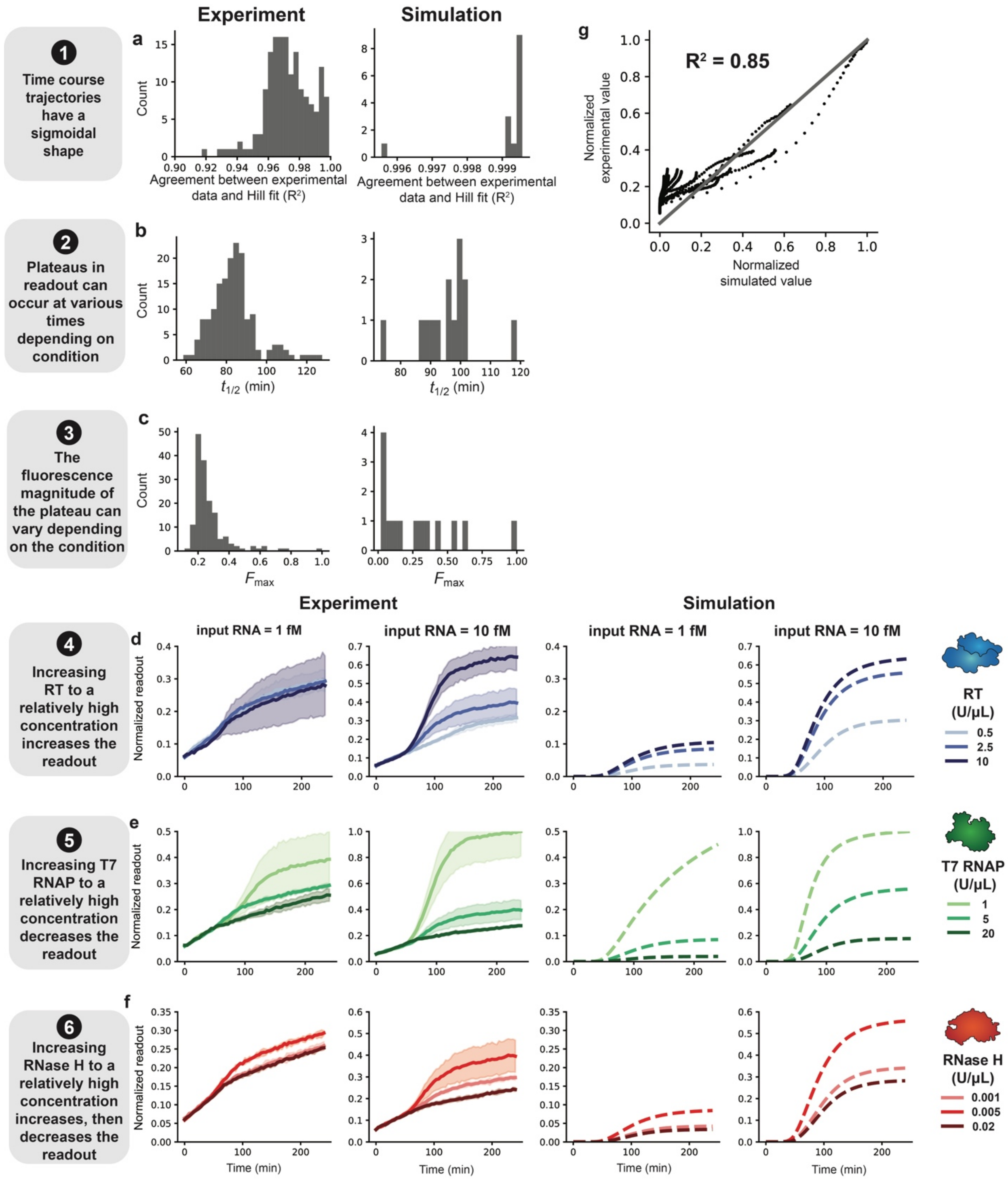


**Supplementary Fig. 24 | Side-by-side comparison of experimental and simulated modeling objectives for Data Set 3. a-c**, Hill-like functions were fit to each time course trajectory, and summary metrics ($n$, $t_{1/2}$, $F_0$, and $F_{max}$) were parameterized (**Fig. 3b** is a visual representation of these metrics). **a,** For each time course, $R^2$ for the normalized data and Hill fit were calculated; values are plotted as a histogram for all conditions in the data set (experimental column) or all conditions in the simulated training data set (simulation column). **b–c**, Histograms of values across all conditions simulated in the data set (experimental column) or all conditions in the training data set (simulation column) were calculated for: **b**, $t_{1/2}$ and **c**, $F_{max}$. **d–f**, Time course trajectories for data subsets: **d**, mid-range RNase H and T7 RNAP and high Cas13a-gRNA, **e**, mid-range RNase H and RT and high Cas13a-gRNA and **f**, mid-range T7 RNAP and RT and high Cas13a-gRNA. **g**, Parity plot for the correlation between normalized experimental data and normalized simulated data. Each point in the plot represents a combination of enzyme doses and time point. In a scenario of perfect agreement, all points would be on the $y = x$ line (gray).

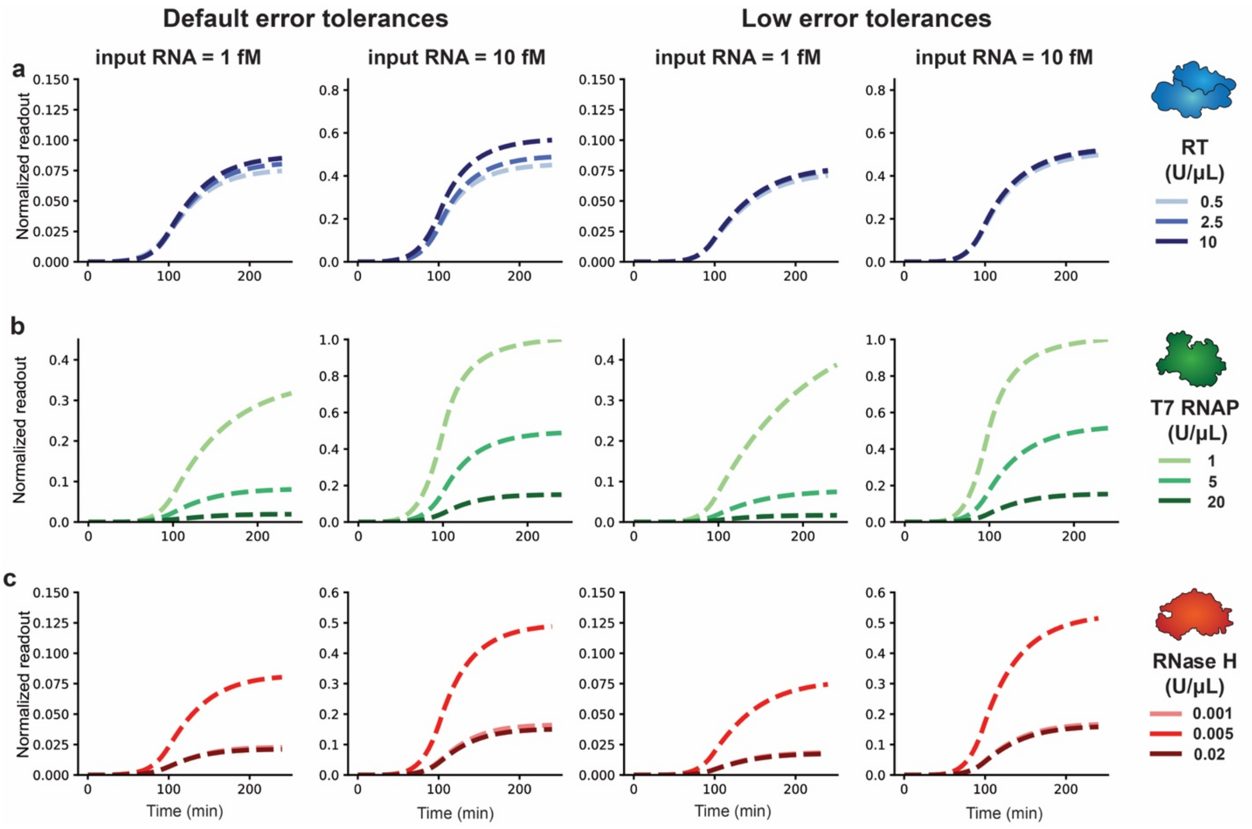


**Supplementary Fig. 25 | Comparison of time course trajectories for optimized parameters with default vs. low ODE solver tolerances for data set 2, model D.** The default solve_ivp error tolerances were initially used to run simulations for parameter estimation, but simulated concentration values sometimes took negative values, so optimization was repeated with decreased error tolerances to check whether parameters were relatively insensitive to these errors (**Materials and Methods**). **a**–**c**, Time course trajectories for data subsets with simulated data using the default error tolerances (left) or the low error tolerances (right) generated with **a**, mid-range RNase H and T7 RNAP and high Cas13a-gRNA, **b**, mid-range RNase H and RT and high Cas13a-gRNA and **c**, mid-range T7 RNAP and RT and high Cas13a-gRNA.

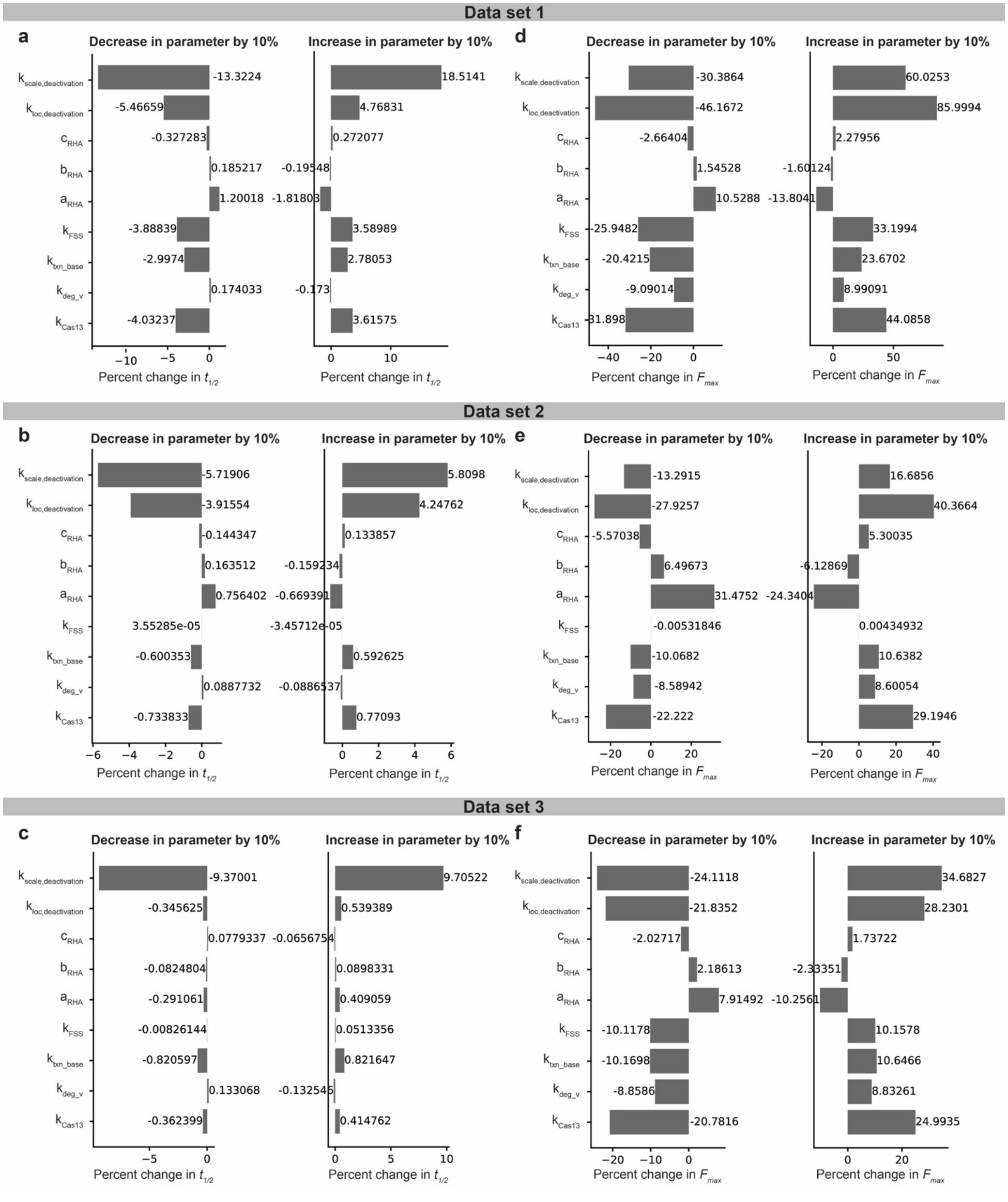


**Supplementary Fig. 26 | Parameter sensitivity analysis using $F_{max}$ or $t_{1/2}$ as a metric. a-c**, Percent change in $t_{1/2}$ when increasing each parameter individually by 10% (left) or decreasing each parameter individually by 10% (right), relative to the metric for the calibrated parameter set, for Data Sets 1-3, respectively. **d-f**, Percent change in $F_{max}$ when increasing each parameter individually by 10% (left) or decreasing each parameter individually by 10% (right), relative to the metric for the calibrated parameter set, for Data Sets 1-3, respectively.

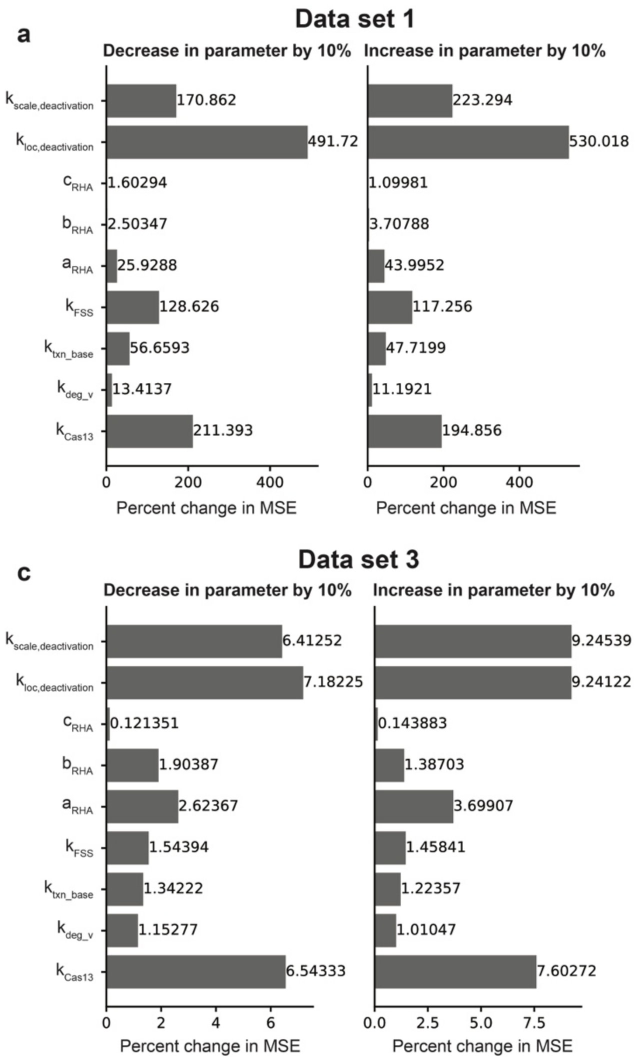


**Supplementary Fig. 27 | Parameter sensitivity analysis using MSE as a metric. a-c**, percent change in MSE when increasing each parameter individually by 10% (left) or decreasing each parameter individually by 10% (right), relative to the MSE for the calibrated parameter set, for Data Sets 1-3, respectively.

### Supplementary Note 1 | Mathematical implementation

*Effective rate constant for T7 RNAP-mediated transcription*

The effective rate constant $k_{\text{txn, effective}}$ was calculated for each initial condition of T7 RNAP, $[\mathbf{x}_{\mathbf{T7\ RNAP}}]_0$, where $k_{\text{txn,base}}$ is a free parameter for base case transcription:

$$k_{\text{txn, effective}} = \frac{k_{\text{txn,base}}}{[\mathbf{x}_{\mathbf{T7\ RNAP}}]_0} \quad \textbf{(Equation S1)}$$

*Cas13 deactivation*

An exponential decay function was used to define deactivation of Cas13a indiscriminate ssRNase activity over time. At each time ($t$), the fraction of active Cas13a ($frac_{Cas13}$) was calculated using the Python package SciPy's stats module[1] `expon` function:

$$frac_{Cas13} = expon(t,\ k_{\text{loc,deactivation}},\ k_{\text{scale,deactivation}}) \quad \textbf{(Equation S2)}$$

$k_{\text{loc,deactivation}}$ and $k_{\text{scale,deactivation}}$ are parameters that define the exponential decay function. $k_{\text{loc,deactivation}}$ determines the time between initialization of the simulation and the start of deactivation, and $k_{\text{scale,deactivation}}$ determines the rate of deactivation once it has started. The total activated Cas13a (i.e., with indiscriminate ssRNase activity) ($\mathbf{x}_{\mathbf{aCas13}}$) is calculated as follows, where $\mathbf{x}_{\mathbf{Cas13}}$ is the total target-activated Cas13a:

$$x_{aCas13} = frac_{Cas13} \cdot \mathbf{x}_{\mathbf{Cas13}} \quad \textbf{(Equation S3)}$$

*Non-monotonic RNase H activity*

A beta distribution was used to define $k_{\text{RHA}}$ for each initial condition of RNase H, $\mathbf{x}_{\mathbf{RNase\ H,0}}$, where $a_{RHA}$ and $b_{RHA}$ are shape parameters for the beta distribution and $c_{RHA}$ determines the scaling on the distribution:

$$k_{\text{RHA}} = \text{c}_{\text{RHA}} \cdot (x_{RNase\ H,0})^{a_{RHA}-1} \cdot \left(1 - x_{RNase\ H,0}\right)^{b_{RHA}-1} \quad \textbf{(Equation S4)}$$

RNase H initial conditions were scaled such that each condition was between 0 and 1, which are the bounds of the beta distribution.

*Re-scaling*

To avoid numerical instability, we re-scaled $\text{x}_{\text{input RNA}}$ such that:

$$\text{x}'_{\text{input RNA}} = C_{\text{scale}} \cdot \mathbf{x}_{\mathbf{input\ RNA}} \quad \textbf{(Equation S5)}$$

With $C_{\text{scale}} = 10^6$. We show separate derivations for the ODE describing $\mathbf{x}_{\text{input RNA}}$ and all other ODEs. For the ODE that tracks $\mathbf{x}_{\text{input RNA}}$, we used the following derivation for rescaling:

$$\frac{d\mathbf{x}_{\text{input RNA}}}{dt} = k \cdot \mathbf{x}_{\text{input RNA}} \cdot \mathbf{x}_{\text{other}} \qquad \textbf{(Equation S6)}$$

where $\boldsymbol{x}_{\text{other}}$ is all state variables that are not $\mathbf{x}_{\text{input RNA}}$. Substituting $\frac{\mathbf{x}'_{\text{input RNA}}}{C_{\text{scale}}}$ for $x_v$, the equation is:

$$\frac{1}{C_{\text{scale}}} \cdot \frac{d\mathbf{x}'_{\text{input RNA}}}{dt} = k \cdot \frac{1}{C_{\text{scale}}} \cdot \mathbf{x}'_{\text{input RNA}} \cdot \mathbf{x}_{\text{other}} \qquad \textbf{(Equation S7)}$$

After simplification, the equation becomes:

$$\frac{d\mathbf{x}'_{\text{input RNA}}}{dt} = k \cdot \mathbf{x}'_{\text{input RNA}} \cdot \mathbf{x}_{\text{other}} \qquad \textbf{(Equation S8)}$$

Therefore, no changes are required for the ODE that tracks $\mathbf{x}_{\text{input RNA}}$. For each remaining term involving $\mathbf{x}_{\text{input RNA}}$, we used the following equation for re-scaling:

$$\frac{d\mathbf{x}_{\text{other}}}{dt} = k \cdot \mathbf{x}_{\text{input RNA}} \cdot \mathbf{x}_{\text{other}} + k_2 \cdot \mathbf{x}_{\text{other}} \cdot \mathbf{x}_2 \qquad \textbf{(Equation S9)}$$

After substituting $\frac{\mathbf{x}'_{\text{input RNA}}}{C_{\text{scale}}}$ for $\boldsymbol{x}_v$, the equation becomes:

$$\frac{d\mathbf{x}_{\text{other}}}{dt} = k \cdot \frac{1}{C_{\text{scale}}} \cdot \mathbf{x}'_{\text{input RNA}} \cdot \mathbf{x}_{\text{other}} + k_2 \cdot \mathbf{x}_{\text{other}} \cdot \mathbf{x}_2 \qquad \textbf{(Equation S10)}$$

Therefore, for each term including $\mathbf{x}'_{\text{input RNA}}$ (in all equations except for the one that tracks $\mathbf{x}_{\text{input RNA}}$), the term is divided by $C_{\text{scale}}$. Re-scaling is accounted for in **Supplementary Table 4**.

*Conservation laws*

Conservation laws were applied to internal model states that by definition are conserved. At each time step, the following equations were used to calculate concentrations. In each equation, the first term after the equals sign is the initial value.

$$\begin{aligned}\mathbf{x}_{\text{p1}} = \mathbf{x}_{\text{p1,0}} &- \mathbf{x}_{\text{p1v}} - \mathbf{x}_{\text{p1cv}} - \mathbf{x}_{\text{RTp1v}} - \mathbf{x}_{\text{RTp1cv}} - \mathbf{x}_{\text{cDNA1v}} - \mathbf{x}_{\text{RNasecDNA1v}} \\ &- \mathbf{x}_{\text{cDNA1}} - \mathbf{x}_{\text{p2cDNA1}} - \mathbf{x}_{\text{p1cDNA2}} - \mathbf{x}_{\text{RTp2cDNA1}} - \mathbf{x}_{\text{RTp1cDNA2}} \\ &- \mathbf{x}_{\text{pro}} - \mathbf{x}_{\text{T7pro}}\end{aligned} \qquad \textbf{(Equation S11)}$$

$$\begin{aligned}\mathbf{x}_{\text{p2}} = \mathbf{x}_{\text{p2,0}} &- \mathbf{x}_{\text{p2u}} - \mathbf{x}_{\text{p2cu}} - \mathbf{x}_{\text{RTp2u}} - \mathbf{x}_{\text{RTp2cu}} - \mathbf{x}_{\text{cDNA2u}} - \mathbf{x}_{\text{RNasecDNA2u}} \\ &- \mathbf{x}_{\text{cDNA2}} - \mathbf{x}_{\text{p2cDNA1}} - \mathbf{x}_{\text{p1cDNA2}} - \mathbf{x}_{\text{RTp2cDNA1}} - \mathbf{x}_{\text{RTp1cDNA2}} \\ &- \mathbf{x}_{\text{pro}} - \mathbf{x}_{\text{T7pro}}\end{aligned} \qquad \textbf{(Equation S12)}$$

$$\begin{aligned}\mathbf{x}_{\text{RT}} = \mathbf{x}_{\text{RT,0}} &- \mathbf{x}_{\text{RTp1v}} - \mathbf{x}_{\text{RTp2u}} - \mathbf{x}_{\text{RTp1cv}} - \mathbf{x}_{\text{RTp2cu}} - \mathbf{x}_{\text{RTp2cDNA1}} \\ &- \mathbf{x}_{\text{RTp1cDNA2}}\end{aligned} \qquad \textbf{(Equation S13)}$$

$$\mathbf{x_{RNase}} = \mathbf{x_{RNase,0}} - \mathbf{x_{RNasecDNA1v}} - \mathbf{x_{RNasecDNA2u}} \quad \textbf{(Equation S14)}$$

$$\mathbf{x_{T7}} = \mathbf{x_{T7,0}} - \mathbf{x_{T7pro}} \quad \textbf{(Equation S15)}$$

$$\mathbf{x_{iCas13}} = \mathbf{x_{iCas13,0}} - \mathbf{x_{Cas13}} \quad \textbf{(Equation S16)}$$

$$\mathbf{x_{q}} = \mathbf{x_{qRf,0}} - \mathbf{x_{qRf}} \quad \textbf{(Equation S17)}$$

$$\mathbf{x_{f}} = \mathbf{x_{qRf,0}} - \mathbf{x_{qRf}} \quad \textbf{(Equation S18)}$$

**Supplementary Note 2 | Evaluation of parameter estimation method (PEM)**

*Algorithm for PEM evaluation*

Before using the PEM to estimate parameters to describe the training data, we evaluated the ability of the proposed PEM to find parameter sets yielding good agreement with the training data. To accomplish this, we generated three PEM evaluation data sets using three different sets of PEM evaluation parameters (**Supplementary Fig. 10b**). Each PEM evaluation data set has the same structure as the training data, and each data set was generated using the proposed model in question. This process was repeated for each new version of the model (A, B, C, and D). After generation of the PEM evaluation data, parameters were fit to the data, and the ability of the PEM to identify parameter sets with good agreement with the training data was evaluated by calculating the PEM evaluation criterion. The PEM evaluation criterion was satisfied for each model, for each data set (**Supplementary Fig.11a-l**), which supports that the proposed PEM is appropriate for each model given the structure of the training data. PEM evaluation is described further in the initial report of the GAMES workflow[2].

*Generation of PEM evaluation training data*

It is important to accurately represent measurement error when evaluating a parameter estimation method for a given set of training data. For each PEM evaluation data set, we first simulated each component condition (defined as the raw simulated data), then added noise to approximate the experimental measurement error. For each condition, we first calculated the standard error, $\sigma_{SEM}$, associated with each data point, *i*, using the following equation:

$$\sigma_{SEM,i} = \frac{\sigma_{SD,i}}{\sqrt{n_{replicates}}} \quad (\textbf{Equation S19})$$

$\sigma_{SD}$ is the standard deviation of each data point, all of which were collected in triplicate ($n_{replicates}$ = 3).

Next, for each condition, *j*, we calculated the mean, $\mu_j$, and standard deviation, $\sigma_{SD,j}$, of $\sigma_{SEM}$ across all data points for each condition. $\mu_j$ and $\sigma_{SD,j}$ were then used to define a normal distribution from which values were randomly generated and added or subtracted to each data point in the raw simulated data. If subtracting a noise value led a data point that was less than 0, another random number was generated until the data point with noise was greater than or equal to 0.

To determine whether the noise value was added or subtracted from the raw data point, we used a random number generator that output a value of either 0 or 1. If the value was 0, the noise value was subtracted from the raw value and if the value was 1, the noise value was added to the raw value.

We found that, for all conditions, measurement error was near zero for the first 10 time points, so we did not include these data points in the mean measurement error calculations. We also did not add noise to these data points.

*Determination of PEM evaluation criterion*
For each PEM evaluation data set, we calculated the $R^2$ value between the raw simulated data and the simulated data with added noise. The minimum $R^2$ value across the 3 PEM evaluation data sets was used to define the PEM evaluation criterion.

### Supplementary Note 3 | Calibration and analysis of suboptimal candidate models

*Parameter estimation method details*

For all models, parameter estimation simulations were initiated with the literature values in **Supplementary Table 2** and allowed to vary 3 orders of magnitude in each direction. We used the literature values for the free parameters ($k_{\mathrm{Cas13}}$, $k_{\mathrm{deg\_v}}$, $k_{\mathrm{txn_{base}}}$, $k_{\mathrm{FSS}}$, $k_{\mathrm{RHA}}$) only as order of magnitude estimates, as these values were all determined with different systems and under different conditions than our training data. In initial simulations, we found that large values of $k_{\mathrm{Cas13}}$ led to stiff dynamics that stalled out the ODE solver, even when an algorithm appropriate for stiff ODEs was used, so we initialized $k_{\mathrm{Cas13}}$ at a value 2 orders of magnitude below the literature estimate. Literature estimates for $k_{\mathrm{loc,deactivation}}$ and $k_{\mathrm{scale,deactivation}}$ were unavailable. These parameters were each allowed to vary between 0 and 240. Bounds for the beta distribution shape parameters, $a_{RHA}$ and $b_{RHA}$, were based on visual inspection of the beta distribution to enable non-monotonic behavior in the regime of the scaled $[RNase\ H]_0$ values. $a_{RHA}$ was allowed to vary between 1 and 10 and $b_{RHA}$ was allowed to vary between 1 and 100. Because $c_{RHA}$ acted as a scaling parameter and did not change the shape of the beta distribution, it was allowed to vary between $10^{-3}$ and $10^{3}$.

For Models A, C, and D, we found it necessary to implement a timeout function in the global search such that if a parameter set takes too long to solve (in our case, we used t = 100s as the timeout condition), it was skipped, and the cost function was set to an arbitrarily high value of 3. Without the timeout function, the global search simulation stalled indefinitely, making the subsequent optimization step impossible. Our timeout function was incompatible with our parallelization code, so Models A and C were run without parallelization. Model B did not require the timeout function and was run with parallelization. We hypothesize that the stalling phenomenon may occur for only some model structures and parameter sets because the ODE solver may be forced to take extremely small time steps in some parameter regimes. Skipping these parameter sets in the global search is appropriate because only a few parameter sets were thrown out, therefore not significantly affecting the downstream results.

We note that this timeout function was an extremely important part of our modeling work. Without the timeout function, we were forced to rely on manual parameter estimation, as our parameter estimation workflow could not be run from start-to-finish without stalling out. This non-rigorous, manual parameter estimation method initially led us to erroneously conclude that Model A did not meet any of the modeling objectives and we subsequently added in an additional mechanism to slow down the reactions. After adding in the timeout function and running our full parameter estimation workflow, we concluded that this additional mechanism was in fact not necessary. This anecdote highlights the importance of using an appropriately rigorous parameter estimation workflow, even when extra effort is required to enable the workflow to run (in our case, the timeout function).

*Further analysis of sigmoidal behavior of the fits to data sets 2 and 3 for models A and B*

The calibrated parameters for models A and B for data sets 2 and 3 did not result in a visually sigmoidal behavior (**Supplementary Figs. 13-14** for model A and **Supplementary Figs. 16-17** for model B), although the Hill-fit metrics did enable each one to satisfy modeling Objectives 1 and 2 (**Table 1**). In each case, we performed further analysis to determine whether we could achieve a more visually sigmoidal behavior. For each model and data set, we initialized the PEM optimization (**Materials and Methods**) with the corresponding calibrated parameter values for the fit to data set 1, which did exhibit a visually sigmoidal behavior for models A and B (**Supplementary Figs. 12** and **15**, respectively). We found that this approach did not result in a visually sigmoidal behavior for either model or data set, which indicates that it was not possible to achieve a visually sigmoidal behavior for the fits to data sets 2 and 3 with these mechanisms alone.

**Supplementary Note 4 | Parameter identifiability**

One limitation of the model is that not all parameters are identifiable, i.e., capable of being uniquely estimated within a finite confidence interval given the training data. Therefore, if the model were used to make predictions, such as of an optimal enzyme concentration or predicting the impact of a strategic intervention designed to decrease the time to readout, these predictions might also not be constrained and could be difficult to validate. For any case in which the model is used to make predictions, parameter identifiability analysis could guide model reduction or experimental design with the goal of arriving at a fully identifiable model with well-constrained predictions[3]. However, as the path to an identifiable model often changes depending on the type of prediction that is desired, such an analysis is beyond our current scope, as we focus on the insight gained from the explanatory model.

**Supplementary Table 1 | Internal model states[1].**

| # | States | Name | Initial Value |
|---|---|---|---|
| 0 | x_inputRNA | viral ssRNA (INPUT) | **0, 1, or 10 fM → 0, $10^{-6}$, or $10^{-7}$ nM** |
| 1 | x_p1 | ssDNA primer 1 | 250 nM |
| 2 | x_p2 | ssDNA primer 2 | 250 nM |
| 3 | x_p1v | ssDNA primer : viral ssRNA | 0 |
| 4 | x_p2u | ssDNA primer : target ssRNA | 0 |
| 5 | x_p1vdeg | ssDNA primer : cleaved viral ssRNA | 0 |
| 6 | x_p2udeg | ssDNA primer : cleaved target ssRNA | 0 |
| 7 | x_RT | RT | **0.5, 2.5, or 10 U/µL → 69.55, 347.75, 1391.0 nM** |
| 8 | x_RNase | RNase H | **0.001, 0.005, or 0.02 U/µL → 6.06, 30.3, 121.2 nM** |
| 9 | x_RTp1v | RT-ssDNA primer : viral ssRNA | 0 |
| 10 | x_RTp2u | RT-ss DNA primer : target ssRNA | 0 |
| 11 | x_RTp1vdeg | RT-ssDNA primer : cleaved viral ssRNA | 0 |
| 12 | x_RTp2udeg | RT-ssDNA primer : cleaved target ssRNA | 0 |
| 13 | x_cDNA1v | cDNA : viral ssRNA | 0 |
| 14 | x_cDNA2u | cDNA : target ssRNA | 0 |
| 15 | x_RNasecDNA1v | cDNA : viral ssRNA : RNase H | 0 |
| 16 | x_RNasecDNA2u | cDNA : target ssRNA : RNase H | 0 |
| 17 | x_cDNA1 | cDNA : ssRNA fragments | 0 |
| 18 | x_cDNA2 | cDNA : ssRNA fragments (DNA 2) | 0 |
| 19 | x_p2cDNA1 | cDNA : ssDNA primer 2 | 0 |
| 20 | x_p1cDNA2 | cDNA 2 : ssDNA primer | 0 |
| 21 | x_RTp2cDNA1 | cDNA : ssDNA primer 2 : RT/RNase H | 0 |
| 22 | x_RTp1cDNA2 | cDNA 2 : ssDNA primer : RT/RNase H | 0 |
| 23 | x_T7 | T7 RNAP | **1, 5, or 20 U/µL → 16.16, 80.8, 323.2 nM** |
| 24 | x_pro | dsDNA T7 promoter target | 0 |
| 25 | x_T7pro | T7 RNAP : dsDNA T7 promoter target | 0 |
| 26 | x_u | ssRNA target | 0 |
| 27 | x_iCas13 | Target-inactivated Cas13a-gRNA | **2.25 or 45 nM** |

| | | | |
|---|---|---|---|
| 28 | `x_Cas13` | Target-activated Cas13-gRNA | 0 |
| 29 | `x_uv` | dsRNA | 0 |
| 30 | `x_qRf` | quencher-ssRNA-fluorophore | 2500 nM |
| 31 | `x_q` | quencher | 0 |
| 32 | `x_f` | fluorophore (OUTPUT) | 0 |

[1] Initial values in bold vary in the training data set, and other initial values remain constant. In the text, `x_iCas13` is target-inactivated Cas13a-gRNA and `x_Cas13` is target-activated Cas13a-gRNA. In the initial value column, arrows represent conversions from experimental units (fM or U/μL) to model units (nM).

**Supplementary Table 2 | Parameter labels and descriptions[1].**

| # | Parameter | Free/Fixed | Value (if fixed) or guess (if free) | Units | Description |
|---|---|---|---|---|---|
| 1 | $k_{\mathrm{Cas13}}$ | Free | 0.198 | $nM^{-1}$ $min^{-1}$ | Binding of Cas13-gRNA to RNA target[4] |
| 2 | $k_{\mathrm{deg\_v}}$ | Free | 30.6 | $nM^{-1}$ $min^{-1}$ | Degradation of viral ssRNA by active Cas13-gRNA[5] |
| 3 | $k_{\mathrm{txn\_base}}$ | Free | 36 | $min^{-1}$ | T7 RNAP-induced transcription of RNA[6] |
| 4 | $k_{\mathrm{T7on}}$ | Fixed | 3.36 | $nM^{-1}$ $min^{-1}$ | Binding of T7 RNAP and dsDNA T7 promoter[6] |
| 5 | $k_{\mathrm{T7off}}$ | Fixed | 12 | $min^{-1}$ | Unbinding of T7 RNAP and dsDNA T7 promoter[6] |
| 6 | $k_{b\mathrm{ds}}$ | Free, but not independent | = $k_{\mathrm{Cas13}}$ | $nM^{-1}$ $min^{-1}$ | Binding of complementary double strand[4] |
| 7 | $k_{\mathrm{RTon}}$ | Fixed | 0.024 | $nM^{-1}$ $min^{-1}$ | Binding of RT or RNase H[7] |
| 8 | $k_{\mathrm{RToff}}$ | Fixed | 2.4 | $min^{-1}$ | Unbinding of RT or RNase H[7] |
| 9 | $k_{\mathrm{RNaseon}}$ | Fixed | 0.024 | $nM^{-1}$ $min^{-1}$ | Binding of RT or RNase H[7] |
| 10 | $k_{\mathrm{RNaseoff}}$ | Fixed | 2.4 | $min^{-1}$ | Unbinding of RT or RNase H |
| 11 | $k_{\mathrm{FSS}}$ | Free | 0.6 | $min^{-1}$ | First strand synthesis[8] |
| 12 | $k_{\mathrm{RHA}}$ | Free | 7.8 | $min^{-1}$ | RNase H activity |
| 13 | $a_{\mathrm{RHA}}$ | Free | - | unitles s | Determines shape of beta distribution for RNase H heuristic |
| 14 | $b_{\mathrm{RHA}}$ | Free | - | unitles s | Determines shape of beta distribution for RNase H heuristic |
| 15 | $c_{\mathrm{RHA}}$ | Free | 1.0 | unitles s | Determines scaling of beta distribution for RNase H heuristic |
| 16 | $k_{\mathrm{SSS}}$ | Free, but not independent | = $k_{\mathrm{FSS}}$ | $min^{-1}$ | Second strand synthesis[8] |

| | | | | | |
|---|---|---|---|---|---|
| 17 | $k_{\mathrm{degRrep}}$ | Free, but not independent | $= k_{\mathrm{deg\_v}}$ | $\mathrm{min}^{-1}$ | Degradation of ssRNA-based reporter[5] |
| 18 | $k_{\mathrm{loc,deactivatio}}$ | Free | - | min | Determines the time between initialization of the simulation and start of Cas13 deactivation |
| 19 | $k_{\mathrm{scale,deactiva}}$ | Free | - | unitles s | Determines the rate of Cas13 deactivation after starting |

[1] Values (for fixed parameters) and initial guesses (for free parameters) are based on the corresponding reference in the last column

**Supplementary Table 3 | Calibrated parameter values[1].**

| Free parameter | Model A | | | Model B | | | Model C | | | Model D | | |
|---|---|---|---|---|---|---|---|---|---|---|---|---|
| | Data set 1 | Data set 2 | Data set 3 | Data set 1 | Data set 2 | Data set 3 | Data set 1 | Data set 2 | Data set 3 | Data set 1 | Data set 2 | Data set 3 |
| $k_{\mathrm{Cas13}}$ | 0.4667 | 0.0009046 | 1.275 | 0.0005361 | 0.001723 | 1.352 | 0.000290109010 | 0.0002293 | 8.676*10$^{-5}$ | 0.0002241 | 2.230*10$^{-5}$ | 6.382*10$^{-5}$ |
| $k_{\mathrm{deg\ \_v}}$ | 10300 | 2514 | 3407 | 7116 | 129.0 | 13710 | 56.67 | 2907 | 20630 | 136.1 | 8940 | 28930 |
| $k_{\mathrm{txn\_base}}$ | 0.03626 | 0.03600 | 0.03603 | 0.8923 | 0.2514 | 0.09305 | 1283 | 1.662 | 45.59 | 1151 | 226.3 | 119.2 |
| $k_{\mathrm{FSS}}$ | 20.41 | 7.529 | 0.01744 | 0.0665506655 | 0.02271 | 0.006773 | 0.092623 | 315.1 | 0.1057 | 0.09603 | 232.9 | 0.07789 |
| $k_{\mathrm{RHA}}$[2] | 0.01381 | 0.01701 | 0.9767 | 0.114434 | 2690 | 3.608 | 3933 | 1.007 | 220.10 | | | |
| $a_{\mathrm{RHA}}$ | | | | | | | | | | 1.660 | 1.750 | 1.627 |
| $b_{\mathrm{RHA}}$ | | | | | | | | | | 12.48 | 22.67 | 23.55 |
| $c_{\mathrm{RHA}}$ | | | | | | | | | | 55.71 | 4.676 | 22.28 |
| $k_{\mathrm{loc,deactivation}}$ | | | | 42.66 | 36.30 | 1.507 | 53.19 | 114.9 | 44.44 | 56.73 | 97.31 | 50.03 |
| $k_{\mathrm{scale,deactivation}}$ | | | | 16.63 | 39.24 | 148.3 | 6.449 | 26.31 | 17.93 | 6.907 | 19.22 | 17.07 |

[1] Gray cells indicate that a parameter was not used in a given model. Parameter units are in **Supplementary Table 2**.
[2]In model D, $k_{\mathrm{RHA}}$ was calculated from $a_{\mathrm{RHA}}$, $b_{\mathrm{RHA}}$, and $c_{\mathrm{RHA}}$ using **Equation S4**.

**Supplementary Table 4 | Equations for model D[1].**

| # | State | Rates | Description |
|---|---|---|---|
| 0 | x_v | - k_degv · x_inputRNA · x_aCas13<br>- k_bds · x_inputRNA · x_u<br>- k_bds · x_inputRNA · x_p1 | Mediated degradation<br>Binding<br>Binding |
| 1 | x_p1 | - k_bds · x_inputRNA · x_p1<br>- k_bds · x_p1 · x_cDNA2 | Binding<br>Binding |
| 2 | x_p2 | - k_bds · x_u · x_p2<br>- k_bds · x_p2 · x_cDNA1 | Binding<br>Binding |
| 3 | x_p1v | + k_bds · x_inputRNA · x_p1 / C_scale<br>- k_degv · x_p1v · x_aCas13<br>- k_RTon · x_p1v · x_RT<br>+ k_RToff · x_RTp1v | Binding<br>Mediated degradation<br>Binding<br>Unbinding |
| 4 | x_p2u | + k_bds · x_u · x_p2<br>- k_degv · x_p2u · x_aCas13<br>- k_RTon · x_p2u · x_RT<br>+ k_RToff · x_RTp2u | Binding<br>Mediated degradation<br>Binding<br>Unbinding |
| 5 | x_p1vdeg | + k_degv · x_p1v · x_aCas13<br>- k_RTon · x_p1vdeg · x_RT<br>+ k_RToff · x_RTp1vdeg | Mediated degradation<br>Binding<br>Unbinding |
| 6 | x_p2udeg | + k_degv · x_p2u · x_aCas13<br>- k_RTon · x_p2udeg · x_RT<br>+ k_RToff · x_RTp2udeg | Mediated degradation<br>Binding<br>Unbinding |
| 7 | x_RT | + k_RToff · x_RTp1v<br>+ k_RToff · x_RTp1vdeg<br>+ k_RToff · x_RTp2cDNA1<br>+ k_RToff · x_RTp2u<br>+ k_RToff · x_RTp2udeg<br>+ k_RToff · x_RTp1cDNA2<br>- k_RTon · x_RT · x_p1v<br>- k_RTon · x_RT · x_p1vdeg<br>- k_RTon · x_RT · x_p2cDNA1<br>- k_RTon · x_RT · x_p2u<br>- k_RTon · x_RT · x_p2udeg<br>- k_RTon · x_RT · x_p1cDNA2<br>+ k_FSS · x_RTp1v<br>+ k_FSS · x_RTp2u<br>+ k_SSS · x_RTp2cDNA1<br>+ k_SSS · x_RTp1cDNA2 | Unbinding<br>Unbinding<br>Unbinding<br>Unbinding<br>Unbinding<br>Unbinding<br>Binding<br>Binding<br>Binding<br>Binding<br>Binding<br>Binding<br>Production/unbinding<br>Degradation/unbinding<br>Degradation/unbinding<br>Production/unbinding |
| 8 | x_RNase | + k_RNaseoff · x_RNasecDNA1v<br>+ k_RNaseoff · x_RNasecDNA2u<br>- k_RNaseon · x_RNase · x_cDNA1v<br>- k_RNaseon · x_RNase · x_cDNA2u<br>+ k_RHA · x_RNasecDNA1v<br>+ k_RHA · x_RNasecDNA2u | Unbinding<br>Unbinding<br>Binding<br>Binding<br>Production/unbinding<br>Production/unbinding |
| 9 | x_RTp1v | - k_RToff · x_RTp1v<br>+ k_RTon · x_RT · x_p1v<br>- k_degv · x_RTp1v · x_aCas13 | Unbinding<br>Binding<br>Degradation |

| | | | |
|---|---|---|---|
| | | − k_FSS · x_RTp1v | Production/unbinding |
| 10 | x_RTp2u | − k_RToff · x_RTp2u<br>+ k_RTon · x_RT · x_p2u<br>− k_degv · x_RTp2u · x_aCas13<br>− k_FSS · x_RTp2u | Unbinding<br>Binding<br>Degradation<br>Production/unbinding |
| 11 | x_RTp1vdeg | − k_RToff · x_RTp1vdeg<br>+ k_RTon · x_RT · x_p1vdeg<br>+ k_degv · x_RTp1v · x_aCas13 | Unbinding<br>Binding<br>Degradation |
| 12 | x_RTp2udeg | − k_RToff · x_RTp2udeg<br>+ k_RTon · x_RT · x_p2udeg<br>+ k_degv · x_RTp2u · x_aCas13 | Unbinding<br>Binding<br>Degradation |
| 13 | x_cDNA1v | + k_FSS · x_RTp1v<br>− k_RNaseon · x_cDNA1v · x_RNase<br>+ k_RNaseoff · x_RNasecDNA1v | Production/unbinding<br>Binding<br>Unbinding |
| 14 | x_cDNA2u | + k_FSS · x_RTp2u<br>− k_RNaseon · x_cDNA2u · x_RNase<br>+ k_RNaseoff · x_RNasecDNA2u | Production/unbinding<br>Binding<br>Unbinding |
| 15 | x_RNasecDNA1v | − k_RHA · x_RNasecDNA1v<br>− k_RNaseoff · x_RNasecDNA1v<br>+ k_RNaseon · x_RNase · x_cDNA1v | Degradation/unbinding<br>Unbinding<br>Binding |
| 16 | x_RNasecDNA2u | − k_RHA · x_RNasecDNA2u<br>− k_RNaseoff · x_RNasecDNA2u<br>+ k_RNaseon · x_RNase · x_cDNA2u | Degradation/unbinding<br>Unbinding<br>Binding |
| 17 | x_cDNA1 | + k_RHA · x_RNasecDNA1v<br>− k_bds · x_cDNA1 · x_p2 | Degradation/unbinding |
| 18 | x_cDNA2 | + k_RHA · x_RNasecDNA2u<br>− k_bds · x_cDNA2 · x_p1 | Degradation/unbinding |
| 19 | x_p2cDNA1 | + k_bds · x_cDNA1 · x_p2<br>+ k_RToff · x_RTp2cDNA1<br>− k_RTon · x_RT · x_p2cDNA1 | Binding<br>Unbinding<br>Binding |
| 20 | x_p1cDNA2 | + k_bds · x_cDNA2 · x_p1<br>+ k_RToff · x_RTp1cDNA2<br>− k_RTon · x_RT · x_p1cDNA2 | Binding<br>Unbinding<br>Binding |
| 21 | x_RTp2cDNA1 | + k_RTon · x_RT · x_p2cDNA1<br>− k_RToff · x_RTp2cDNA1<br>− k_SSS · x_RTp2cDNA1 | Binding<br>Unbinding<br>Production/unbinding |
| 22 | x_RTp1cDNA2 | + k_RTon · x_RT · x_p1cDNA2<br>− k_RToff · x_RTp1cDNA2<br>− k_SSS · x_RTp1cDNA2 | Binding<br>Unbinding<br>Production/unbinding |
| 23 | x_T7 | + k_T7off · x_T7pro<br>− k_T7on · x_T7 · x_pro<br>+ k_txn_eff · x_T7pro | Unbinding<br>Binding<br>Transcription |
| 24 | x_pro | + k_SSS · x_RTp2cDNA1<br>+ k_SSS · x_RTp1cDNA2<br>− k_T7on · x_T7 · x_pro<br>+ k_T7off · x_T7pro | Production/unbinding<br>Production/unbinding<br>Binding<br>Unbinding |

| | | | |
|---|---|---|---|
| | | + k_txn_eff · x_T7pro | Transcription |
| 25 | x_T7pro | - k_T7off · x_T7pro<br>+ k_T7on · x_T7 · x_pro<br>- k_txn_eff · x_T7pro | Unbinding<br>Binding<br>Transcription |
| 26 | x_u | + k_txn_eff · x_T7pro<br>- k_bds · x_u · x_inputRNA / C_scale<br>- k_degv · x_u · x_aCas13<br>- k_cas13 · x_u · x_iCas13<br>- k_bds · x_u · x_p2 | Transcription<br>Binding<br>Mediated degradation<br>Activation<br>Binding |
| 27 | x_iCas13 | - k_cas13 · x_u · x_iCas13 | Activation |
| 28 | x_Cas13^ | + k_cas13 · x_u · x_iCas13 | Activation |
| 29 | x_uv | + k_bds · x_u · x_v / C_scale | Binding |
| 30 | x_qRf | - k_degRrep · x_aCas13 · x_qRf | Mediated degradation |
| 31 | x_q | + k_degRrep · x_aCas13 · x_qRf | Mediated degradation |
| 32 | x_f | + k_degRrep · x_aCas13 · x_qRf | Mediated degradation |

[1] The symbol ^ denotes that target-activated Cas13 encompasses two states: target-activated Cas13a-gRNA (aCas13) and deactivated target-activated Cas13a-gRNA (d_aCas13) ([Cas13] = [aCas13] + [d_aCas13]) (**Supplementary Note 1**). Only aCas13 has indiscriminate ssRNase activity and d_aCas13 is the target-activated Cas13a-gRNA that is removed from the system over time due to deactivation. This type of deactivation is distinct from un-binding of Cas13a-gRNA and the activator RNA, which contributes to the pool of iCas13 (target-inactivated Cas13 a-gRNA).

**Supplementary Table 5 | Comparison of most sensitive parameters across datasets[1].**

| | | $k_{Cas13}$ | $k_{deg\_v}$ | $k_{txn\_base}$ | $k_{FSS}$ | $a_{RHA}$ | $b_{RHA}$ | $c_{RHA}$ | $k_{loc,deactivation}$ | $k_{scale,deactivation}$ |
|---|---|---|---|---|---|---|---|---|---|---|
| Data set 1 | High tolerance | **0.000636** | 239.9 | 858.2 | 0.028 | 1.754 | 10.65 | 42.38 | **61.83** | **8.241** |
| | Low tolerance | **0.000224** | 136.1 | 1151 | 0.096 | 1.66 | 12.48 | 55.71 | **56.73** | **6.907** |
| Data set 2 | High tolerance | **$1.517 * 10^{-5}$** | 12220 | 315.8 | 0.382 | 2.197 | 29.34 | 68.69 | **101.5** | **17.97** |
| | Low tolerance | **$2.230 * 10^{-5}$** | 8940 | 226.3 | 232.9 | 1.75 | 22.67 | 4.676 | **97.31** | **19.22** |
| Data set 3 | High tolerance | **0.000126** | 30450 | 41.57 | 0.079 | 1.071 | 14.71 | 0.627 | **46.40** | **18.54** |
| | Low tolerance | **$6.382 * 10^{-5}$** | 28930 | 119.2 | 0.078 | 1.627 | 23.55 | 22.28 | **50.03** | **17.07** |

[1]Bold indicates that a parameter was one of the most sensitive across all 3 data sets. Red indicates that, for the given data set, the parameter from the low ODE solver tolerance optimization was greater than 1 order of magnitude different from the parameter from the high ODE solver tolerance optimization.

**Supplementary Table 6 | Model fits to each test data set**

| Model (data set used to estimate parameters) | Test data set[1] | $R^2$ | MSE |
|---|---|---|---|
| Data set 1 | Data set 1 training data | 0.99 | 0.0019 |
| | Data set 1 out of sample data | 0.68 | 0.034 |
| | Data set 2 training data | 0.56 | 0.046 |
| | Data set 2 out of sample data | 0.41 | 0.069 |
| | Data set 3 training data | 0.65 | 0.035 |
| | Data set 3 out of sample data | 0.19 | 0.073 |
| Data set 2 | Data set 2 training data | 0.96 | 0.0026 |
| | Data set 2 out of sample data | 0.63 | 0.014 |
| | Data set 1 training data | 0.71 | 0.045 |
| | Data set 1 out of sample data | 0.48 | 0.054 |
| | Data set 3 training data | 0.84 | 0.022 |
| | Data set 3 out of sample data | 0.26 | 0.046 |
| Data set 3 | Data set 3 training data | 0.85 | 0.016 |
| | Data set 3 out of sample data | 0.38 | 0.043 |
| | Data set 1 training data | 0.83 | 0.023 |
| | Data set 1 out of sample data | 0.64 | 0.032 |
| | Data set 2 training data | 0.82 | 0.010 |
| | Data set 2 out of sample data | 0.62 | 0.027 |

[1]The first row for each model is the fit to training data set used to estimate parameters.

**References cited in Supplementary Information**


1. Virtanen, P. *et al.* SciPy 1.0: fundamental algorithms for scientific computing in Python. *Nat. Methods* **17**, 261–272 (2020).
2. Dray, K. E., Muldoon, J. J., Mangan, N. M., Bagheri, N. & Leonard, J. N. GAMES: A Dynamic Model Development Workflow for Rigorous Characterization of Synthetic Genetic Systems. *ACS Synth. Biol.* acssynbio.1c00528 (2022) doi:10.1021/acssynbio.1c00528.
3. Wieland, F.-G., Hauber, A. L., Rosenblatt, M., Tönsing, C. & Timmer, J. On structural and practical identifiability. *Current Opinion in Systems Biology* **25**, 60–69 (2021).
4. Aschenbrenner, S. *et al.* Coupling Cas9 to artificial inhibitory domains enhances CRISPR-Cas9 target specificity. *Sci. Adv.* **6**, eaay0187 (2020).
5. Chen, J. S. *et al.* CRISPR-Cas12a target binding unleashes indiscriminate single-stranded DNase activity. *Science* **360**, 436–439 (2018).
6. Újvári, A. & Martin, C. T. Thermodynamic and Kinetic Measurements of Promoter Binding by T7 RNA Polymerase. *Biochemistry* **35**, 14574–14582 (1996).
7. Reardon, J. E. Human immunodeficiency virus reverse transcriptase: steady-state and pre-steady-state kinetics of nucleotide incorporation. *Biochemistry* **31**, 4473–4479 (1992).
8. Wöhrl, B. M., Krebs, R., Goody, R. S. & Restle, T. Refined model for primer/template binding by HIV-1 reverse transcriptase: pre-steady-state kinetic analyses of primer/template binding and nucleotide incorporation events distinguish between different binding modes depending on the nature of the nucleic acid substrate 1 1Edited by J. Karn. *Journal of Molecular Biology* **292**, 333–344 (1999).
